# Mammals, birds and reptiles evolved with signature proportions of numbers of neurons across their brain structures

**DOI:** 10.1101/2022.06.20.496835

**Authors:** Suzana Herculano-Houzel

## Abstract

Modern mammals, birds, and “reptiles” (squamates, turtles and crocodiles) share their developmental and evolutionary origins in the ancestral, stem amniotes of 300 million years ago. This study explores how the brains of amniotes diverged in their neuronal composition. The systematic analysis of a large dataset on the cellular composition of the major parts of the brain of 242 amniote species shows that it is not the case that a general theme in amniote evolution is the proportional expansion of the telencephalon or pallium, for such expansion only happened in owls, in primates, and in the largest mammalian species. Instead, the brains of extant mammalian, avian, and reptile species are characterized by signature proportions of numbers of neurons across the brain divisions. Reptile brains, with few exceptions, are characterized by having fewer than 10 million neurons in each brain division, with about 1.5 neurons in the telencephalon and 0.5 neuron in the cerebellum to every neuron in the rest of brain. In contrast, the brains of the closely related birds are characterized by much larger numbers of neurons that occur at a higher, fixed proportion of 4.5 neurons in the cerebellum to every neuron in the rest of brain, with variable ratios of pallial neurons. The brains of mammalian species, in turn, are characterized by larger numbers of neurons that occur at an average 4 neurons in the cerebellum to every neuron in the pallium regardless of numbers of neurons in the rest of brain. There is a striking continuity in the scaling of the pallium (or telencephalon) of extant mammalian and squamate brains that argues for a quantitative continuum between the two groups and dispels the mistaken notion that mammalian brains evolved with a qualitative change or an “addition” of structures to the “reptilian brain”. The shared scaling rules between mammalian and squamate brain structures also allow for predicting the composition of early synapsid brains in amniote evolution.

## Introduction

Extant mammals (synapsids) and sauropsids (birds, squamates, turtles and crocodiles), which together constitute the amniotes amongst animals, are markedly different in appearance, body size, and life histories. Yet, they derive from common stem amniote ancestors that lived ca. 300 million years ago (Kemp, 2006), and their brains still share a common developmental program (Puelles et al., 2013). The largest amniote brains are all mammalian, but mammalian brains can be as small as those of some lizards; modern bird brains, in turn, are larger than most squamate brains, but not larger than those of dog-sized mammals (Kverkova et al., 2022). How did so much diversity in brain size arise, how come the different ranges, and how do they compare in terms of numbers of neurons?

Understanding the evolutionary history of modern amniote brains – those of mammals, birds, squamates, turtles and crocodiles – is an enterprise that is necessarily built on the comparative study of these extant brains in order to infer how they arose from evolutionary changes over time in their respective developmental programs. Until the advent of the isotropic fractionator, a method to reliably and rapidly count the neuronal and non-neuronal cells that compose brains without having to painstakingly cut them into thin sections for histological processing, microscopy and stereological analysis (Herculano-Houzel and Lent, 2005; Herculano-Houzel et al., 2015a), comparative studies of vertebrate brains were necessarily focused on the available data on structure size. While any data were better than no data, the limitation to data on structure size posed a number of problems. First, the interpretation of comparative studies of brain structure volume required the tacit assumption that the underlying scaling rules governing the cellular composition of brain tissues were shared across species and clades, or any results would be meaningless. Data obtained during the early 2000s using the isotropic fractionator rapidly proved this assumption to be invalid, for the relationship between the size of a brain or brain structure and its number of neurons is clade-specific such that structures of a similar size can have numbers of neurons that vary by a factor of up to 10-fold across species of different clades (Herculano-Houzel et al., 2006, 2007). Of note, the human cerebral cortex would indeed appear to be an outlier in its number of neurons, with ten times the number of neurons expected for a rodent cerebral cortex of similar mass – but its average of 16 billion neurons matches the prediction for a generic primate cortex of its mass (Azevedo et al., 2009). Second, absolute and relative measurements of brain structure size often yielded conflicting interpretations (for instance, see Clark et al., 2001 and Sultan, 2002), for their meaning depends additionally on whether or not the size of the whole brain is a relevant reference and normalizer for brain structures. This, in turn, depends on whether the relative size of a brain structure accurately reflects the relative number of brain neurons located in that structure – which, it turns out, it does not (Herculano-Houzel, 2010; Herculano-Houzel et al., 2014a). For instance, the number of neurons in the mammalian cerebellum closely accompanies the number of neurons in the cerebral cortex across species such that an average rate of four neurons in the former for every neuron in the latter applies to the majority of mammals, even though the relative size of the cerebellum becomes smaller as the relative size of the cerebral cortex increases in larger species (Herculano-Houzel, 2010).

An even more problematic complication is the use of normalization of brain or brain structure size to body mass for comparisons across species, an expedient that became popular once it finally succeeded in setting humans apart from other species (Jerison, 1973). The problem with using body mass as a normalizer is that it assumes a mandatory and universal relationship between body mass and the numbers of neurons, and the mass of the brain structures housing those neurons, that operate the body (Jerison, 1973). In contrast, we have observed repeatedly that body size is largely free to vary across individuals and species without mandatory consequences for the numbers of neurons that compose brain structures or spinal cord (Burish et al., 2010; Herculano-Houzel et al., 2015b, 2015c). Put another way, bodies of a similar size occur with up to 50-fold different numbers of neurons in the spinal cord and brainstem across mammalian species. To understand evolutionary changes in brain composition, then, one must look at exactly that: variations in the numbers of neurons that compose the brain, regardless of variations in body mass.

Using the isotropic fractionator, my colleagues and I generated a body of data on the numbers of neuronal and non-neuronal cells that compose the major brain structures of 75 mammalian, 28 bird species, and the Nile crocodile (Herculano-Houzel et al., 2015b; Ngwenya et al., 2016; Olkowicz et al., 2016; Dos Santos et al., 2017; Jardim-Messeder et al., 2017). Those data showed that while similar scaling relationships for each of the major brain parts applied to many mammalian (with the exception of primates) and bird species, there were obviously higher neuronal densities in the telencephalon of songbirds and parrots – even higher than in primates (Olkowicz et al., 2016). While those findings provided an answer to the question of how some birds can be as cognitively capable as primates (answer: because their small brains harbor primate-like numbers of pallial, and particularly associative pallial, neurons; Ströckens et al., 2022, Sol et al., 2022), addressing the question of how bird brain composition came to differ from mammalian brain composition required obtaining data on the third group of amniotes that links them together: extant ectothermic sauropsids. That is because while birds are a derived group amongst sauropsids, appearing within theropod dinosaurs during the Jurassic (Norell and Xu, 2005), modern squamates, which are ectothermic Sauropsids, are a sister group to mammals, with whom they share common ancestors that lived ca. 300 Mya (Kemp, 2006). Thus, comparison of the neuronal scaling rules between squamates and mammals might reveal the scaling rules that applied to the brains of ancestral amniotes.

That required dataset, with the numbers of neurons that compose the brains of 107 ectothermic sauropsid species in 6 major clades and an additional 37 endothermic sauropsid species covering a total of 9 avian (hereafter “bird”) clades, was published recently (Kverkova et al., 2022). In this dataset, sauropsid brains were divided into telencephalon/cerebellum/rest of brain, in which case the subpallium is included in the telencephalon, and “rest of brain” is the brainstem plus diencephalon. In birds, numbers of cells were also available for the two parts of the telencephalon: the pallium (directly equivalent and indeed homologous to the mammalian cerebral cortex; Puelles et al., 2013) and subpallium (Olkowicz et al., 2016; Kverkova et al., 2022). By moving numbers for the subpallium into the “rest of brain”, the bird data can be compared directly with the published data for mammals, in which the brain is subdivided into cortex/cerebellum/rest of brain, and the latter also includes the subpallium. While it is unfortunate that the subpallium was not separated from the pallium in the telencephalon of the ectothermic sauropsids examined, the total number of neurons in their telencephalon is small compared to the “rest of brain” in these species (Kverkova et al., 2022). With these caveats in mind, the combined dataset allows for the comparison of numbers of cells in the ectothermic sauropsid telencephalon/bird pallium/mammalian cerebral cortex, cerebellum, and the “rest of brain” (hindbrain to diencephalon in ectothermic sauropsids, hindbrain to subpallium in birds and mammals). For the sake of simplicity, and absent a better word to describe the ensemble of ectothermic Sauropsids (that is, the ensemble of Archosauria that includes squamates, turtles and crocodiles but excludes birds), I will heretofore refer to the ectothermic Sauropsids as “reptiles”.

The study that originally reported the reptile and extended bird dataset (Kverkova et al., 2022) focused on describing the grade shifts in the relationship between brain and body mass that characterize amniote evolution, but did not address a number of fundamental issues in amniote brain evolution, starting with the neuronal scaling rules that apply to each clade specifically whose systematic comparison is necessary to shed light on the evolution of the amniote brain. Further, anchoring interpretations of brain evolution on relationships to body mass is a flawed expedient that assumes a mandatory and universal relationship between brain and body mass that is not there, as attested recently by studies on bird and mammalian brain evolution (Ksepka et al., 2020; Bertrand et al., 2022) and our demonstrations that there is no mandatory, much less a universal, scaling relationship between numbers of brain neurons and body mass across species (Herculano-Houzel et al., 2015b,c; Ngwenya et al., 2016). Additionally, studies of brain scaling often resort to “correcting” for phylogenetic relationships in the dataset (as done in Kverkova et al., 2022), a procedure that aspires to eliminate sampling bias but glosses over differences in scaling relationships across clades, and contaminates scaling relationships in the raw data with inferences about phylogenetic relations that may or may not be correct. Since the point of the present analysis is understanding the mathematics of the relationship between the size of the brain and of how many cells brain tissue is made across different animal clades, I focus the analysis on the scaling relationships that apply to the raw data; the only phylogenetic information taken into consideration is that pertaining to the nested clades that each animal belongs to. Indeed, the expedient of analyzing scaling relationships separately per clade achieves on its own the most important type of “correcting” for phylogenetic relationships, which is exactly the possibility that clade-specific scaling relationships exist.

Here I use the combined dataset of 242 amniote species to determine, regardless of body mass, the cellular scaling relationships that apply to each clade among the three groups of amniote animals – reptiles, mammals, and birds – to then address the following questions. First, is the relationship between brain structure mass and number of non-neuronal cells (the so-called non-neuronal scaling rule) that applies universally to mammals (Herculano-Houzel and Dos Santos, 2018) extensive to reptiles and birds? Non-neuronal cells in the brain include the endothelial cells that compose the brain vasculature, which may or may not scale similarly across endothermic and ectothermic species. Is the scaling of non-neuronal cells shared at least by all sauropsids (ectothermic reptiles and endothermic birds), or by all endothermic species? Second, are the main brain structures of amniotes fundamentally different in their neuronal composition, or do they form a continuum along a single scaling relationship across reptiles, birds and mammals, at least across the more closely related clades of reptiles and birds? Third, if there are such continuums, can the scaling rules that applied to the main divisions of the brain of ancestral amniote species be inferred? In that case, can amniote brain diversity be understood as a simple evolutionary story of continuity with occasional deviations, without major grade shifts, or are there major grade shifts to be accounted for – for example, associated with the independent evolution of endothermy in birds and mammals (Nespolo et al., 2011)?

The findings below are based exclusively on directly comparable data on numbers of neurons and mass of the telencephalon, cerebellum and rest of brain obtained with the isotropic fractionator for 242 species of reptiles, mammals and birds, compiled in Kverkova et al. (2022). For consistency, the olfactory bulbs, which are often not collected whole in brain dissections, are not included in any structure. All power functions were calculated using least-squares regression of log-transformed values in JMP 16 (Carey, NC).

### Relationships of brain mass to body mass

Any systematic examination of how the neuronal composition of the brain evolved amongst amniotes must begin with laying down the basic aspects of how those brains have changed, and that starts with the relationship between the size of the brain and the size of the body. Across the 242 species of reptiles, mammals, and birds in the combined dataset, whole brain mass varies almost 500,000-fold, from 0.01 g in the Algerian sand gecko to 4,618.62 g in the African elephant, whereas body mass varies ten times as much, by a factor of 5 million, from 1 g to 5 Tons in the same species (Figure 1A). As a whole, larger species have larger brains, but in a clade-specific manner. The power functions that describe the scaling of body mass with brain mass across species of each clade amongst the three groups of amniotes in the dataset are provided in Table 1. Importantly, within each group – reptiles, birds, and mammals –, a number of clades share scaling rules, which is key to inferring what was once the scaling rule that applied to ancestral animals in each group. The power functions that apply for each clade, and to groups of clades that share scaling rules, are documented in Table 1.

**Figure 1.**
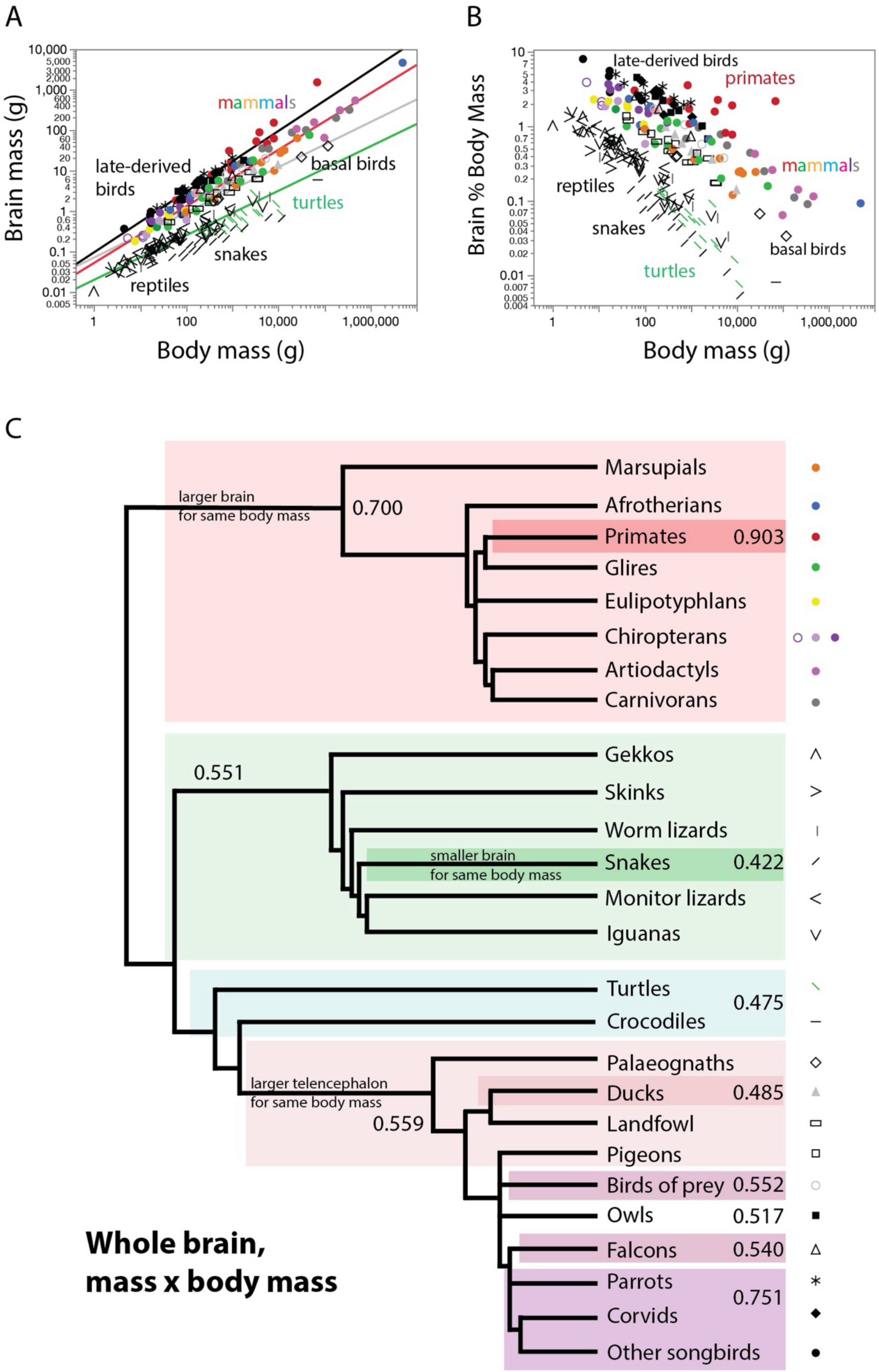
Larger amniotes have larger brains, but relatively smaller brains, in a clade-specific manner. **A**, absolute brain mass increases as different power functions of body mass across reptiles (minus snakes, turtles and crocodiles; plotted in green), basal birds (plotted in grey), late-derived birds (plotted in black), and mammals (minus primates; plotted in red). **B**, relative brain mass decreases with increasing body mass. For the sake of clarity, power functions are not plotted here. **C**, evolutionary tree showing the clades that share similar scaling exponents, which are shown where applicable. Power functions are listed in Table 1. Each data point corresponds to one species. The key used to depict each clade, employed throughout this chapter, is shown to the right.

**Table 1.** Scaling of mass of the different brain structures of reptiles, birds and mammals with varying body mass. Values correspond to the equation Mstr = e^a±SE^ Mbd ^b±SE^, the p-values for a and b, and the r^2^ for the power fit, where Mstr is the mass of a given brain structure and Mbd is the average body mass for the species. * pig excluded. **capybara excluded.

| $M_{str} \times M_{bd}$ | n | a in $e^a$ | b in $M_{bd}^b$ | p, a | p, b | $r^2$ |
| --- | --- | --- | --- | --- | --- | --- |
| **Whole brain** |  |  |  |  |  |  |
| **Reptiles** |  |  |  |  |  |  |
| All | 108 | -3.749±0.104 | **0.460**±0.020 | <0.0001 | <0.0001 | 0.842 |
| Anguimorpha | 7 | -4.643±0.376 | **0.646**±0.057 | <0.0001 | <0.0001 | 0.962 |
| Crocodylia | 1 |  |  |  |  |  |
| Gekkota | 14 | -4.294±0.172 | **0.695**±0.058 | <0.0001 | <0.0001 | 0.923 |
| Iguania | 20 | -3.787±0.162 | **0.505**±0.032 | <0.0001 | <0.0001 | 0.932 |
| Lacertoidea | 15 | -3.847±0.176 | **0.561**±0.045 | <0.0001 | <0.0001 | 0.924 |
| Scincoidea | 14 | -4.178±0.231 | **0.598**±0.061 | <0.0001 | <0.0001 | 0.891 |
| Serpentes | 18 | -4.148±0.259 | **0.422**±0.043 | <0.0001 | <0.0001 | 0.861 |
| Testudines | 19 | -3.528±0.512 | **0.422**±0.072 | <0.0001 | <0.0001 | 0.668 |
| Testudines and Crocodylia | 20 | -3.877±0.420 | **0.475**±0.057 | <0.0001 | <0.0001 | 0.793 |
| All minus Serpentes, Testudines and Crocodylia | 70 | -3.946±0.083 | **0.551**±0.019 | <0.0001 | <0.0001 | 0.927 |
| **Birds** |  |  |  |  |  |  |
| All | 65 | -1.424±0.241 | **0.470**±0.039 | <0.0001 | <0.0001 | 0.701 |
| Accipitriformes | 4 | -1.748±0.340 | **0.552**±0.048 | 0.0359 | 0.0076 | 0.985 |
| Anseriformes | 7 | -1.731±0.283 | **0.485**±0.039 | 0.0017 | <0.0001 | 0.968 |
| Columbiformes | 5 | -3.189±0.142 | **0.681**±0.025 | 0.0002 | 0.0001 | 0.996 |
| Falconiformes | 3 | -1.710±0.290 | **0.540**±0.052 | 0.1070 | 0.0611 | 0.991 |
| Galliformes | 9 | -3.023±0.285 | **0.612**±0.044 | <0.0001 | <0.0001 | 0.966 |
| Palaeognathae | 6 | -2.814±0.080 | **0.562**±0.009 | <0.0001 | <0.0001 | 0.999 |
| Passeriformes | 13 | -2.269±0.239 | **0.714**±0.051 | <0.0001 | <0.0001 | 0.946 |
| Psittaciformes | 11 | -2.488±0.254 | **0.785**±0.048 | <0.0001 | <0.0001 | 0.968 |
| Strigiformes | 7 | -1.155±0.256 | **0.517±0.042** | 0.0063 | <0.0001 | 0.969 |
| Passeriformes,<br>Psittaciformes | 24 | -2.376±0.174 | **0.751±0.035** | <0.0001 | <0.0001 | 0.954 |
| Galliformes,<br>Columbiformes,<br>Palaeognathae | 20 | -2.674±0.146 | **0.559±0.021** | <0.0001 | <0.0001 | 0.976 |
| Accipitriformes,<br>Falconiformes | 7 | -1.771±0.159 | **0.554±0.025** | 0.0001 | <0.0001 | 0.990 |
| **Mammals** |  |  |  |  |  |  |
| All | 69 | -2.983±0.183 | **0.729±0.024** | <0.0001 | <0.0001 | 0.931 |
| All non-primates | 57 | -2.982±0.127 | **0.700±0.017** | <0.0001 | <0.0001 | 0.969 |
| Afrotheria | 6 | -2.927±0.270 | **0.740±0.033** | 0.0004 | <0.0001 | 0.992 |
| Artiodactyla* | 4 | -0.939±0.445 | **0.548±0.038** | 0.1690 | 0.0048 | 0.990 |
| Carnivora | 8 | -1.877±0.499 | **0.609±0.051** | 0.0094 | <0.0001 | 0.959 |
| Chiroptera | 13 | -2.856±0.340 | **0.693±0.092** | <0.0001 | <0.0001 | 0.838 |
| Eulipotyphla | 5 | -3.112±0.329 | **0.727±0.094** | 0.0025 | 0.0045 | 0.952 |
| Marsupialia | 9 | -3.680±0.510 | **0.742±0.061** | 0.0002 | <0.0001 | 0.955 |
| Primata | 11 | -3.323±0.645 | **0.903±0.082** | 0.0006 | <0.0001 | 0.931 |
| Glires | 10 | -3.287±0.489 | **0.712±0.071** | 0.0001 | <0.0001 | 0.927 |
| Scandentia | 1 |  |  |  |  |  |
| **Cortex/pallium/telencephalon** |  |  |  |  |  |  |
| **Reptiles** |  |  |  |  |  |  |
| All | 108 | -4.586±0.110 | **0.503±0.021** | <0.0001 | <0.0001 | 0.850 |
| Anguimorpha | 7 | -5.231±0.514 | **0.651±0.078** | 0.0002 | 0.0004 | 0.933 |
| Crocodylia | 1 |  |  |  |  |  |
| Gekkota | 14 | -4.851±0.219 | **0.690±0.074** | <0.0001 | <0.0001 | 0.880 |
| Iguania | 20 | -4.793±0.165 | **0.527±0.033** | <0.0001 | <0.0001 | 0.934 |
| Lacertoidea | 15 | -4.742±0.184 | **0.616±0.046** | <0.0001 | <0.0001 | 0.931 |
| Scincoidea | 14 | -4.959±0.223 | **0.640±0.058** | <0.0001 | <0.0001 | 0.909 |
| Serpentes | 18 | -5.326±0.309 | **0.519±0.051** | <0.0001 | <0.0001 | 0.867 |
| Testudines | 19 | -3.845±0.585 | **0.415±0.083** | <0.0001 | 0.0001 | 0.598 |
| Testudines and<br>Crocodylia | 20 | -4.179±0.474 | **0.465±0.065** | <0.0001 | <0.0001 | 0.742 |
| All minus<br>Serpentes | 90 | -4.593±0.092 | **0.532±0.018** | <0.0001 | <0.0001 | 0.912 |
| **Birds** |  |  |  |  |  |  |
| All | 65 | -2.029±0.299 | **0.481±0.048** | <0.0001 | <0.0001 | 0.614 |
| Accipitriformes | 4 | -2.979±0.334 | **0.646±0.047** | 0.0124 | 0.0054 | 0.989 |
| Anseriformes | 7 | -2.316±0.370 | **0.494±0.052** | 0.0015 | 0.0002 | 0.948 |
| Columbiformes | 5 | -4.106±0.162 | **0.708±0.029** | 0.0001 | 0.0001 | 0.995 |
| Falconiformes | 3 | -2.449±0.282 | **0.564±0.050** | 0.0729 | 0.0568 | 0.992 |
| Galliformes | 9 | -3.997±0.334 | **0.647±0.051** | <0.0001 | <0.0001 | 0.958 |
| Palaeognathae | 6 | -3.841±0.086 | **0.611±0.010** | <0.0001 | <0.0001 | 0.999 |
| Passeriformes | 13 | -2.999±0.276 | **0.771±0.059** | <0.0001 | <0.0001 | 0.939 |
| Psittaciformes | 11 | -3.118±0.277 | **0.823±0.052** | <0.0001 | <0.0001 | 0.965 |
| Strigiformes | 7 | -1.573±0.315 | **0.532±0.051** | 0.0041 | 0.0001 | 0.956 |
| Passeriformes,<br>Psittaciformes,<br>Strigiformes | 31 | -2.837±0.184 | **0.749±0.035** | <0.0001 | <0.0001 | 0.940 |
| Galliformes,<br>Columbiformes,<br>Palaeognathae | 20 | -3.697±0.151 | **0.605±0.021** | <0.0001 | <0.0001 | 0.978 |
| Accipitriformes,<br>Falconiformes | 7 | -2.708±0.184 | **0.608±0.028** | <0.0001 | <0.0001 | 0.989 |
| **Mammals** |  |  |  |  |  |  |
| All | 70 | -3.837±0.220 | **0.774±0.029** | <0.0001 | <0.0001 | 0.912 |
| All non-primates | 57 | -3.878±0.150 | **0.743±0.020** | <0.0001 | <0.0001 | 0.962 |
| Afrotheria | 6 | -3.670±0.264 | **0.761±0.032** | 0.0002 | <0.0001 | 0.993 |
| Artiodactyla* | 4 | -1.759±0.334 | **0.589±0.029** | 0.0341 | 0.0023 | 0.995 |
| Carnivora | 8 | -2.475±0.610 | **0.631±0.062** | 0.0067 | <0.0001 | 0.945 |
| Chiroptera | 13 | -3.856±0.454 | **0.761±0.123** | <0.0001 | <0.0001 | 0.777 |
| Eulipotyphla | 5 | -3.758±0.476 | **0.725±0.137** | 0.0042 | 0.0131 | 0.904 |
| Marsupialia | 9 | -4.728±0.584 | **0.797±0.070** | <0.0001 | <0.0001 | 0.949 |
| Primata | 12 | -3.901±0.652 | **0.942±0.084** | 0.0001 | <0.0001 | 0.926 |
| Glires | 10 | -4.319±0.530 | **0.760±0.077** | <0.0001 | <0.0001 | 0.925 |
| Scandentia | 1 |  |  |  |  |  |
| **Cerebellum** |  |  |  |  |  |  |
| **Reptiles** |  |  |  |  |  |  |
| All | 108 | -7.731±0.213 | **0.528±0.040** | <0.0001 | <0.0001 | 0.625 |
| Anguimorpha | 7 | -11.226±1.110 | **1.081±0.169** | 0.0002 | 0.0014 | 0.891 |
| Crocodylia | 1 |  |  |  |  |  |
| Gekkota | 14 | -8.310±0.234 | **0.814±0.079** | <0.0001 | <0.0001 | 0.900 |
| Iguania | 20 | -6.926±0.196 | **0.498±0.039** | <0.0001 | <0.0001 | 0.901 |
| Lacertoidea | 15 | -7.614±0.300 | **0.568±0.076** | <0.0001 | <0.0001 | 0.813 |
| Scincoidea | 14 | -8.453±0.548 | **0.673±0.144** | <0.0001 | 0.0005 | 0.646 |
| Serpentes | 18 | -8.274±0.593 | **0.387±0.097** | <0.0001 | 0.0011 | 0.496 |
| Testudines | 19 | -6.785±0.748 | **0.443±0.105** | <0.0001 | 0.0008 | 0.510 |
| Testudines and Crocodilia | 20 | -7.408±0.624 | **0.537±0.085** | <0.0001 | <0.0001 | 0.690 |
| All minus Serpentes | 90 | -7.814±0.159 | **0.604±0.030** | <0.0001 | <0.0001 | 0.818 |
| All minus Iguania, Serpentes | 70 | -8.033±0.180 | **0.627±0.034** | <0.0001 | <0.0001 | 0.833 |
| **Birds** |  |  |  |  |  |  |
| All | 65 | -3.856±0.161 | **0.507±0.026** | <0.0001 | <0.0001 | 0.859 |
| Accipitriformes | 4 | -3.014±0.155 | **0.453±0.022** | 0.0027 | 0.0024 | 0.995 |
| Anseriformes | 7 | -4.263±0.328 | **0.548±0.046** | <0.0001 | <0.0001 | 0.966 |
| Columbiformes | 5 | -5.264±0.274 | **0.709±0.048** | 0.0003 | 0.0007 | 0.986 |
| Falconiformes | 3 | -4.104±0.321 | **0.625±0.057** | 0.0496 | 0.0583 | 0.992 |
| Galliformes | 9 | -5.033±0.271 | **0.614±0.041** | <0.0001 | <0.0001 | 0.969 |
| Palaeognathae | 6 | -4.890±0.122 | **0.577±0.014** | <0.0001 | <0.0001 | 0.998 |
| Passeriformes | 13 | -4.327±0.163 | **0.646±0.035** | <0.0001 | <0.0001 | 0.969 |
| Psittaciformes | 11 | -4.410±0.234 | **0.667±0.044** | <0.0001 | <0.0001 | 0.962 |
| Strigiformes | 7 | -3.530±0.155 | **0.490±0.025** | <0.0001 | <0.0001 | 0.987 |
| Passeriformes, Psittaciformes, Strigiformes | 31 | -4.184±0.119 | **0.613±0.023** | <0.0001 | <0.0001 | 0.962 |
| Anseriformes, Galliformes, Columbiformes, Palaeognathae | 27 | -4.639±0.169 | **0.570±0.024** | <0.0001 | <0.0001 | 0.959 |
| Accipitriformes, Falconiformes | 7 | -3.546±0.237 | **0.527±0.037** | <0.0001 | <0.0001 | 0.976 |
| **Mammals** |  |  |  |  |  |  |
| All | 72 | -4.803±0.168 | **0.704±0.022** | <0.0001 | <0.0001 | 0.937 |
| All non-primates | 58 | -4.772±0.131 | **0.676±0.017** | <0.0001 | <0.0001 | 0.964 |
| Afrotheria | 6 | -5.274±0.398 | **0.798±0.048** | 0.0002 | <0.0001 | 0.986 |
| Artiodactyla* | 4 | -3.940±1.228 | **0.612±0.105** | 0.0850 | 0.0282 | 0.944 |
| Carnivora | 8 | -3.936±0.407 | **0.606±0.042** | <0.0001 | <0.0001 | 0.972 |
| Chiroptera | 13 | -4.226±0.225 | **0.578±0.061** | <0.0001 | <0.0001 | 0.891 |
| Eulipotyphla | 5 | -5.661±0.494 | **0.887±0.142** | 0.0014 | 0.0083 | 0.929 |
| Marsupialia | 10 | -5.124±0.425 | **0.684±0.050** | <0.0001 | <0.0001 | 0.959 |
| Primata | 13 | -4.321±0.631 | **0.739±0.074** | <0.0001 | <0.0001 | 0.900 |
| Glires** | 9 | -5.370±0.691 | **0.743±0.109** | 0.0001 | 0.0002 | 0.869 |
| Scandentia | 1 |  |  |  |  |  |
| **Rest of brain** |  |  |  |  |  |  |
| **Reptiles** |  |  |  |  |  |  |
| All | 108 | -4.231±0.111 | **0.411±0.021** | <0.0001 | <0.0001 | 0.789 |
| Anguimorpha | 7 | -5.384±0.422 | **0.644±0.064** | <0.0001 | 0.0002 | 0.953 |
| Crocodylia | 1 |  |  |  |  |  |
| Gekkota | 14 | -4.847±0.141 | **0.640±0.047** | <0.0001 | <0.0001 | 0.938 |
| Iguania | 20 | -4.319±0.176 | **0.496±0.035** | <0.0001 | <0.0001 | 0.918 |
| Lacertoidea | 15 | -4.338±0.166 | **0.533±0.042** | <0.0001 | <0.0001 | 0.925 |
| Scincoidea | 14 | -4.676±0.257 | **0.564±0.067** | <0.0001 | <0.0001 | 0.853 |
| Serpentes | 18 | -4.521±0.294 | **0.370±0.048** | <0.0001 | <0.0001 | 0.786 |
| Testudines | 19 | -4.751±0.427 | **0.445±0.060** | <0.0001 | <0.0001 | 0.762 |
| Testudines and Crocodylia | 20 | -5.134±0.359 | **0.502±0.049** | <0.0001 | <0.0001 | 0.854 |
| All minus Testudines, Crocodylia, Serpentes | 70 | -4.514±0.085 | **0.535±0.019** | <0.0001 | <0.0001 | 0.919 |
| **Birds** |  |  |  |  |  |  |
| All | 65 | -2.563±0.174 | **0.457±0.028** | <0.0001 | <0.0001 | 0.810 |
| Accipitriformes | 4 | -2.114±0.490 | **0.428±0.069** | 0.0497 | 0.0254 | 0.950 |
| Anseriformes | 7 | -2.650±0.187 | **0.436±0.026** | <0.0001 | <0.0001 | 0.982 |
| Columbiformes | 5 | -3.852±0.163 | **0.630±0.029** | 0.0002 | 0.0002 | 0.994 |
| Falconiformes | 3 | -2.431±0.297 | **0.457±0.053** | 0.0773 | 0.0737 | 0.987 |
| Galliformes | 9 | -3.707±0.228 | **0.570±0.035** | <0.0001 | <0.0001 | 0.974 |
| Palaeognathae | 6 | -3.543±0.753 | **0.530±0.086** | 0.0093 | 0.0035 | 0.906 |
| Passeriformes | 13 | -3.110±0.203 | **0.614±0.044** | <0.0001 | <0.0001 | 0.948 |
| Psittaciformes | 11 | -3.508±0.229 | **0.728±0.043** | <0.0001 | <0.0001 | 0.969 |
| Strigiformes | 7 | -2.498±0.152 | **0.470±0.025** | <0.0001 | <0.0001 | 0.986 |
| Passeriformes, Psittaciformes, Strigiformes | 31 | -3.003±0.176 | **0.595±0.033** | <0.0001 | <0.0001 | 0.916 |
| Anseriformes, Galliformes, Columbiformes, Palaeognathae | 27 | -3.343±0.185 | **0.520±0.026** | <0.0001 | <0.0001 | 0.941 |
| Accipitriformes, Falconiformes | 7 | -2.408±0.244 | **0.464±0.038** | 0.0002 | <0.0001 | 0.968 |
| **Mammals** |  |  |  |  |  |  |
| All | 70 | -3.683±0.125 | **0.635±0.017** | <0.0001 | <0.0001 | 0.956 |
| All non-primates | 57 | -3.697±0.105 | **0.622±0.014** | <0.0001 | <0.0001 | 0.973 |
| Afrotheria | 6 | -3.508±0.277 | **0.641±0.034** | 0.0002 | <0.0001 | 0.989 |
| Artiodactyla* | 4 | -0.636±0.657 | **0.378±0.056** | 0.4352 | 0.0215 | 0.958 |
| Carnivora | 8 | -2.914±0.341 | **0.540±0.035** | 0.0001 | <0.0001 | 0.976 |
| Chiroptera | 13 | -3.819±0.287 | **0.668±0.078** | <0.0001 | <0.0001 | 0.871 |
| Eulipotyphla | 5 | -4.010±0.119 | **0.673±0.034** | <0.0001 | 0.0003 | 0.992 |
| Marsupialia | 9 | -4.376±0.457 | **0.679±0.055** | <0.0001 | <0.0001 | 0.957 |
| Primata | 12 | -3.726±0.591 | **0.706±0.076** | <0.0001 | <0.0001 | 0.896 |
| Glires** | 9 | -4.130±0.615 | **0.689±0.097** | 0.0003 | 0.0002 | 0.878 |
| Scandentia | 1 |  |  |  |  |  |

The wealth of data in the combined dataset can be summarized as follows. First: In every clade, brain mass scales with brain mass raised to an exponent that is smaller than unity. This means that while larger animals have larger brains, the brain does not increase in mass as rapidly as the body does. The consequence is that larger animals have *relatively smaller* brains, as shown in Figure 1B. The range of variation is enormous: amniote evolution has resulted in brains that represent as little as 0.005% (in the snake *Boa constrictor*) and as much as 8% of body mass (in the songbird *Regulus regulus*).

Second: As has been well established, and as Figures 1A and B illustrate, reptiles (the ectothermic Sauropsids) have systematically smaller brains for their body sizes compared to birds and mammals (Jerison, 1973), such that all data points for reptile species fall well below the data points for the endothermic species (Figure 1A,B). The advent of endothermy was thus accompanied by a step increase in the relative size of the brain (Jerison, 1973). Importantly, however, similar step-like changes apply amongst birds and mammals. Amongst the former, “basal birds” – Palaeognaths, Galliformes and Columbiformes, in particular – have much smaller brains than “late-derived birds” – Passeriformes and Psittaciformes – of similar body mass. Amongst mammals, it is primates that have larger brains than other species of similar body mass (Figure 1B).

Third: As mentioned above, there is variation across clades within each of the three groups of amniotes in the scaling of brain mass with body mass. The schematic in Figure 1C captures the main differences. Most reptiles share the scaling of brain mass with body mass raised to an exponent of 0.551±0.019, but snakes, and turtles and crocodiles, stand out with not only relatively smaller brains but also smaller scaling exponents (Table 1). Interestingly, the scaling exponent that applies to basal birds, of 0.559±0.021, is not significantly different from the exponent that applies to most reptiles – although, as mentioned above, basal birds have brains that are roughly 3.6x as large as reptiles of a similar body mass. The similar scaling exponent with a shifted intercept (Table 1) indicates that birds arose with a step change towards a relatively larger brain size – which, mechanistically, could result both from the enlargement of the brain for a given body mass, and from the shrinking of the body that forms around the brain. Mammals, in contrast, show both the step increase in relative brain size and a significantly different scaling exponent for brain mass with body mass, of 0.700±0.017 across non-primate species (Figure 1C, Table 1).

These major differences in the scaling of brain mass with body mass within and across reptiles, birds and mammals, depicted schematically in Figure 1C, show that a change in relative brain size did occur twice independently, first at the origin of mammals (assuming that ancestral amniotes were ectothermic, as modern reptiles remain; see Nespolo et al., 2011), and later again at the origin of birds. However, similar step changes apparently happened anew at the origin of Strigiformes; at the origin of Passeriformes and Psittaciformes; and at the origin of Primates. This indicates that there are mechanisms other than whatever caused endothermy that may lead to a relative enlargement of the amniote brain.

The three fundamental observations regarding the scaling of brain mass with body mass in amniotes – namely, the scaling exponents smaller than unity, the step change with the advent of endothermy, and the clade-specific scaling – also apply to each of the three main subdivisions of the brain: the telencephalon/pallium (which also describes the mammalian cerebral cortex), the cerebellum, and the “rest of brain”. Figure 2 shows the scaling of the reptilian telencephalon, and the pallium of birds and mammals, where it is apparent that larger animals have an absolutely larger but also relatively smaller pallium; that birds and mammals have a larger pallium than reptiles of a similar body mass; and that clade-specific scaling rules apply. Most reptiles share the increase in mass of the telencephalon with body mass raised to the power of 0.532±0.018; interestingly, amongst reptiles of a given body mass, snakes have a relatively smaller telencephalon (expressed as a percentage of body mass, not of brain mass), although the scaling exponent remains similar (0.519±0.051). In both mammals and basal birds, there is a shift towards a relatively larger pallium compared to the body – which, again, could be the result of enlargement of the pallium or of a reduction in the size of the body –, and both within mammals and birds, there are some clades that have a particularly enlarged pallium for a given body mass (namely, primates, songbirds, parrots, and owls).

**Figure 2.**
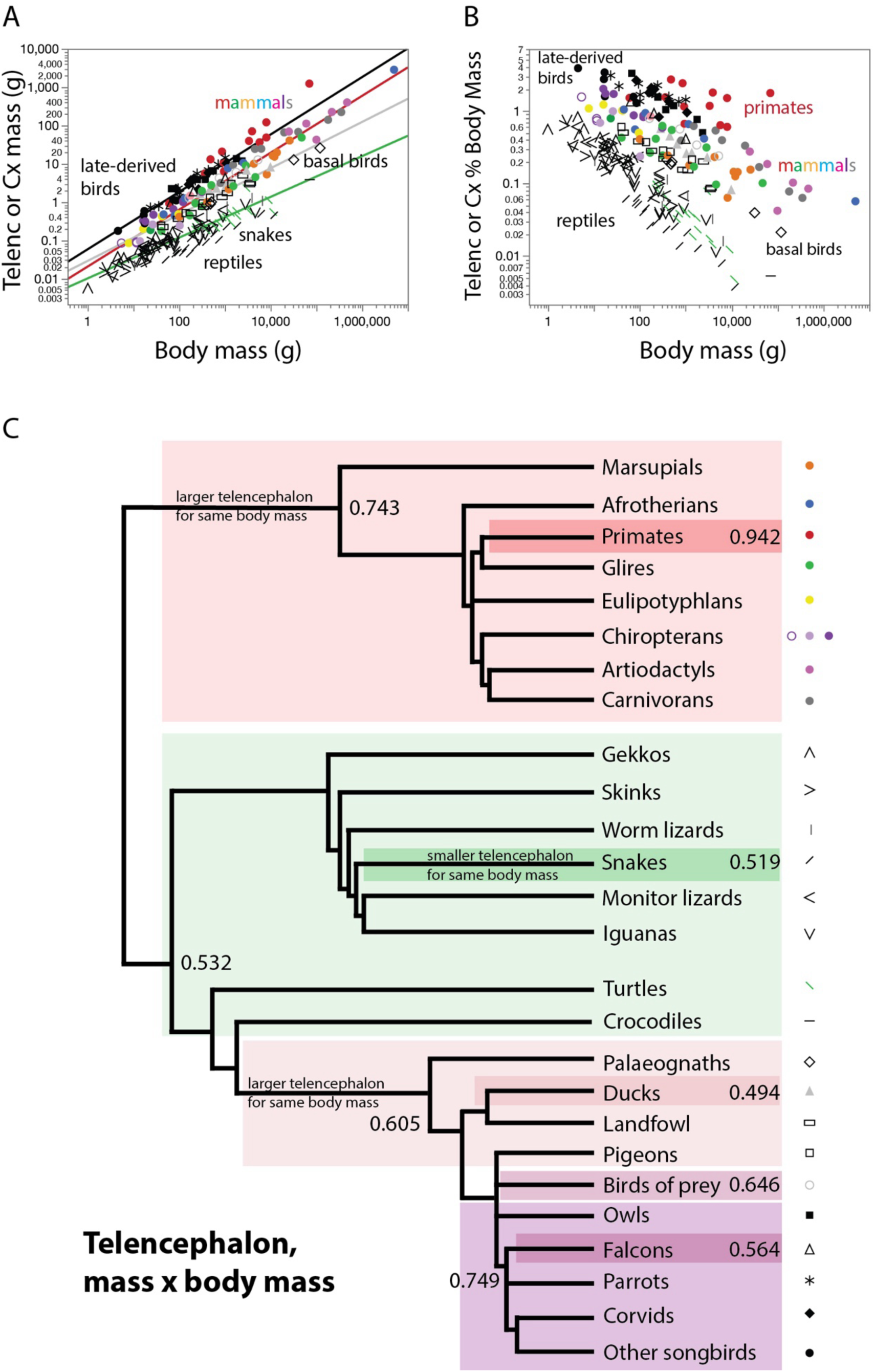
Larger amniotes have a larger pallium (or telencephalon) that is also relatively smaller, in a clade-specific manner. **A**, absolute cortical/pallial/telencephalic mass (in mammals, birds, and reptiles, respectively) increases as different power functions of body mass across reptiles (minus snakes; plotted in green), basal birds (plotted in grey), late-derived birds (plotted in black), and mammals (minus primates; plotted in red). **B**, relative cortical/pallial/telencephalic mass, expressed as a percentage of body mass, decreases with increasing body mass. For the sake of clarity, power functions are not plotted here. **C**, evolutionary tree showing the clades that share similar scaling exponents, which are shown where applicable. Power functions are listed in Table 1. Each data point corresponds to one species. The key used to depict each clade is shown to the right.

Similarly, Figure 3 shows the scaling of the amniote cerebellum, where it is again apparent that larger animals have an absolutely larger but also relatively smaller cerebellum (expressed as a fraction of body mass, not brain mass); that birds and mammals have a larger cerebellum than reptiles of a similar body mass; and that clade-specific scaling rules apply. Most reptiles, with the exception of snakes and iguanas, share an increase in mass of the cerebellum with body mass raised to the power of 0.627±0.034; interestingly, for a given body mass, snakes have a relatively smaller cerebellum compared to the size of the body, and iguanas have a relatively larger cerebellum, with scaling exponents that also veer away from the exponent shared by other clades (Figure 3C). In both mammals and basal birds, there is a shift towards a relatively larger cerebellum compared to the body, with a notable difference to the pallium: endothermic clades have a cerebellum that appears to be truly enlarged, with very little overlap with reptiles in absolute cerebellar mass (Figure 3A). Additionally, there is far less spread amongst endothermic species and clades in the scaling of cerebellar mass compared to the pallium (Figure 2). These key differences and their implications for understanding how brain evolution changed together with endothermy are explored in detail in a separate chapter (Herculano-Houzel, 2025).

**Figure 3.**
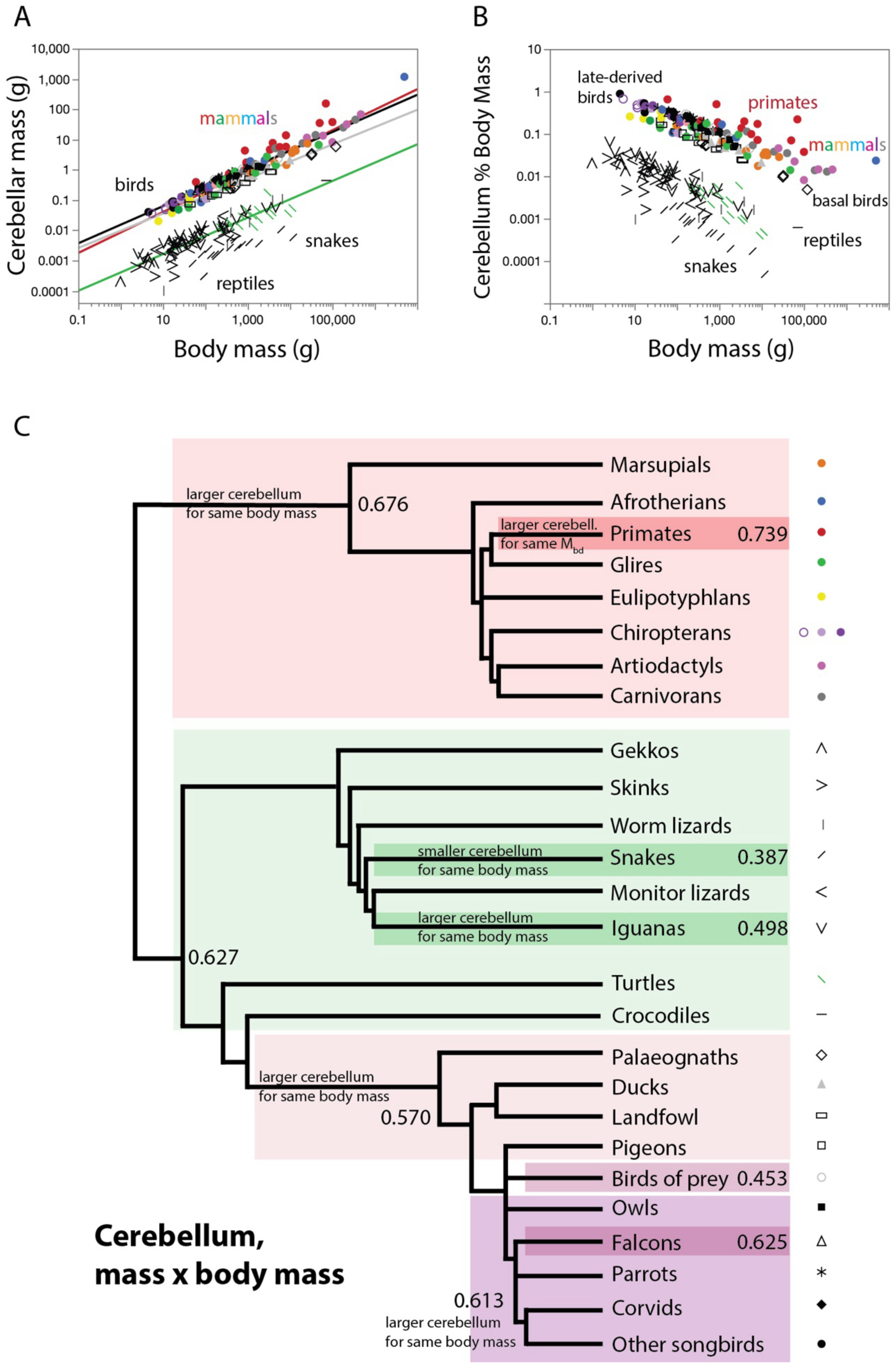
Larger amniotes have a larger cerebellum that is also relatively smaller, in a clade-specific manner. **A**, the absolute mass of the cerebellum increases as different power functions of body mass across reptiles (minus snakes; plotted in green), basal birds (plotted in grey), late-derived birds (plotted in black), and mammals (minus primates; plotted in red). **B**, relative cerebellar mass, expressed as a percentage of body mass, decreases with increasing body mass. For the sake of clarity, power functions are not plotted here. **C**, evolutionary tree showing the clades that share similar scaling exponents, which are shown where applicable. Power functions are listed in Table 1. Each data point corresponds to one species. The key used to depict each clade is shown to the right.

Figure 4 shows the scaling of the amniote rest of brain, which once more is absolutely larger, but also relatively smaller (as a fraction of body mass, not brain mass), in larger species. Again, the rest of brain is larger in birds and mammals than in reptiles of a similar body mass; and clade-specific scaling rules apply. Snakes are once again an exception amongst reptiles in the scaling of brain structure mass with body mass, but here they are joined by turtles and crocodiles, all of which have a smaller rest of brain than other reptiles of a similar body mass, whether with a similar scaling exponent (turtles and crocodiles) or a different one (snakes; Figure 4C). As seen for the pallium, there are some clades within mammals and birds that have a particularly enlarged rest of brain for a given body mass (namely, primates, songbirds, parrots, and owls); and there is a shift in both mammals and basal birds towards a relatively larger rest of brain compared to the body – which, again, could be the result of enlargement of the rest of brain or of a reduction in the size of the body.

**Figure 4.**
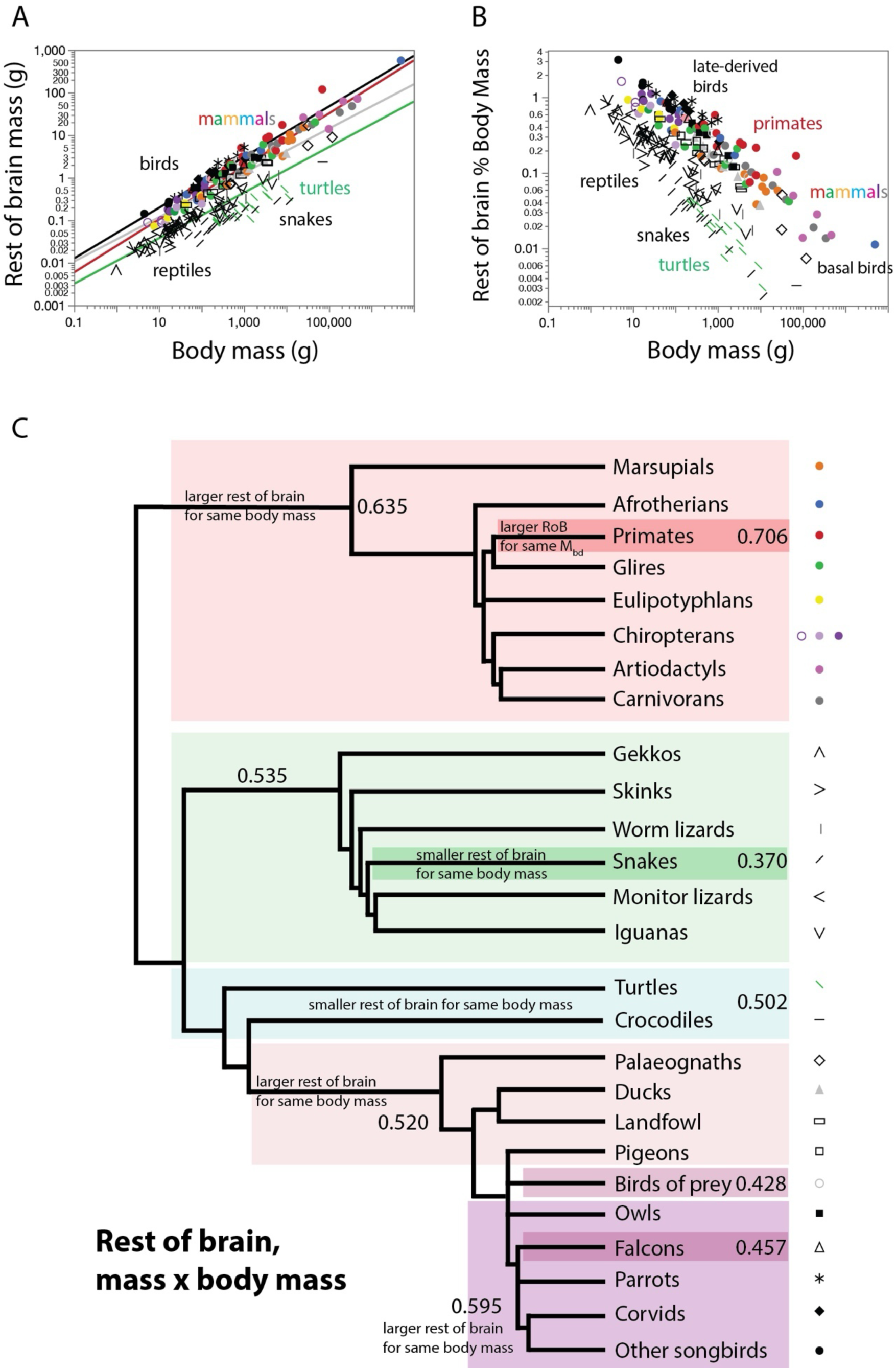
Larger amniotes have a larger rest of brain that is also relatively smaller, in a clade-specific manner. **A**, the absolute mass of the rest of brain increases as different power functions of body mass across reptiles (minus snakes, turtles, and crocodiles; plotted in green), basal birds (plotted in grey), late-derived birds (plotted in black), and mammals (minus primates; plotted in red). **B**, relative cerebellar mass, expressed as a percentage of body mass, decreases with increasing body mass. For the sake of clarity, power functions are not plotted here. **C**, evolutionary tree showing the clades that share similar scaling exponents, which are shown where applicable. Power functions are listed in Table 1. Each data point corresponds to one species. The key used to depict each clade is shown to the right.

### Proportional size of brain structures

While the relative size of the brain and each of its major subdivisions expressed as a percentage of body mass decreases with increasing brain size, the relative proportions of the parts of the brain may or may not scale relative to each other. To avoid confusion, I refer to the latter as *proportional size*: the mass of a brain structure expressed as a percentage of the mass of the whole brain. Thus, whereas the relative size indicates how a part of the brain scales with the body, the proportional size indicates how it scales with the brain.

Figure 5 shows that there is extensive overlap in the proportional size of the pallium/cerebral cortex/telencephalon across reptiles, birds, and mammals; it is only the largest primates, carnivorans and artiodactyls among mammals, and owls and parrots among birds, that have a pallium of distinctively larger proportional size amongst amniotes. Strikingly, while the proportional size of the pallium/telencephalon can be described to slowly scale up together with body mass across all reptiles, and across all mammals in the dataset, close examination clade by clade reveals that the norm is that the proportional size of the pallium/cerebral cortex/telencephalon does *not* scale with body mass. Such scaling is only evident in primates, artiodactyls and carnivorans among mammals, and in parrots and songbirds among birds (Table 2). It is thus *not* the case that a general theme in amniote evolution is the proportional expansion of the telencephalon or pallium.

**Figure 5.**
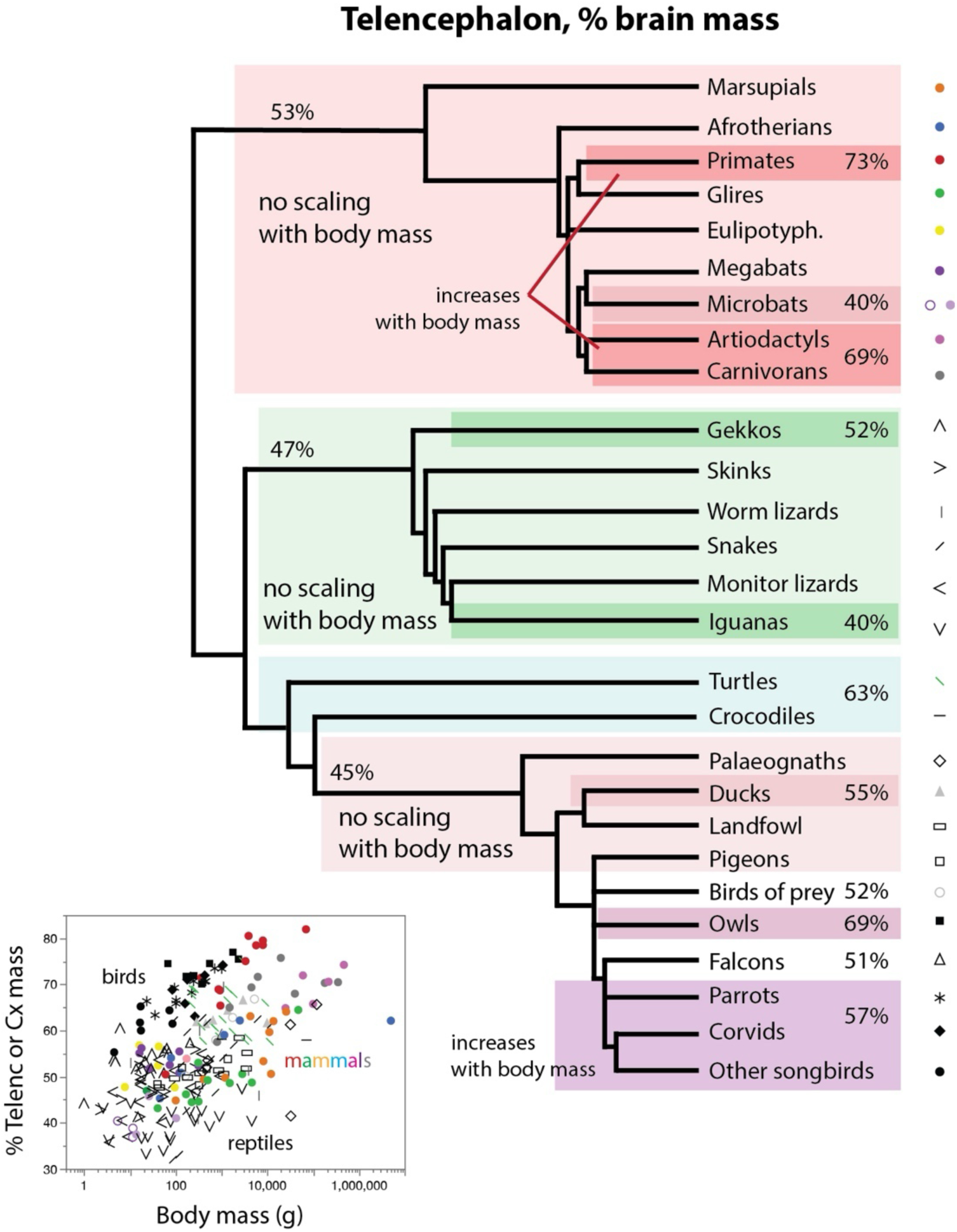
Amniotes have a telencephalon/pallium/cerebral cortex of mostly overlapping proportional size that, with few exceptions, does not scale with body mass. The evolutionary tree depicts the average proportional size of the telencephalon/pallium/cerebral cortex shared by clades in each rectangle. The graph shows the proportional size of the telencephalon/pallium/cerebral cortex plotted against the body mass of each species. For the sake of clarity, no power functions are plotted here, but note that the proportional size of this subdivision of the brain only scales with body mass in the few clades indicated in the schematic. Power functions are listed in Table 2. Each data point corresponds to one species. The key used to depict each clade is shown to the right.

**Table 2.** Scaling of the proportional mass of the different brain structures of reptiles, birds and mammals, expressed as % of total brain mass, as a function of body mass. Values correspond to the equation %Mstr = e^a±SE^ Mbd^b±SE^, the p-values for a and b, and the r^2^ for the power fit, where %Mstr is the proportional mass of a given brain structure and Mbd is the average body mass for the species. Numbers in parentheses are the average relative structure mass ± SD for that clade. *pig excluded.**capybara excluded. ***elephant excluded.

| $\%M_{str} \times M_{bd}$ | n | a in $e^a$ | b in $M_{bd}^b$ | p, a | p, b | $r^2$ |
| --- | --- | --- | --- | --- | --- | --- |
| **Cortex/pallium/telencephalon** |  |  |  |  |  |  |
| **Reptiles** |  |  |  |  |  |  |
| All (50.67±9.61) | 108 | 3.717±0.040 | **0.039**±0.008 | <0.0001 | <0.0001 | 0.202 |
| Anguimorpha (53.16±9.35) | 7 | 3.992±0.240 | **-0.005**±0.036 | <0.0001 | 0.8961 | 0.004 |
| Crocodylia (57.93) | 1 |  |  |  |  |  |
| Gekkota (52.05±4.72) | 14 | 3.886±0.066 | **0.023**±0.022 | <0.0001 | 0.3292 | 0.079 |
| Iguania (40.06±3.83) | 20 | 3.599±0.052 | **0.019**±0.010 | <0.0001 | 0.0879 | 0.153 |
| Lacertoidea (46.38±5.77) | 15 | 3.668±0.067 | **0.045**±0.017 | <0.0001 | 0.0207 | 0.348 |
| Scincoidea (48.77±4.40) | 14 | 3.751±0.054 | **0.038**±0.014 | <0.0001 | 0.0204 | 0.373 |
| Serpentes (51.31±9.91) | 18 | 3.467±0.178 | **0.076**±0.029 | <0.0001 | 0.0191 | 0.298 |
| Testudines (63.71±5.08) | 19 | 4.244±0.109 | **-0.013**±0.015 | <0.0001 | 0.4035 | 0.041 |
| Testudines and Crocodylia (63.42±5.11) | 20 | 4.266±0.087 | **-0.017**±0.012 | <0.0001 | 0.1758 | 0.099 |
| All minus Testudines (46.87±7.07) | 70 | 3.798±0.041 | **0.009**±0.009 | <0.0001 | 0.3145 | 0.015 |
| **Birds** |  |  |  |  |  |  |
| All (53.72±8.69) | 65 | 3.930±0.068 | **0.007**±0.011 | <0.0001 | 0.5328 | 0.006 |
| Accipitriformes (51.93±6.83) | 4 | 3.312±0.025 | **0.091**±0.003 | <0.0001 | 0.0015 | 0.997 |
| Anseriformes (54.59±1.95) | 7 | 3.943±0.090 | **0.008**±0.013 | <0.0001 | 0.5524 | 0.075 |
| Columbiformes (42.84±2.48) | 5 | 3.583±0.099 | **0.031**±0.017 | <0.0001 | 0.724 | 0.515 |
| Falconiformes (50.70±1.90) | 3 | 3.778±0.005 | **0.027**±0.001 | 0.0008 | 0.0212 | 0.999 |
| Galliformes<br>(43.78±3.11) | 9 | 3.565±0.070 | **0.033**±0.011 | <0.0001 | 0.0165 | 0.584 |
| Palaeognathae<br>(47.97±8.36) | 6 | 3.595±0.254 | **0.031**±0.029 | 0.0001 | 0.3423 | 0.225 |
| Passeriformes<br>(56.71±6.38) | 13 | 3.761±0.062 | **0.061**±0.013 | <0.0001 | 0.0008 | 0.656 |
| Psittaciformes<br>(57.67±2.53) | 11 | 3.890±0.033 | **0.032**±0.006 | <0.0001 | 0.0007 | 0.740 |
| Strigiformes<br>(68.86±2.68) | 7 | 4.132±0.072 | **0.016**±0.012 | <0.0001 | 0.2194 | 0.283 |
| Passeriformes,<br>Psittaciformes<br>(57.15±4.92) | 24 | 3.806±0.041 | **0.050**±0.008 | <0.0001 | <0.0001 | 0.615 |
| Galliformes,<br>Columbiformes,<br>Palaeognathae<br>(44.80±5.33) | 20 | 3.571±0.067 | **0.033**±0.009 | <0.0001 | 0.0023 | 0.413 |
| **Mammals** |  |  |  |  |  |  |
| All<br>(58.02±11.72) | 70 | 3.719±0.042 | **0.047**±0.006 | <0.0001 | <0.0001 | 0.513 |
| All non-primates<br>(55.23±10.03) | 57 | 3.711±0.036 | **0.042**±0.005 | <0.0001 | <0.0001 | 0.580 |
| Afrotheria<br>(55.79±6.77) | 6 | 3.859±0.083 | **0.021**±0.010 | <0.0001 | 0.1049 | 0.522 |
| Artiodactyla*<br>(70.41±3.99) | 4 | 3.830±0.205 | **0.036**±0.018 | 0.0029 | 0.1746 | 0.681 |
| Carnivora<br>(68.51±5.39) | 8 | 4.008±0.120 | **0.023**±0.012 | <0.0001 | 0.1148 | 0.362 |
| Chiroptera:<br>Pteropodidae<br>(53.47±2.07) | 7 | 4.044±0.058 | **-0.016**±0.014 | <0.0001 | 0.2956 | 0.214 |
| Chiroptera:<br>Rhinolophoidea,<br>Yangochiroptera<br>(40.04±3.21) | 6 | 3.604±0.108 | **0.029**±0.036 | <0.0001 | 0.4633 | 0.141 |
| Eulipotyphla<br>(52.21±4.48) | 5 | 3.959±0.178 | **-0.002**±0.051 | 0.0002 | 0.9708 | 0.001 |
| Marsupialia<br>(55.20±7.10) | 9 | 3.557±0.135 | **0.055**±0.016 | <0.0001 | 0.0117 | 0.621 |
| Primata<br>(72.64±9.19) | 12 | 3.775±0.094 | **0.065**±0.012 | <0.0001 | 0.0004 | 0.771 |
| Glires**<br>(47.61±3.23) | 9 | 3.689±0.086 | **0.028±0.014** | <0.0001 | 0.0809 | 0.372 |
| Scandentia | 1 |  |  |  |  |  |
| Afrotheria,<br>Eulipotyphla,<br>Glires,<br>Marsupialia,<br>Chiroptera<br>(Pteropodidae)<br>(52.96±5.94) | 38 | 3.840±0.039 | **0.020±0.006** | <0.0001 | 0.0013 | 0.251 |
| Artiodactyla,<br>Carnivora<br>(68.88±4.75) | 13 | 4.019±0.089 | **0.020±0.008** | <0.0001 | 0.0344 | 0.346 |
| **Cerebellum** |  |  |  |  |  |  |
| **Reptiles** |  |  |  |  |  |  |
| All (2.86±1.55) | 108 | 0.572±0.143 | **0.064±0.027** | 0.0001 | 0.0176 | 0.052 |
| Anguimorpha<br>(2.75±1.77) | 7 | -2.003±0.842 | **0.425±0.128** | 0.0631 | 0.0210 | 0.688 |
| Crocodylia (6.73) | 1 |  |  |  |  |  |
| Gekkota<br>(2.38±0.64) | 14 | 0.428±0.157 | **0.147±0.053** | 0.0185 | 0.0166 | 0.392 |
| Iguania<br>(4.26±1.00) | 20 | 1.466±0.159 | **-0.010±0.032** | <0.0001 | 0.7505 | 0.006 |
| Lacertoidea<br>(2.27±0.68) | 15 | 0.796±0.175 | **-0.003±0.044** | 0.0005 | 0.9397 | 0.000 |
| Scincoidea<br>(1.81±0.70) | 14 | 0.256±0.330 | **0.071±0.087** | 0.4524 | 0.4313 | 0.052 |
| Serpentes<br>(1.31±0.54) | 18 | 0.520±0.432 | **-0.056±0.071** | 0.2465 | 0.4419 | 0.037 |
| Testudines<br>(4.31±1.40) | 19 | 1.303±0.468 | **0.015±0.066** | 0.0127 | 0.8206 | 0.003 |
| Testudines and<br>Crocodylia<br>(4.43±1.47) | 20 | 1.037±0.379 | **0.055±0.052** | 0.0136 | 0.2991 | 0.060 |
| All minus<br>Iguania,<br>Testudines,<br>Crocodylia,<br>Serpentes<br>(2.24±0.92) | 50 | 0.406±0.155 | **0.080±0.037** | 0.0117 | 0.0370 | 0.088 |
| **Birds** |  |  |  |  |  |  |
| All (10.36±2.77) | 65 | 2.103±0.115 | **0.033±0.018** | <0.0001 | 0.0773 | 0.049 |
| Accipitriformes (13.22±2.21) | 4 | 3.277±0.200 | **-0.102±0.028** | 0.0037 | 0.0696 | 0.866 |
| Anseriformes (11.51±1.86) | 7 | 1.995±0.361 | **0.062±0.050** | 0.0027 | 0.2739 | 0.232 |
| Columbiformes (13.49±1.03) | 5 | 2.424±0.163 | **0.032±0.029** | 0.0007 | 0.3523 | 0.287 |
| Falconiformes (13.60±1.67) | 3 | 2.122±0.034 | **0.088±0.007** | 0.0102 | 0.0438 | 0.995 |
| Galliformes (12.51±0.40) | 9 | 2.528±0.049 | **-0.000±0.008** | <0.0001 | 0.9726 | 0.002 |
| Palaeognathae (12.51±1.36) | 6 | 2.546±0.197 | **-0.003±0.022** | 0.0002 | 0.9023 | 0.004 |
| Passeriformes (8.67±1.24) | 13 | 2.434±0.095 | **-0.064±0.021** | <0.0001 | 0.0095 | 0.472 |
| Passeriformes: Corvids (7.70±0.65) | 7 | 2.049±0.249 | **-0.002±0.045** | 0.0004 | 0.9654 | 0.000 |
| Passeriformes: others (9.80±0.60) | 6 | 2.251±0.093 | **0.009±0.028** | <0.0001 | 0.7566 | 0.027 |
| Psittaciformes (7.14±1.32) | 11 | 2.598±0.164 | **-0.125±0.031** | <0.0001 | 0.0028 | 0.647 |
| Strigiformes (7.58±0.72) | 7 | 2.175±0.194 | **-0.025±0.032** | <0.0001 | 0.4568 | 0.115 |
| Passeriformes, Psittaciformes (7.97±1.47) | 24 | 2.526±0.097 | **-0.098±0.020** | <0.0001 | <0.0001 | 0.534 |
| Galliformes, Columbiformes, Palaeognathae (12.76±0.98) | 20 | 2.573±0.062 | **-0.004±0.009** | <0.0001 | 0.6228 | 0.014 |
| All minus Passeriformes, Psittaciformes, Strigiformes (12.63±1.48) | 34 | 2.567±0.079 | **-0.006±0.011** | <0.0001 | 0.6196 | 0.008 |
| **Mammals** |  |  |  |  |  |  |
| All (14.07±3.89) | 69 | 2.815±0.071 | **-0.030±0.009** | <0.0001 | 0.0024 | 0.130 |
| All non-primates (14.58±3.81) | 57 | 2.815±0.070 | **-0.025±0.009** | <0.0001 | 0.0115 | 0.111 |
| Afrotheria<br>(15.29±5.44) | 6 | 2.255±0.179 | **0.058±0.022** | 0.0002 | 0.0534 | 0.648 |
| Artiodactyla*<br>(10.52±1.65) | 4 | 1.649±0.855 | **0.060±0.073** | 0.1935 | 0.5000 | 0.250 |
| Carnivora<br>(12.73±2.80) | 8 | 2.547±0.418 | **-0.003±0.043** | 0.0009 | 0.9535 | 0.001 |
| Chiroptera:<br>Pteropodidae<br>(14.22±1.07) | 7 | 2.797±0.208 | **-0.036±0.026** | <0.0001 | 0.2242 | 0.278 |
| Chiroptera:<br>Rhinolophoidea,<br>Yangochiroptera<br>(21.23±4.40) | 6 | 3.155±0.283 | **-0.041±0.095** | 0.0004 | 0.6877 | 0.045 |
| Eulipotyphla<br>(13.60±2.72) | 5 | 2.057±0.263 | **0.159±0.075** | 0.0043 | 0.1252 | 0.598 |
| Marsupialia<br>(14.56±2.56) | 9 | 3.218±0.214 | **-0.068±0.026** | <0.0001 | 0.0329 | 0.501 |
| Primata<br>(11.61±3.81) | 11 | 3.144±0.290 | **-0.095±0.037** | <0.0001 | 0.0293 | 0.427 |
| Glires**<br>(15.10±0.81) | 9 | 2.618±0.080 | **0.015±0.013** | <0.0001 | 0.2622 | 0.175 |
| Scandentia | 1 |  |  |  |  |  |
| Afrotheria***,<br>Eulipotyphla,<br>Glires,<br>Marsupialia,<br>Chiroptera<br>(Pteropodidae)<br>(14.15±2.13) | 36 | 2.708±0.073 | **-0.012±0.011** | <0.0001 | 0.3097 | 0.030 |
| Artiodactyla,<br>Carnivora<br>(12.05±2.52) | 13 | 2.603±0.320 | **-0.013±0.030** | <0.0001 | 0.6760 | 0.016 |
| **Rest of brain** |  |  |  |  |  |  |
| **Reptiles** |  |  |  |  |  |  |
| All (46.46±9.70) | 108 | 4.071±0.046 | **-0.052±0.009** | <0.0001 | <0.0001 | 0.259 |
| Anguimorpha<br>(44.09±8.50) | 7 | 3.838±0.291 | **-0.011±0.044** | <0.0001 | 0.8125 | 0.012 |
| Crocodylia<br>(35.34) | 1 |  |  |  |  |  |
| Gekkota<br>(45.57±4.78) | 14 | 3.890±0.069 | **-0.028±0.023** | <0.0001 | 0.2603 | 0.104 |
| Iguania<br>(55.68±4.20) | 20 | 4.073±0.042 | -0.012±0.008 | <0.0001 | 0.1651 | 0.104 |
| Lacertoidea<br>(51.35±5.62) | 15 | 4.072±0.054 | -0.039±0.014 | <0.0001 | 0.0147 | 0.378 |
| Scincoidea<br>(49.42±4.19) | 14 | 4.034±0.047 | -0.039±0.012 | <0.0001 | 0.0083 | 0.453 |
| Serpentes<br>(47.37±9.59) | 18 | 4.273±0.168 | -0.073±0.028 | <0.0001 | 0.0170 | 0.307 |
| Testudines<br>(31.99±4.76) | 19 | 3.338±0.230 | 0.017±0.033 | <0.0001 | 0.6170 | 0.015 |
| Testudines and<br>Crocodilia<br>(32.15±4.70) | 20 | 3.312±0.182 | 0.020±0.025 | <0.0001 | 0.4197 | 0.037 |
| Anguimorpha,<br>Gekkota,<br>Scincoidea,<br>Serpentes<br>(47.00±7.20) | 53 | 3.953±0.049 | -0.026±0.010 | <0.0001 | 0.0120 | 0.117 |
| Iguania,<br>Lacertoidea<br>(53.82±5.26) | 35 | 4.040±0.039 | -0.014±0.009 | <0.0001 | 0.1103 | 0.075 |
| **Birds** |  |  |  |  |  |  |
| All (27.79±6.73) | 65 | 3.396±0.099 | -0.017±0.016 | <0.0001 | 0.2990 | 0.017 |
| Accipitriformes<br>(27.35±5.20) | 4 | 4.177±0.137 | -0.127±0.019 | 0.0011 | 0.0225 | 0.956 |
| Anseriformes<br>(25.97±1.88) | 7 | 3.608±0.116 | -0.050±0.016 | <0.0001 | 0.0275 | 0.655 |
| Columbiformes<br>(35.73±2.47) | 5 | 3.836±0.076 | -0.047±0.013 | <0.0001 | 0.0390 | 0.805 |
| Falconiformes<br>(28.92±3.14) | 3 | 3.795±0.010 | -0.080±0.002 | 0.0017 | 0.0146 | 0.999 |
| Galliformes<br>(35.81±3.11) | 9 | 3.854±0.087 | -0.044±0.013 | <0.0001 | 0.0131 | 0.609 |
| Palaeognathae<br>(33.33±9.73) | 6 | 3.893±0.453 | -0.050±0.051 | 0.0010 | 0.3859 | 0.191 |
| Passeriformes<br>(25.59±4.48) | 13 | 3.651±0.089 | -0.096±0.019 | <0.0001 | 0.0004 | 0.697 |
| Psittaciformes<br>(23.91±2.02) | 11 | 3.500±0.052 | -0.063±0.010 | <0.0001 | 0.0001 | 0.821 |
| Strigiformes<br>(18.81±1.80) | 7 | 3.207±0.170 | -0.046±0.028 | <0.0001 | 0.1581 | 0.355 |
| Passeriformes,<br>Psittaciformes<br>(24.82±3.60) | 24 | 3.602±0.057 | **-0.084±0.012** | <0.0001 | <0.0001 | 0.706 |
| Galliformes,<br>Columbiformes,<br>Palaeognathae<br>(35.05±5.62) | 20 | 3.852±0.113 | **-0.045±0.016** | <0.0001 | 0.0102 | 0.314 |
| **Mammals** |  |  |  |  |  |  |
| All (27.91±9.56) | 69 | 3.897±0.086 | **-0.093±0.011** | <0.0001 | <0.0001 | 0.497 |
| All non-primates<br>(30.19±8.19) | 57 | 3.892±0.056 | **-0.078±0.007** | <0.0001 | <0.0001 | 0.667 |
| All minus<br>primates,<br>carnivorans,<br>megabats<br>(32.01±7.66) | 42 | 3.919±0.054 | **-0.074±0.007** | <0.0001 | <0.0001 | 0.717 |
| Afrotheria<br>(29.13±9.93) | 6 | 4.021±0.041 | **-0.099±0.005** | <0.0001 | <0.0001 | 0.990 |
| Artiodactyla<br>(19.58±4.12) | 5 | 5.003±0.677 | **-0.177±0.058** | 0.0051 | 0.0560 | 0.755 |
| Carnivora<br>(18.76±3.94) | 8 | 3.568±0.236 | **-0.068±0.024** | <0.0001 | 0.0297 | 0.573 |
| Chiroptera:<br>Pteropodidae<br>(32.31±2.23) | 7 | 3.301±0.085 | **0.042±0.020** | <0.0001 | 0.0907 | 0.467 |
| Chiroptera:<br>Rhinolophoidea,<br>Yangochiroptera<br>(38.73±3.36) | 6 | 3.671±0.132 | **-0.006±0.044** | <0.0001 | 0.8971 | 0.005 |
| Eulipotyphla<br>(34.19±4.67) | 5 | 3.708±0.254 | **-0.054±0.073** | 0.0007 | 0.5113 | 0.155 |
| Marsupialia<br>(30.24±4.88) | 9 | 3.909±0.191 | **-0.063±0.022** | <0.0001 | 0.0289 | 0.518 |
| Primata<br>(15.75±6.17) | 11 | 4.179±0.222 | **-0.194±0.028** | <0.0001 | <0.0001 | 0.841 |
| Glires**<br>(37.29±3.16) | 9 | 3.858±0.097 | **-0.039±0.015** | <0.0001 | 0.0367 | 0.468 |
| Scandentia | 1 |  |  |  |  |  |

Similarly, there is no systematic variation in the proportional size of the cerebellum with increasing body size in any clade of amniotes (Figure 6, Table 2). However, there is a striking difference in the proportional size of the cerebellum between ectothermic (reptiles) and endothermic amniotes (birds and mammals), of 1-4% in the former, depending on the clade versus 8-13% in birds and 12-21% in mammals (average proportional cerebellar mass per clade; see Table 2). Amongst reptiles, the proportional mass of the cerebellum of snakes, iguanas, and turtles and crocodiles stands out from an otherwise shared average of ca. 2% (Figure 6). Amongst mammals, primates, artiodactyls and carnivorans have cerebellar of slightly smaller proportional size (ca. 12% on average, compared to a shared average of 14%), whereas microbats stand out with a proportionately larger cerebellum. Amongst birds, it is those species with the proportionately largest pallium (parrots, songbirds, and also owls) that have a proportionately smaller cerebellum. Additionally, it is only across the ensemble of Passeriformes and Psittaciformes that larger body mass is accompanied by a smaller proportional size of the cerebellum (Table 2).

**Figure 6.**
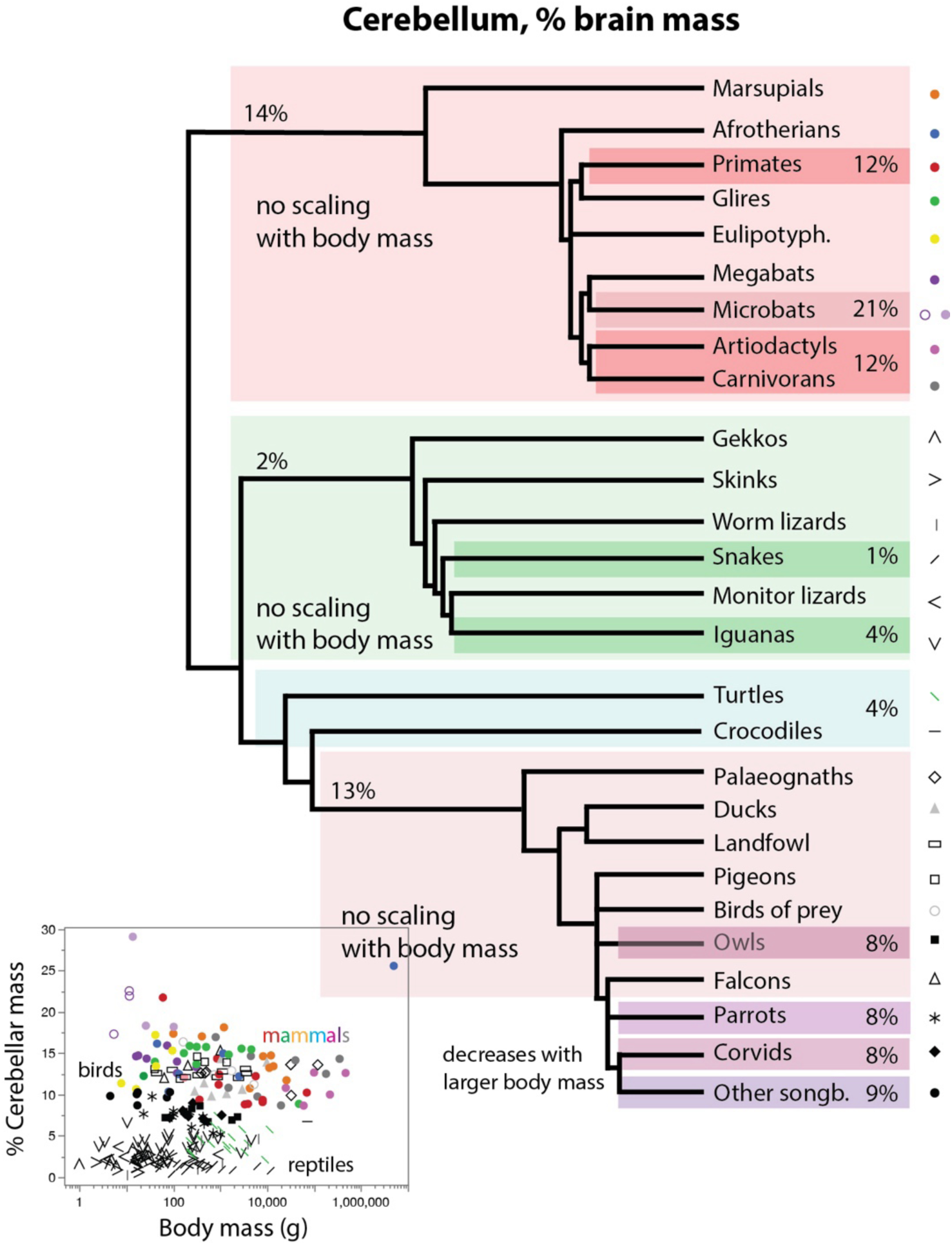
Endothermic amniotes have a proportionately larger cerebellum that does not scale with body mass. The evolutionary tree depicts the average proportional size of the cerebellum shared by clades in each rectangle. The graph shows the proportional size of the cerebellum plotted against the body mass of each species. The proportional size of this subdivision of the brain only scales with body mass (not plotted for clarity) in the ensemble of Passeriformes and Psittaciformes, as indicated in the schematic. Power functions are listed in Table 2. Each data point corresponds to one species. The key used to depict each clade is shown to the right.

As found for the pallium/telencephalon, there is no systematic decrease in the proportional size of the rest of brain across species with increasing body mass within each of the three groups of amniotes (Figure 7). Among mammals, such a decrease is only apparent when a number of clades are pooled together, besides applying specifically within primates, carnivorans, and glires (Table 2). Within marsupials, the correlation is only significant when the largest species are considered. Among birds, there is no significant decrease in the proportional size of the rest of brain across the early-derived clades; the decrease appears later, within the later-derived clades, and even then, the scaling across clades (Figure 7, Table 2). Strikingly, however, there is a large decrease in the proportional size of the rest of brain between the ectothermic reptiles and the endothermic mammals and birds, an increase that parallels the large increase in the proportional size of the cerebellum (but not pallium) in the endothermic groups.

**Figure 7.**
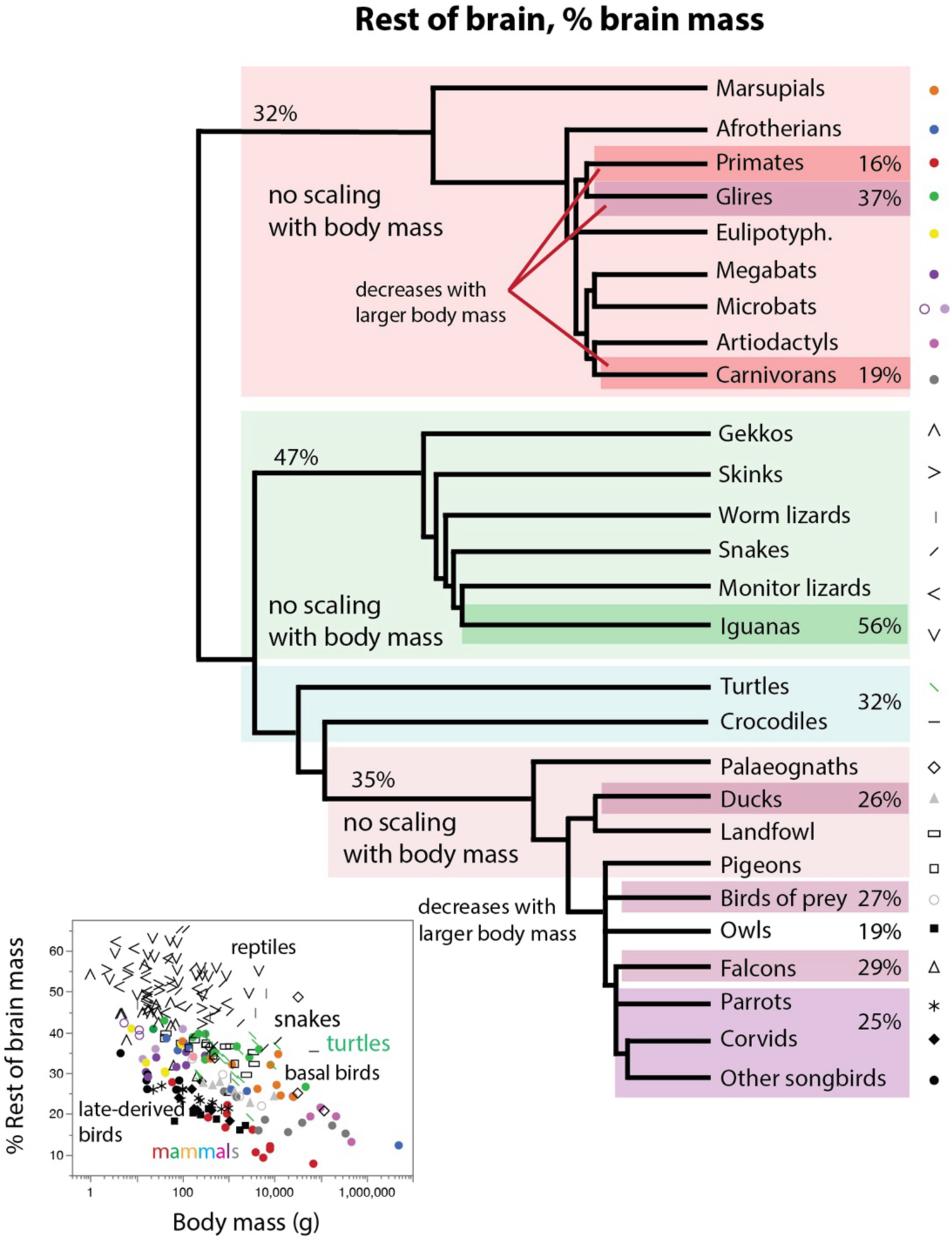
Endothermic amniotes have a rest of brain of decreased proportional mass compared to ectothermic reptiles, but the proportional size of the rest of brain only decreases with body mass within few clades. The evolutionary tree depicts the average proportional size of the rest of brain shared by clades in each rectangle. The graph shows the proportional size of the rest of brain plotted against the body mass of each species. For the sake of clarity, no power functions are plotted here, but note that the proportional size of this subdivision of the brain only scales with body mass in the few clades indicated in the schematic. Power functions are listed in Table 2. Each data point corresponds to one species. The key used to depict each clade is shown to the right.

### Scaling rules for non-neuronal cells are group-specific but not clade-specific

Across the brains of the 242 species of reptiles, mammals, and birds in the combined dataset, which vary in whole brain mass by almost 500,000-fold, the total number of non-neuronal cells varies over 100,000-fold, from just under 2 million in the Algerian sand gecko to 216 billion in the African elephant. At first sight, amniote brains as a whole vary in mass as an essentially linear power function of their numbers of non-neuronal cells (Figure 8A; power fit, exponent 0.992±0.010, r^2^=0.978, p<0.0001; linear fit, r^2^=0.993, p<0.0001). This means that a 1,000 times heavier amniote brain has approximately 1,000 times more non-neuronal cells, with roughly constant densities of non-neuronal cells.

**Figure 8.**
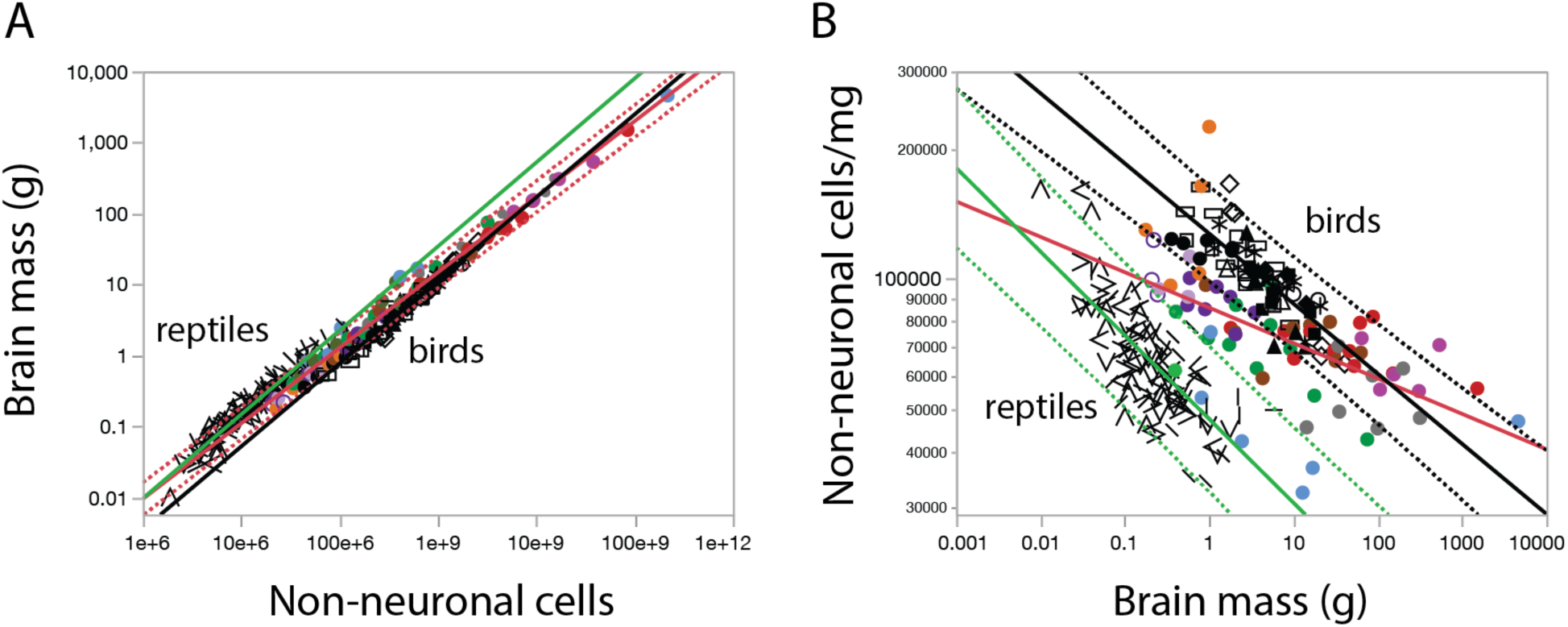
Birds, reptiles and mammals share a nearly linear relationship between brain mass and numbers of non-neuronal cells, although birds and mammals have significantly higher densities of non-neuronal cells than reptiles of similar brain mass. **A**, scaling of brain mass with non-neuronal cells across 108 species of reptiles, 69 species of mammals and 65 species of birds. Power functions are plotted for mammals (in red, with 95% prediction interval: exponent, 1.070±0.013, r^2^=0.990, p<0.0001), reptiles (in green; exponent, 1.184±0.026, r^2^=0.952, p<0.0001) and birds (in black; exponent, 1.170±0.020, r^2^=0.982, p<0.0001). The power function fit across all 242 species has an exponent of 0.992±0.010 that is indistinguishable from linearity. **B**, closer inspection of the ratio between non-neuronal cells and brain mass, which is non-neuronal cell density, makes evident the small but highly significant decrease in non-neuronal cell density with increasing structure mass within each clade (power functions plotted), with exponents -0.081±0.014 (mammals, in red, r^2^=0.354, p<0.0001), -0.192 (reptiles, in green, r^2^=0.556, p<0.0001) and -0.161±0.015 (birds, in black, r^2^=0.658, p<0.0001). Non-neuronal cell densities are overlapping between the larger bird and mammalian brains, but for brains around 1 g, birds have consistently higher non-neuronal cell densities than mammals, which are much higher than non-neuronal cell densities in reptiles. All scaling relationships are listed in Table 3. All symbols as in previous figures.

**Table 3.** Non-neuronal scaling rules for the telencephalon of reptiles and the pallium of birds and mammals; the cerebellum; and the rest of brain (hindbrain to diencephalon in reptiles and birds, hindbrain to striatum in mammals). Values correspond to the equations Mstr = e^a±SE^ O^b±SE^ and Ostr/Mstr = e^a±SE^ M ^b±SE^, the p-values for a and b, and the r^2^ for the power fit, where Mstr is structure mass, Ostr is the number of non-neurons (“other cells”) in the structure, and Ostr/Mstr is the density of non-neuronal (other) cells in the structure.

|  |  |  |  |  |  |  |
| --- | --- | --- | --- | --- | --- | --- |
| **Whole brain** |  |  |  |  |  |  |
| **$M_{str} \times O_{str}$** | **n** | **a in $e^a$** | **b in $N_{str}^b$** | **p, a** | **p, b** | **$r^2$** |
| All amniotes | 242 | -17.980±0.179 | **0.992±0.010** | <0.0001 | <0.0001 | 0.978 |
| All reptiles | 108 | -20.989±0.424 | **1.184±0.026** | <0.0001 | <0.0001 | 0.952 |
| All mammals | 69 | -19.499±0.264 | **1.070±0.013** | <0.0001 | <0.0001 | 0.990 |
| All birds | 65 | -21.811±0.405 | **1.170±0.020** | <0.0001 | <0.0001 | 0.982 |
| **$O_{str} / M_{str} \times M_{str}$** | **n** | **a in $e^a$** | **b in $N_{str}^b$** | **p, a** | **p, b** | **$r^2$** |
| All reptiles<br>(66,439±21,903) | 108 | 10.772±0.031 | **-0.192±0.017** | <0.0001 | <0.0001 | 0.556 |
| All mammals<br>(77,068±28,861) | 68 | 11.362±0.042 | **-0.081±0.014** | <0.0001 | <0.0001 | 0.354 |
| All birds<br>(104,237±21,850) | 65 | 11.754±0.025 | **-0.161±0.015** | <0.0001 | <0.0001 | 0.658 |
| **Telencephalon or cortex** |  |  |  |  |  |  |
| **$M_{str} \times O_{str}$** | **n** | **a in $e^a$** | **b in $N_{str}^b$** | **p, a** | **p, b** | **$r^2$** |
| All amniotes | 242 | -17.331±0.175 | **0.967±0.010** | <0.0001 | <0.0001 | 0.976 |
| All reptiles | 108 | -21.137±0.500 | **1.221±0.032** | <0.0001 | <0.0001 | 0.932 |
| All mammals | 70 | -18.617±0.268 | **1.033±0.014** | <0.0001 | <0.0001 | 0.988 |
| All birds | 65 | -19.531±0.424 | **1.069±0.022** | <0.0001 | <0.0001 | 0.973 |
| All mammals and birds | 135 | -18.950±0.232 | **1.044±0.012** | <0.0001 | <0.0001 | 0.983 |
| **$O_{str} / M_{str} \times M_{str}$** | **n** | **a in $e^a$** | **b in $N_{str}^b$** | **p, a** | **p, b** | **$r^2$** |
| All reptiles<br>(52,454±25,334) | 108 | 10.286±0.049 | **-0.236±0.020** | <0.0001 | <0.0001 | 0.566 |
| All mammals<br>(66,931±17,592) | 70 | 11.137±0.037 | **-0.043**±0.013 | <0.0001 | 0.0011 | 0.145 |
| All birds<br>(83,690±17,249) | 65 | 11.388±0.027 | **-0.089**±0.019 | <0.0001 | <0.0001 | 0.259 |
| **Cerebellum** |  |  |  |  |  |  |
| **M<sub>str</sub> x O<sub>str</sub>** | **n** | **a in e<sup>a</sup>** | **b in N<sub>str</sub><sup>b</sup>** | **p, a** | **p, b** | **r<sup>2</sup>** |
| All amniotes | 245 | -19.728±0.184 | **1.072**±0.011 | <0.0001 | <0.0001 | 0.974 |
| All reptiles | 108 | -20.990±0.424 | **1.184**±0.026 | <0.0001 | <0.0001 | 0.952 |
| All mammals | 72 | -19.501±0.444 | **1.073**±0.024 | <0.0001 | <0.0001 | 0.966 |
| All birds | 65 | -23.076±0.790 | **1.242**±0.044 | <0.0001 | <0.0001 | 0.927 |
| All mammals and<br>reptiles | 180 | -20.053±0.193 | **1.100**±0.012 | <0.0001 | <0.0001 | 0.978 |
| **O<sub>str</sub> /M<sub>str</sub> x M<sub>str</sub>** | **n** | **a in e<sup>a</sup>** | **b in N<sub>str</sub><sup>b</sup>** | **p, a** | **p, b** | **r<sup>2</sup>** |
| All reptiles<br>(156,454±98,672) | 108 | 11.042±0.156 | **-0.152**±0.029 | <0.0001 | <0.0001 | 0.205 |
| All mammals<br>(86,037±37,832) | 72 | 11.281±0.046 | **-0.100**±0.020 | <0.0001 | <0.0001 | 0.262 |
| All birds<br>(146,992±52,510) | 65 | 11.627±0.035 | **-0.254**±0.027 | <0.0001 | <0.0001 | 0.592 |
| **Rest of brain** |  |  |  |  |  |  |
| **M<sub>str</sub> x O<sub>str</sub>** | **n** | **a in e<sup>a</sup>** | **b in N<sub>str</sub><sup>b</sup>** | **p, a** | **p, b** | **r<sup>2</sup>** |
| All amniotes | 242 | -17.980±0.179 | **0.992**±0.010 | <0.0001 | <0.0001 | 0.978 |
| All reptiles | 108 | -20.050±0.415 | **1.129**±0.026 | <0.0001 | <0.0001 | 0.946 |
| All mammals | 70 | -19.804±0.360 | **1.087**±0.019 | <0.0001 | <0.0001 | 0.980 |
| All birds | 65 | -21.227±0.546 | **1.130**±0.029 | <0.0001 | <0.0001 | 0.959 |
| **O<sub>str</sub> /M<sub>str</sub> x M<sub>str</sub>** | **n** | **a in e<sup>a</sup>** | **b in N<sub>str</sub><sup>b</sup>** | **p, a** | **p, b** | **r<sup>2</sup>** |
| All reptiles<br>(68,583±18,770) | 108 | 10.742±0.048 | **-0.162**±0.019 | <0.0001 | <0.0001 | 0.394 |
| All mammals<br>(81,222±26,324) | 70 | 11.318±0.034 | **-0.099**±0.016 | <0.0001 | <0.0001 | 0.364 |
| All birds<br>(127,807±32,384) | 64 | 11.778±0.024 | **-0.189**±0.025 | <0.0001 | <0.0001 | 0.488 |
| **All structures** |  |  |  |  |  |  |
| **M<sub>str</sub> x O<sub>str</sub>** | **n** | **a in e<sup>a</sup>** | **b in N<sub>str</sub><sup>b</sup>** | **p, a** | **p, b** | **r<sup>2</sup>** |
| All amniotes | 242 | -17.980±0.179 | **0.992**±0.010 | <0.0001 | <0.0001 | 0.978 |
| All reptiles | 108 | -20.050±0.415 | **1.129**±0.026 | <0.0001 | <0.0001 | 0.946 |
| All mammals | 212 | -19.318±0.207 | **1.064**±0.011 | <0.0001 | <0.0001 | 0.978 |
| All birds | 194 | -22.124±0.391 | **1.193**±0.021 | <0.0001 | <0.0001 | 0.944 |
| **O<sub>str</sub> / M<sub>str</sub> x M<sub>str</sub>** | **n** | **a in e<sup>a</sup>** | **b in N<sub>str</sub><sup>b</sup>** | **p, a** | **p, b** | **r<sup>2</sup>** |
| All reptiles<br>(92,497±75,164) | 324 | 10.460±0.039 | **-0.246±0.011** | <0.0001 | <0.0001 | 0.626 |
| All mammals<br>(78,139±29,589) | 212 | 11.259±0.023 | **-0.081±0.009** | <0.0001 | <0.0001 | 0.261 |
| All birds<br>(119,496±45,385) | 194 | 11.644±0.017 | **-0.209±0.014** | <0.0001 | <0.0001 | 0.538 |

The power functions calculated separately for each of the three groups of amniotes reveal, however, that each of them deviates significantly from linearity (Figure 8A). That non-linearity is evidence in the scaling of overall non-neuronal cell density in the brain, which is the ratio between numbers of non-neuronal cells and brain mass, shown in Figure 8B). Even more importantly, examination of this ratio, as opposed to the two absolute values in the ratio, reveals that the scaling of numbers of non-neuronal cells with brain mass is not even shared across reptiles, birds, and mammals. The difference between Figures 8A and 8B highlights the need for analyzing ratios (such as cell density) between variables that scale across several orders of magnitude. For a given brain mass, there are more non-neuronal cells per gram of tissue in mammals compared to reptiles, and even more so in birds compared to mammals (Figure 8B), even though reptiles, birds and mammals share the same limited range of densities of non-neuronal cells that vary mostly by ca. 3-fold, concentrated between 50,000 and 120,000 non-neuronal cells/mg of brain (Figure 8B). Thus, the apparent shared and mathematically significant linear scaling of numbers of non-neuronal cells with brain mass across all amniotes in the dataset is not real, since densities of non-neuronal cells vary systematically with brain mass; and different scaling relationships apply to each amniote clade, with higher densities of non-neuronal cells in the brains of endothermic compared to ectothermic sauropsid species (birds, 104,237±21,850; mammals, 77,068±28,861; reptiles, 66,439±21,903 non-neuronal cells/mg; Table 3).

Separate analysis of the scaling of non-neuronal cell density with the mass of each of the three main subdivisions of the amniote brains in the dataset reveals that the pallium of mammals and birds has up to twice higher densities of non-neuronal cells compared to reptilian telencephalons of similar mass (Figure 9A,B). Non-neuronal densities in the pallium vary little but decrease slowly with increasing structure mass across birds and mammals, with negative exponents that are small but non-overlapping and significantly different from zero (-0.089±0.019 and -0.043±0.013, respectively; Figure 9B, Table 3). In contrast, densities of non-neuronal cells drop rapidly in the telencephalon of reptiles as the mass of structure increases, with an exponent of -0.236±0.020 (Figure 9B, Table 3).

**Figure 9.**
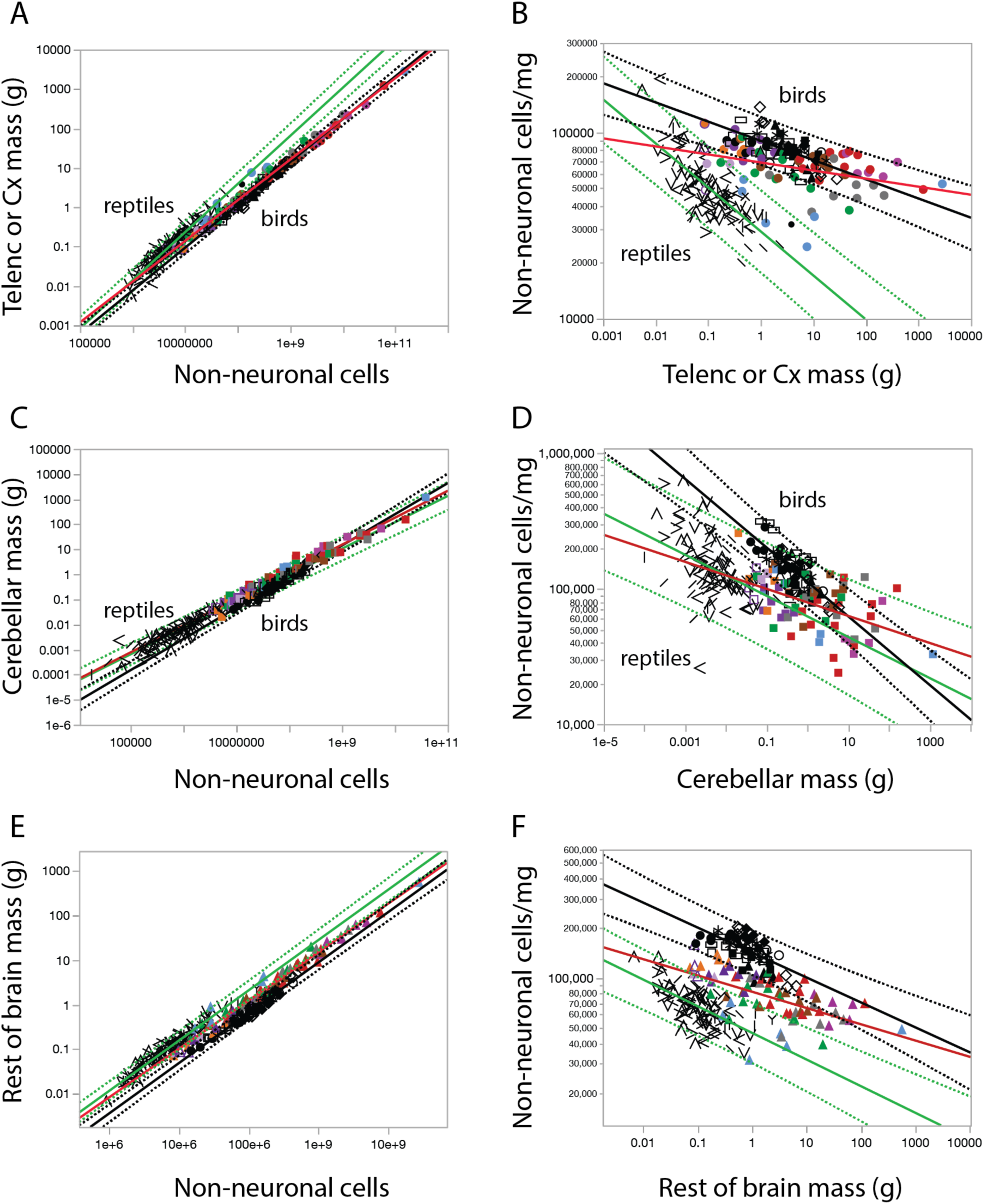
Scaling of density of non-neuronal cells with structure mass differs across brain structures and amniote clades. **A, B**, pallium/cerebral cortex/telencephalon; **C, D**, cerebellum; **E, F**, rest of brain. **A, C, E**: scaling of brain structure mass with number of non-neuronal cells in the structure. **B, D, F**: scaling of non-neuronal cell density with structure mass. Power functions with 95% prediction intervals are plotted for reptiles (green), birds (black), and mammals (red, without the prediction interval for clarity). All symbols as in previous figures. All power functions are listed in Table 3.

In the cerebellum, birds and reptiles share similar non-neuronal cell densities that are also higher than those found in most mammals (Figure 9D). As in the pallium/telencephalon, non-neuronal cell densities scale differently with increasing cerebellar mass in each of the three groups (Figure 9D, Table 3). In the rest of brain, like in the pallium/telencephalon, birds have larger densities of non-neuronal cells than mammals, which in turn are larger than in reptiles, and once more, neuronal cell densities scale differently with increasing rest of brain mass in each of the three groups (Figure 9E,F; Table 3).

In contrast to the clade-specific scaling of absolute, relative and proportional brain structure mass in amniotes, the scaling of numbers of non-neuronal cells with structure mass is shared within each group of amniotes, with no systematic differences across clades.

### Ratios of non-neuronal to neuronal cells are shared between birds and mammals

Across the 242 amniote species in the dataset, the total number of brain neurons varies over 100,000 fold, from just under 2 million in the Algerian sand gecko (Kverkova et al., 2022) to 257 billion in the African elephant (Herculano-Houzel et al., 2014b), which is a similar range as the variation of total numbers of non-neuronal cells. However, neuronal densities vary by over 1,000 times across brain structures and species, whereas non-neuronal densities vary by less than a factor of 10 (Figure 10). As numbers of neurons in the three brain subdivisions increase across species, there is an overall tendency for increasing numbers of neurons to be accompanied by decreasing neuronal densities spanning three orders of magnitude (and, conversely, by average neuronal cell size that increases by the same amount; Mota and Herculano-Houzel, 2014; Figure 10D), while variation in non-neuronal cell densities remains contained within one order of magnitude (Figure 10E).

**Figure 10.**
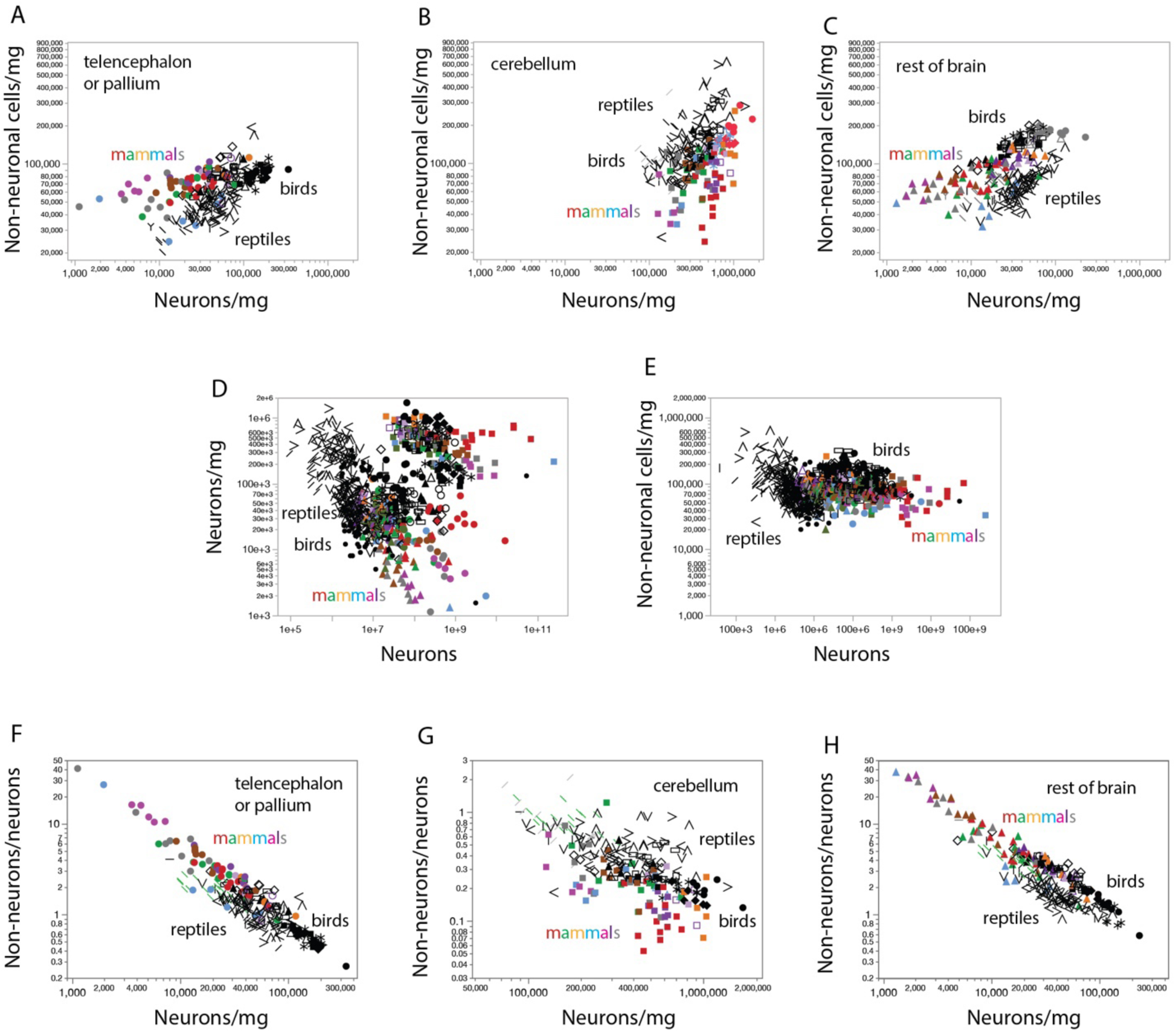
Neuronal cell densities vary 1,000-fold across species and structures, while non-neuronal cell densities vary only ca. 10-fold, but with a relationship that is shared between mammals and birds. **A-C**, comparison of non-neuronal and neuronal cell densities in the telencephalon or pallium (cerebral cortex; **A**), cerebellum (**B**) and rest of brain (**C**). **D, E**, combined dataset contrasting the enormous variation in neuronal densities across species and structures of variable numbers of neurons (**D**) with the small variation in non-neuronal cell densities in the same structures and species (**E**). **F-H**, the scaling of the ratio between numbers of non-neuronal and neuronal cells is shared between mammals and birds in all three brain divisions: pallium (**F**), cerebellum (**G**) and rest of brain (**H**). All symbols as in previous figures.

As noted above, birds have higher densities of non-neuronal cells in the pallium and rest of brain compared to mammalian structures of similar mass (Figure 9), which raises the possibility that densities of glial cells and/or capillary cells have become distinctively higher in the brains of birds. However, the ratio of non-neuronal cells per neuron scales in a similar fashion across birds and mammals (Figure 10F-H). The perfect overlap and continuity in this relationship across the two groups indicates that densities of non-neuronal cells are higher for a same structure mass of the pallium and rest of brain of birds compared to mammals for the simple reason that neuronal densities are higher in birds compared to mammals (Figure 10A,C). Reptiles, in contrast, have consistently lower ratios of non-neuronal cells per neuron for similar neuronal densities in the telencephalon (Figure 10F) and rest of brain (Figure 10H) compared to both birds and mammals, which indicates that the higher ratios of non-neuronal cells per neuron in birds and mammals are associated with the evolution of endothermy, and occurred twice, independently, in each of these two groups.

### Neuronal scaling rules are structure- and clade-specific

The neuronal composition of each of the three brain divisions scales very differently depending on the structure, and also depending on animal clade amongst each group of amniotes, as will be shown in the following sections. Thus, there is no such thing as the neuronal scaling rules for “the brains of mammals” or “the brains of birds”, much less for “the brains of amniotes”. Rather, the neuronal scaling rules that apply to the brains of amniote must be determined separately per brain structure and per clade, as described next. I then devote special attention to the scaling of numbers of neurons *across* brain structures in the different mammalian clades, an important feature of quantitative brain anatomy that informs about the scaling of functional relationships between brain structures in evolution (Herculano-Houzel, 2010; Herculano-Houzel et al., 2014a).

### Neuronal scaling rules for the telencephalon are shared between reptiles and non-primate mammals

It is unfortunate that the study describing the cellular composition of the brains of non-avian reptiles (Kverkova et al., 2022) failed to separate the telencephalon into pallial and subpallial structures, unlike for birds. However, considering that the subpallium contributes only a minority of telencephalic neurons in early derived Palaeognathes and ground-dwelling birds (between 7 and 15%; Olkowicz et al., 2016), and assuming that the same is the case for non-avian reptiles, the neuronal composition of the reptile telencephalon will be considered representative of the reptilian pallium for the purposes of the comparison of scaling rules across clades.

Direct investigation of the relationship between the numbers of neurons that compose the reptilian telencephalon and the mammalian cerebral cortex and the mass of the structures reveals remarkable continuity across all reptilian and mammalian species in the dataset. Reptile telencephalons and non-primate mammalian cerebral cortices of similar mass (which is the case of the smallest mammals in the dataset: eulipotyphlans and bats) are composed of similar numbers of neurons (Figure 11A). The telencephalon/mammalian cerebral cortex scales in mass as a power function of its number of neurons that has statistically indistinguishable exponents between reptiles (1.437±0.064) and non-primate mammals (1.495±0.033; Table 4), and data points for even the largest non-primate mammalian species fall well within the 95% prediction interval for reptiles (Figure 11A). The continuity in neuronal scaling rules between reptiles and mammals, with a combined exponent of 1.377±0.024 (Table 4), strongly indicates that these shared scaling rules already applied to the shared common ancestor between the two clades (Figure 11C).

**Figure 11.**
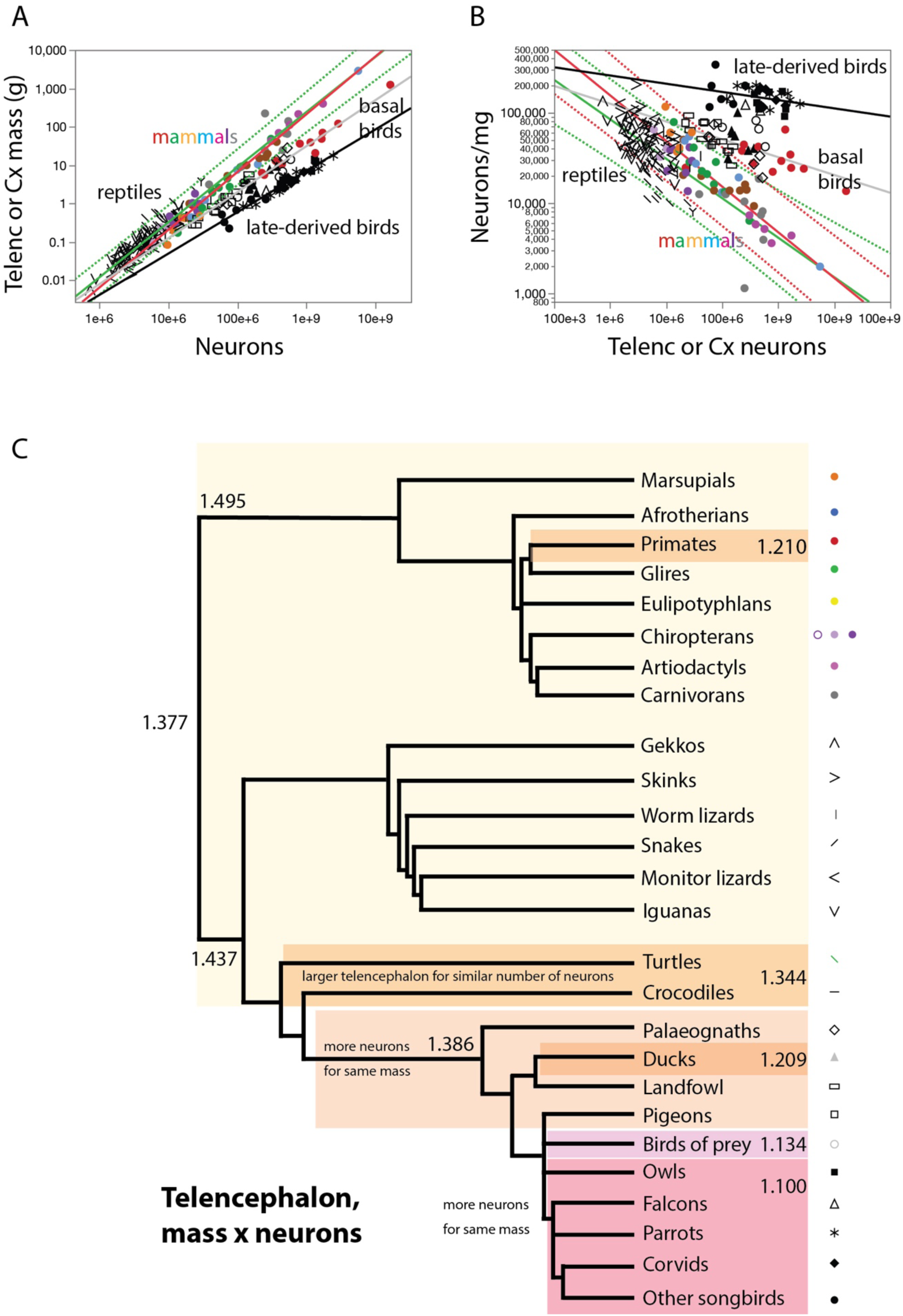
Reptiles and non-primate mammals share neuronal scaling rules for the telencephalon/pallium, while two different scaling rules apply across birds, and primates conform to one of them. **A**, scaling of mass of the telencephalon or pallium with numbers of neurons across non-primate mammals (in red, with 95% prediction interval: exponent, 1.495±0.033), all reptiles (in green; exponent, 1.437±0.064), early-derived birds (Galliformes, Palaeognathae, Columbiformes, in gray; exponent, 1.386±0.060), and late-derived birds (Falconiformes, Passeriformes, Psittaciformes, and Strigiformes, in black; exponent, 1.100±0.044). For the sake of clarity, the 95% prediction interval is only plotted for reptilian species. Note that non-primate mammalian species conform to the prediction interval for reptiles. All power functions are listed in Table 4. **B**, Neuronal density (the ratio between neurons and telencephalic or pallial mass) decreases steeply with increasing numbers of telencephalic or pallial neurons across reptilian and non-primate mammalian species alike, while early-derived birds have higher neuronal densities, even higher in late-derived birds. Power functions plotted have exponents -0.438±0.064 (reptiles, in green), -0.495±0.033 (non-primate mammals, in red), -0.388±0.061 (early-derived birds, in gray) and -0.100±0.044 (late-derived birds, in black). All power functions are listed in Table 5. **C**, Putative cladogram for the evolution of the exponent that describes the scaling relationship between telencephalic mass and number of neurons, showing a shared exponent across most extant reptilian and mammalian species that can be inferred to have applied to ancestral amniotic species. Colored rectangles indicate a shift towards more telencephalic neurons without a change in the scaling exponent at the origin of birds, followed by a shift in scaling exponent at the origin of the recently-derived Passeriformes, Psittaciformes, Falconiformes and Strigiformes. Power functions are listed in Table 4. Each data point corresponds to one species. The key used to depict each clade is shown to the right.

**Table 4.**
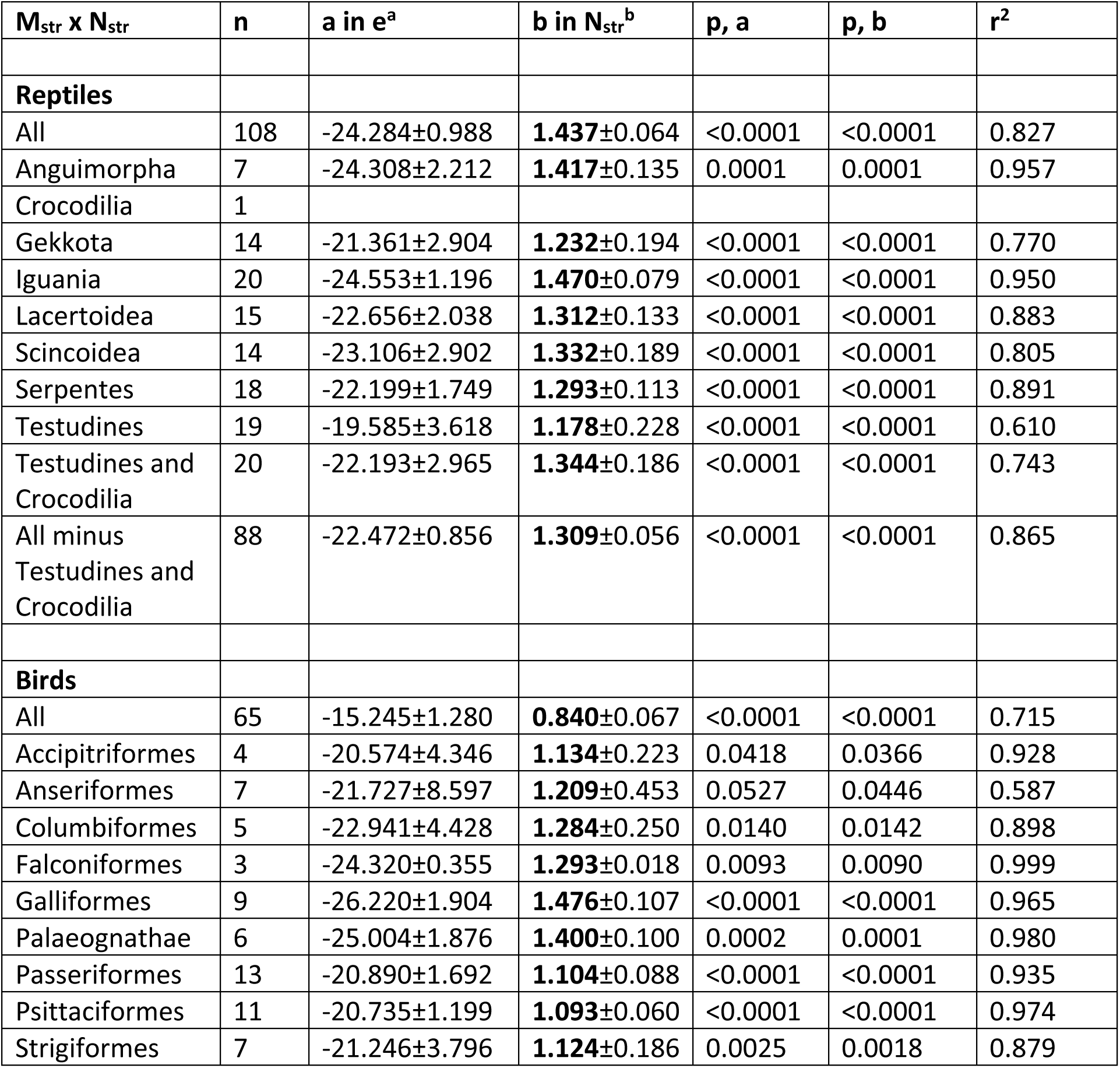
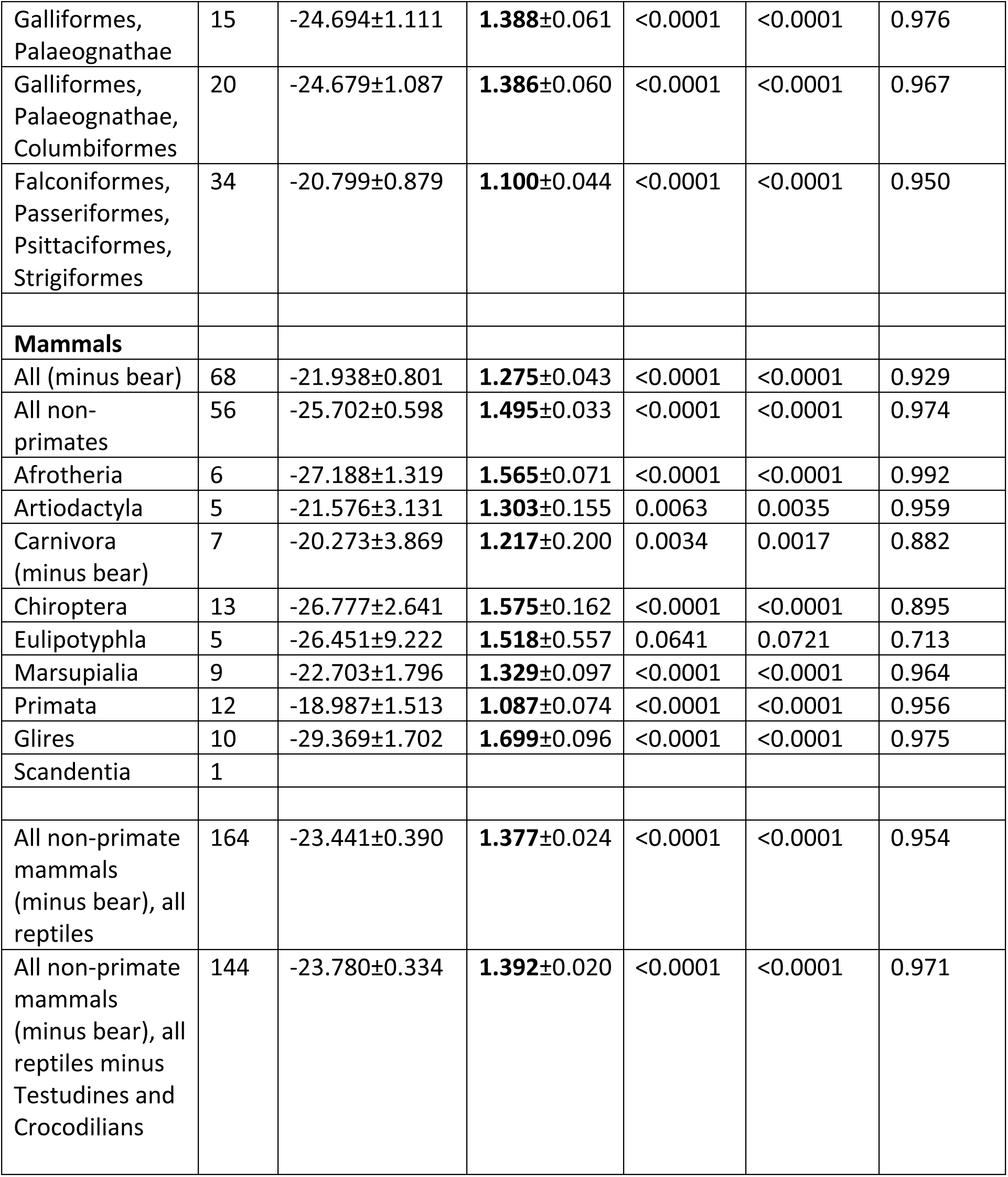
Neuronal scaling rules for the telencephalon of reptiles, and the pallium of birds and mammals. Values correspond to the equation Mstr = e^a±SE^ Nstr^b±SE^, the p-values for a and b, and the r^2^ for the power fit, where Mstr is structure mass and Nstr is the number of neurons in the structure.

**Table 5.** Scaling of neuronal density with varying numbers of neurons in the telencephalon of reptiles, and the pallium of birds and mammals. Values correspond to the equation DNstr = e^a±SE^ Nstr ^b±SE^, the p-values for a and b, and the r^2^ for the power fit, where DNstr is neuronal density (in neurons/mg) and Nstr is the number of neurons in the structure.

| **D<sub>Nstr</sub> x N<sub>Str</sub>** | **n** | **a in e<sup>a</sup>** | **b in N<sub>Str</sub><sup>b</sup>** | **p, a** | **p, b** | **r<sup>2</sup>** |
| --- | --- | --- | --- | --- | --- | --- |
| **Reptiles** |  |  |  |  |  |  |
| All | 108 | 17.390±0.990 | **-0.438±0.064** | <0.0001 | <0.0001 | 0.305 |
| Anguimorpha | 7 | 17.354±2.194 | **-0.414±0.134** | 0.0005 | 0.0273 | 0.656 |
| Crocodylia | 1 |  |  |  |  |  |
| Gekkota | 14 | 14.423±2.902 | **-0.230±0.194** | 0.0003 | 0.2591 | 0.105 |
| Iguania | 20 | 17.756±1.203 | **-0.478±0.080** | <0.0001 | <0.0001 | 0.667 |
| Lacertoidea | 15 | 15.735±2.032 | **-0.311±0.132** | <0.0001 | 0.0352 | 0.298 |
| Scincoidea | 14 | 16.201±2.905 | **-0.332±0.189** | 0.0001 | 0.1050 | 0.204 |
| Serpentes | 18 | 15.323±1.736 | **-0.295±0.112** | <0.0001 | 0.0184 | 0.301 |
| Testudines | 19 | 13.104±3.686 | **-0.204±0.233** | 0.0024 | 0.3920 | 0.043 |
| Testudines and Crocodylia | 20 | 15.486±2.997 | **-0.356±0.188** | <0.0001 | 0.0749 | 0.166 |
| All minus Testudines and Crocodylia | 88 | 15.563±0.858 | **-0.309±0.056** | <0.0001 | <0.0001 | 0.262 |
| **Birds** |  |  |  |  |  |  |
| All | 65 | 8.003±1.273 | **-0.179±0.066** | <0.0001 | 0.0089 | 0.104 |
| Accipitriformes | 4 | 13.794±4.346 | **-0.140±0.223** | 0.0866 | 0.5936 | 0.165 |
| Anseriformes | 7 | 14.721±8.597 | **-0.203±0.453** | 0.1475 | 0.6725 | 0.039 |
| Columbiformes | 5 | 16.826±4.617 | **-0.330±0.260** | 0.0356 | 0.2945 | 0.349 |
| Falconiformes | 3 | 17.404±0.513 | **-0.292±0.027** | 0.0188 | 0.0577 | 0.992 |
| Galliformes | 9 | 19.234±1.899 | **-0.472±0.106** | <0.0001 | 0.0030 | 0.738 |
| Palaeognathae | 6 | 11.964±7.565 | **-0.056±0.402** | 0.1889 | 0.8952 | 0.005 |
| Passeriformes | 13 | 13.982±1.692 | **-0.104±0.088** | <0.0001 | 0.2615 | 0.113 |
| Psittaciformes | 11 | 13.827±1.199 | **-0.093±0.060** | <0.0001 | 0.1531 | 0.213 |
| Strigiformes | 7 | 14.327±3.804 | **-0.123±0.187** | 0.0131 | 0.5380 | 0.080 |
| Galliformes, Palaeognathae | 15 | 13.398±3.092 | **-0.386±0.061** | <0.0001 | <0.0001 | 0.755 |
| Galliformes, Palaeognathae, Columbiformes | 20 | 17.795±1.091 | **-0.388±0.061** | <0.0001 | <0.0001 | 0.697 |
| Falconiformes, Passeriformes, Psittaciformes, Strigiformes | 34 | 13.895±0.879 | **-0.100±0.044** | <0.0001 | 0.0315 | 0.136 |
| **Mammals** |  |  |  |  |  |  |
| All (minus bear) | 68 | 15.023±0.803 | **-0.274±0.044** | <0.0001 | <0.0001 | 0.375 |
| All non-primates | 56 | 18.793±0.600 | **-0.495±0.033** | <0.0001 | <0.0001 | 0.803 |
| Afrotheria | 6 | 20.260±1.332 | **-0.564±0.071** | 0.0001 | 0.0014 | 0.940 |
| Artiodactyla | 5 | 14.668±3.131 | **-0.303±0.155** | 0.0184 | 0.1447 | 0.562 |
| Carnivora<br>(minus bear) | 7 | 13.365±3.870 | **-0.217±0.200** | 0.0182 | 0.3258 | 0.192 |
| Chiroptera | 13 | 19.803±2.656 | **-0.571±0.163** | <0.0001 | 0.0050 | 0.526 |
| Eulipotyphla | 5 | 19.590±9.296 | **-0.521±0.561** | 0.1257 | 0.4217 | 0.223 |
| Marsupialia | 9 | 15.795±1.796 | **-0.329±0.097** | <0.0001 | 0.0117 | 0.620 |
| Primata | 12 | 12.094±1.513 | **-0.087±0.074** | <0.0001 | 0.2644 | 0.123 |
| Glires | 10 | 22.415±1.683 | **-0.696±0.095** | <0.0001 | <0.0001 | 0.870 |
| Scandentia | 1 |  |  |  |  |  |
| All non-primate<br>mammals<br>(minus bear), all<br>reptiles | 164 | 16.526±0.391 | **-0.377±0.024** | <0.0001 | <0.0001 | 0.605 |
| All non-primate<br>mammals,<br>reptiles minus<br>Testudines and<br>Crocodilians | 144 | 16.863±0.335 | **-0.391±0.020** | <0.0001 | <0.0001 | 0.722 |

The shared scaling relationship across reptiles and most mammals between numbers of telencephalic neurons and telencephalic (or cerebral cortical) mass with an exponent of roughly 1.4 means that as the structure gains 10x more neurons, measured neuronal density drops and its inverse, which is average neuronal size, increases by 10^0.4^ = 2.5-fold (Mota and Herculano-Houzel, 2014) in a similar manner across reptiles and non-primate mammals (Figure 11B). The mammalian species with the fewest cerebral cortical neurons, which are bats and eulipotyphlans, have similar neuronal densities as many of the reptiles in the dataset with matching numbers of telencephalic neurons; and as mammalian cortices gain more cortical neurons, their average neuronal density decreases in a way that can be predicted equally well using the power function calculated for non-primate mammals (exponent -0.495±0.033) or for the reptilian telencephalon (exponent -0.438 ± 0.064; Figure 11B; Table 5). In contrast, Testudines and the Nile crocodile have lower neuronal densities than predicted for reptilian telencephalons of their numbers of neurons (Figure 12B).

**Figure 12.**
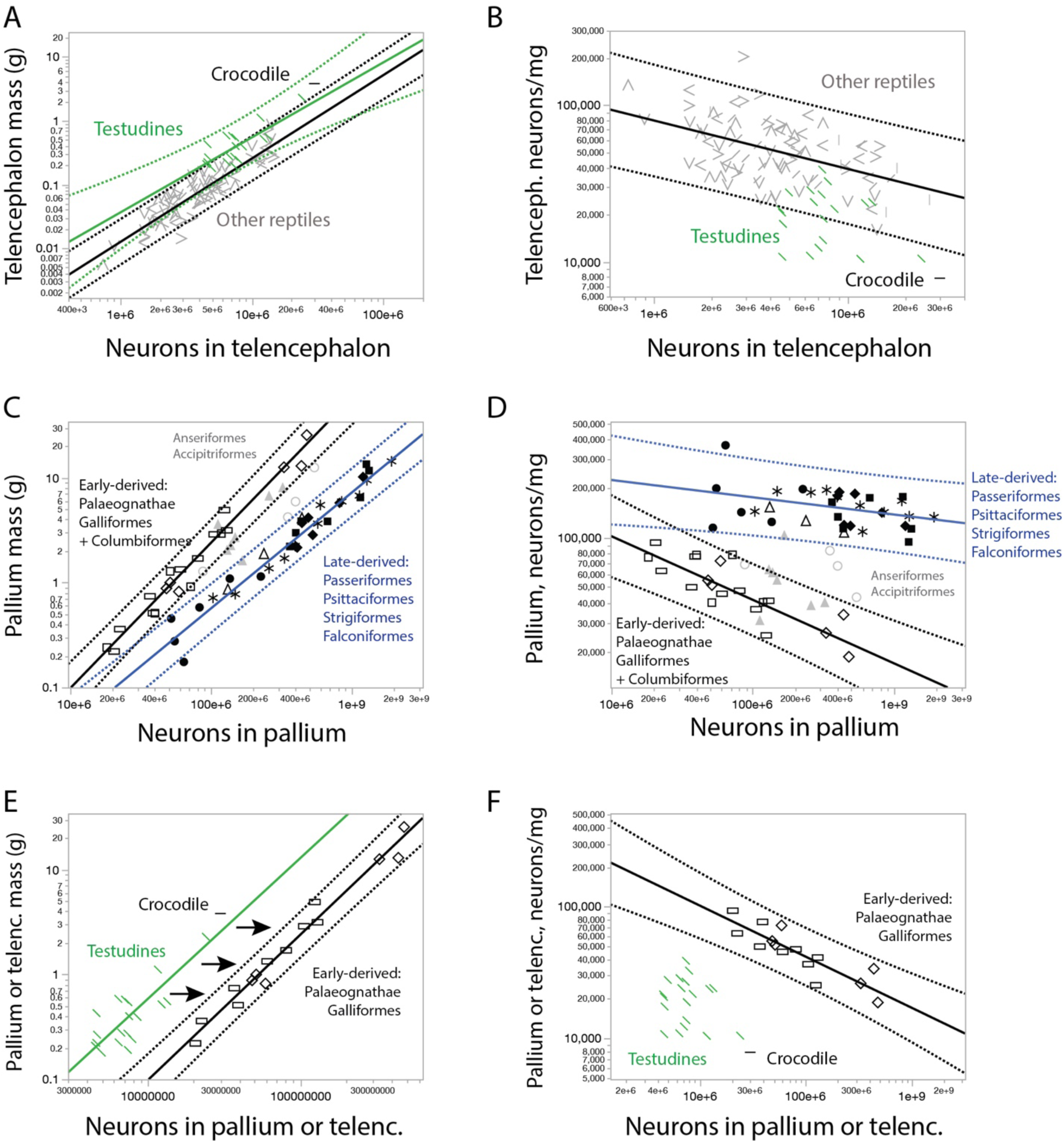
Neuronal scaling rules for the telencephalon of reptiles and birds. **A, C, E,** scaling of telencephalic mass with numbers of telencephalic neurons. **B, D, F,** scaling of neuronal density in the telencephalon with numbers of telencephalic neurons. **A, B,** Testudines and crocodile have larger telencephalons for their numbers of neurons, and thus lower neuronal densities, compared to other reptilian species. **C, D,** distinct neuronal scaling rules apply to the telencephalon of different bird clades, with higher neuronal densities and therefore more neurons in a telencephalon of similar mass in late-derived clades compared to early-derived clades. Anseriformes and Accipitriformes do not conform systematically to either scaling relationships. **E, F,** neuronal scaling rules that apply to the telencephalon of early-derived bird species and Testudines plus crocodile have similar exponents, indicative of a grade shift. All power functions are listed in Tables 4 and 5. Each data point corresponds to one species. Symbols as in all other figures.

Closer examination of each clade in separate reveals that the neuronal scaling rules for the telencephalon are shared across all clades of reptiles examined, with the exception of the Nile crocodile and the Testudines (Table 4), which have systematically larger telencephalons than predicted for the number of telencephalic neurons in a reptile (Figure 12A). While the crocodile telencephalon lies outside the 95% prediction interval for reptilian species, it fits the scaling rules for Testudines (Figure 12A), a finding that is in line with the evolutionary relatedness of Crocodilia and Testudines (Figure 11C). Combining Testudines and the Nile crocodile yields a scaling relationship with exponent 1.344±0.186 (Table 4), with a scaling relationship for all other reptile telencephalons of exponent 1.309±0.056 (Table 4). The similarity between the exponents indicates a grade shift in the neuronal composition of the telencephalon between Crocodilia and Testudines on the one hand, and all other extant non-avian Sauropsids on the other.

The bird telencephalon, in contrast, gains neurons in a clearly distinct manner from non-avian reptilians – but at least two groups are clearly recognizable (Figure 11). In the first, composed by Palaeognathae and Galliformes, which in this dataset are the earliest derived in avian evolution that arose prior to the K-Pg extinction event (Brusatte et al., 2015), the telencephalon gains neurons with a pronounced drop in neuronal density, similar to that observed in reptiles and most mammals, with an exponent of -0.386±0.061 (Figure 11B, Figure 12C,D; Table 5). This scaling relationship captures the later arising Columbiformes very well, such that addition of Columbiformes species leaves the scaling exponent virtually unchanged (- 0.388±0.061; Figure 12C,D, Table 5). Remarkably, while the range of neuronal densities is shared with reptiles and mammals, the neuronal scaling rules for the pallium of these early-arising birds appear shifted to the right compared to Testudines and the Nile crocodile (Figure 12E), as if the same brain structure volume now simply accommodated increased numbers of neurons (that is, with increased neuronal density) while maintaining the same allometry across species. The second group of birds is formed by the later, post-K-Pg-derived Falconiformes, Strigiformes, Passeriformes (including Corvidea) and Psittaciformes. In this group, telencephalic neuronal densities are much higher than in the vast majority of reptiles and mammals (Figure 11B), and barely decrease with varying numbers of neurons (-0.100±0.044, p=0.0315; Figure 12D, Table 5). As a result of these clade- and group-specific scaling rules, early-derived birds have more neurons in the telencephalon than reptiles or non-primate mammals of similar pallial or telencephalic mass; and later-derived birds have even more neurons concentrated in structures of a same mass. Anseriformes and Accipitriformes species lie in between the two clades (Figure 12D).

Mapping these scaling relationships onto the phylogenetic tree of all species in the dataset strongly suggests that the telencephalic scaling rule shared by modern (non-avian) reptiles and non-primate mammals already applied to their common ancestral species (Figure 11C). Birds could have arisen with an increase in numbers of neurons generated in the same volume, causing a shift towards higher neuronal densities, that remains in Palaeognathes such as the emu and ostrich, landfowl, and pigeons; and further later shifts towards even higher neuronal densities at the more recent derivation of songbirds, hawks, owls and parrots (Figure 11C).

### Neuronal scaling rules for the cerebellum are shared across most endothermic species, with a step change from ectothermic reptiles

Differently from the telencephalon, the neuronal scaling rules that apply to the cerebellum are clearly distinct between reptiles and mammals, and overlap across most mammalian and the early-derived bird species (Figure 13). Amongst most reptiles, the cerebellum gains mass faster than it gains neurons, in a manner described by a power function of exponent 1.271±0.051 (Figure 13A, green; Table 6) from which snakes and iguanas are the only clear exceptions (Figure 14). Thus, across most reptilian species as a whole, more cerebellar neurons are also on average larger neurons, with decreasing neuronal densities (Figure 7B), although Iguania is the only clade that shows on its own a clear decrease in cerebellar neuronal density as the cerebellum gains neurons across species (Table 7).

**Figure 13.**
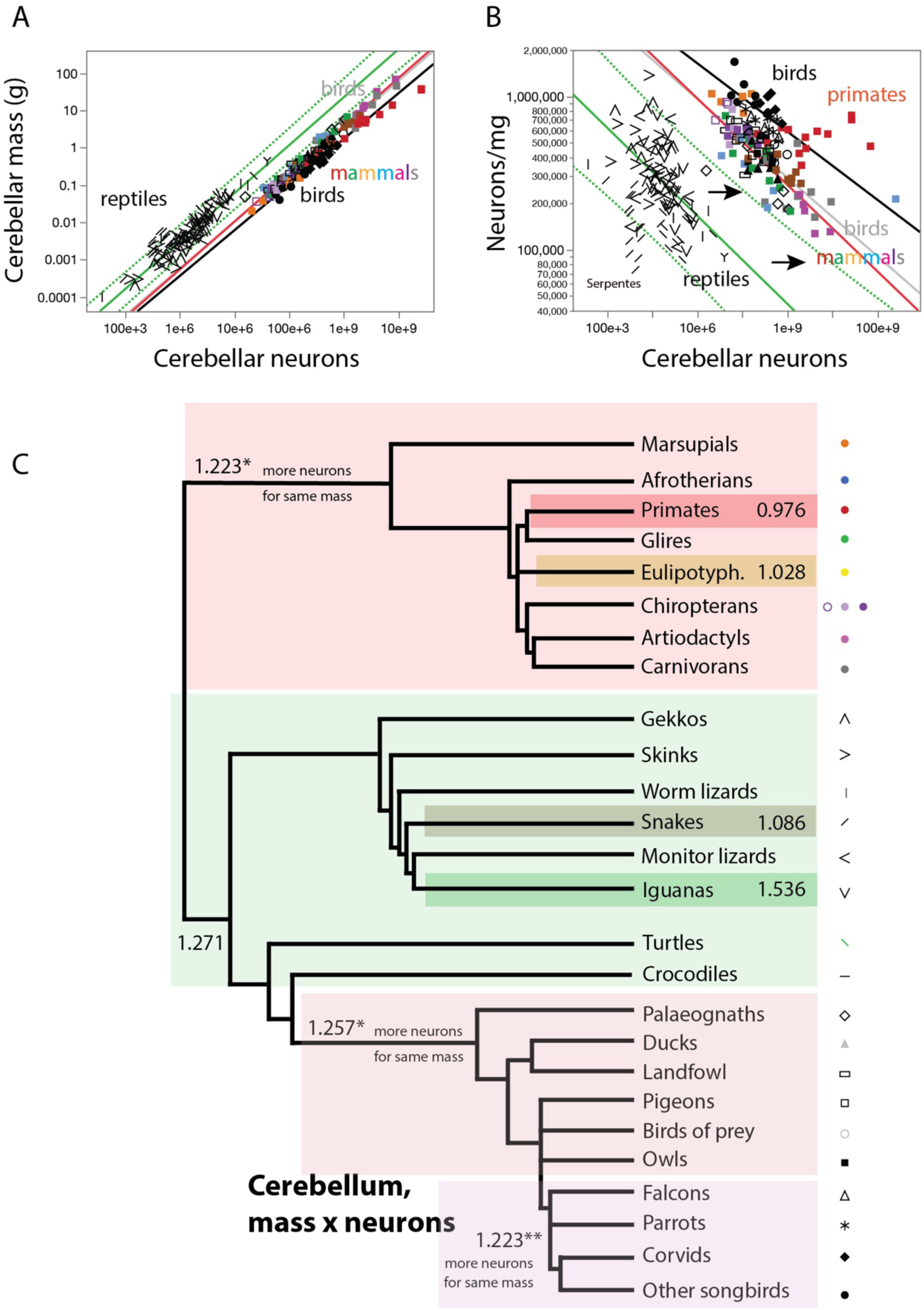
Reptiles, birds and non-primate, non-eulipotyphlans mammals share allometric exponents for the neuronal composition of the cerebellum, although relationships are shifted to the right in the two endotherm clades. **A**, scaling of cerebellar mass with numbers of neurons across non-primate, non-eulipotyphlan mammals (in red, exponent 1.223±0.024), all reptiles minus serpents and iguanas (in green, with 95% prediction interval; exponent, 1.271±0.051), early-derived birds (Galliformes, Palaeognathae, Anseriformes, plus Columbiformes, Accipitriformes and Strigiformes, in gray, with near-perfect overlap with mammals; exponent, 1.257±0.048), and late-derived birds (Falconiformes, Passeriformes, Psittaciformes, in black; exponent, 1.223±0.053). For the sake of clarity, the 95% prediction interval is only plotted for reptilian species. All power functions are listed in Table 6. **B**, Neuronal density in the cerebellum (number of neurons per milligram of cerebellar mass) decreases steeply with increasing numbers of cerebellar neurons in parallel scaling functions across reptilian, non-primate, non-eulipotyphlan mammalian, and early-derived bird species alike, while late-derived birds have higher neuronal densities. Power functions plotted have exponents -0.287±0.046 (reptiles minus serpents, in green), -0.223±0.024 (non-primate, non-eulipotyphlan mammals, in red), -0.257±0.048 (early-derived birds, in gray) and -0.223±0.053 (late-derived birds, in black). For the sake of clarity, the 95% prediction interval is only plotted for reptilian species. All power functions are listed in Table 7. **C**, Putative cladogram for the evolution of the exponent that describes the scaling relationship between cerebellar mass and number of neurons, with a shared exponent across most extant reptilian, basal bird, and mammalian species, with a grade shift towards more neurons in a similar cerebellar mass in the newly endothermic species, then again in primates, eulipotyphlans, and late-derived birds. The scaling relationship shared by most reptilian species can thus be inferred to have applied to ancestral ectothermic amniotic species. Each data point corresponds to one species. The key used to depict each clade is shown to the right.

**Figure 14.**
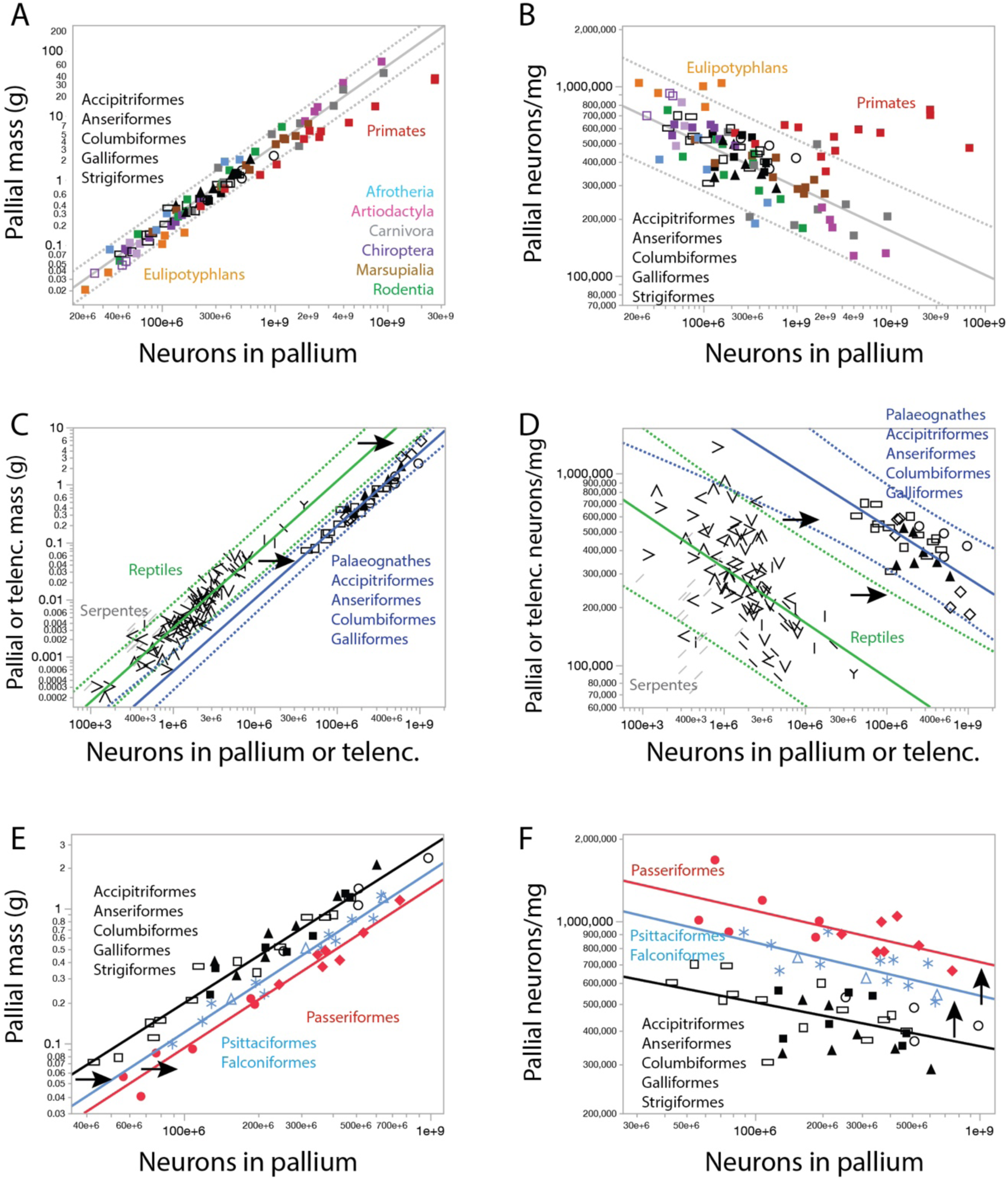
Neuronal scaling rules for the cerebellum of mammals, reptiles and birds. **A, C, E,** scaling of cerebellar mass with numbers of cerebellar neurons. **B, D, F,** scaling of neuronal density in the cerebellum with numbers of cerebellar neurons. **A, B,** Early-derived birds and mammals other than primates and eulipotyphlans share neuronal scaling rules for the cerebellum. Power function plotted applies to non-primate, non-eulipotyphlan mammalian species plus the bird clades indicated. **C, D,** neuronal scaling rules for the cerebellum of reptiles (minus Serpentes) and of early-derived bird clades have similar exponents, indicative of a grade shift. **E, F,** neuronal scaling rules for the cerebellum of three different groups of birds, indicated in black, blue and red, have similar exponents, indicative of two grade shifts in bird evolution. All power functions are listed in Tables 6 and 7.

**Table 6.** Neuronal scaling rules for the cerebellum of reptiles, birds and mammals. Values correspond to the equation Mcb = e^a±SE^ Ncb^b±SE^, the p-values for a and b, and the r^2^ for the power fit, where Mcb is cerebellar mass and Ncb is the number of neurons in the cerebellum.

| $M_{cb} \times N_{cb}$ | n | a in $e^a$ | b in $N_{cb}^b$ | p, a | p, b | $r^2$ |
| --- | --- | --- | --- | --- | --- | --- |
| **Reptiles** |  |  |  |  |  |  |
| All | 108 | -21.597±0.658 | **1.159**±0.046 | <0.0001 | <0.0001 | 0.856 |
| Anguimorpha | 7 | -20.551±1.135 | **1.099**±0.077 | <0.0001 | <0.0001 | 0.976 |
| Crocodylia | 1 |  |  |  |  |  |
| Gekkota | 14 | -22.341±2.108 | **1.179**±0.153 | <0.0001 | <0.0001 | 0.833 |
| Iguania | 20 | -27.286±1.536 | **1.536**±0.148 | <0.0001 | <0.0001 | 0.857 |
| Lacertoidea | 15 | -21.123±2.516 | **1.111**±0.179 | <0.0001 | <0.0001 | 0.747 |
| Scincoidea | 14 | -22.206±2.128 | **1.179**±0.155 | <0.0001 | <0.0001 | 0.828 |
| Serpentes | 18 | -20.114±2.679 | **1.086**±0.206 | <0.0001 | <0.0001 | 0.635 |
| Testudines | 19 | -19.549±1.908 | **1.043**±0.125 | <0.0001 | <0.0001 | 0.803 |
| Testudines and Crocodylia | 20 | -22.828±1.500 | **1.128**±0.098 | <0.0001 | <0.0001 | 0.881 |
| All minus Serpentes | 90 | -23.527±0.673 | **1.285**±0.047 | <0.0001 | <0.0001 | 0.897 |
| All minus Serpentes, Iguania | 70 | -23.313±0.729 | **1.271**±0.051 | <0.0001 | <0.0001 | 0.903 |
| **Birds** |  |  |  |  |  |  |
| All | 65 | -24.549±1.213 | **1.229**±0.063 | <0.0001 | <0.0001 | 0.859 |
| Accipitriformes | 4 | -23.486±3.313 | **1.178**±0.165 | 0.0193 | 0.0191 | 0.962 |
| Anseriformes | 7 | -23.887±3.008 | **1.214**±0.155 | 0.0005 | 0.0005 | 0.924 |
| Columbiformes | 5 | -22.998±2.267 | **1.156**±0.121 | 0.0020 | 0.0024 | 0.968 |
| Falconiformes | 3 | -24.558±0.304 | **1.220**±0.016 | 0.0079 | 0.0081 | 0.999 |
| Galliformes | 9 | -23.365±1.703 | **1.180**±0.090 | <0.0001 | <0.0001 | 0.961 |
| Palaeognathae | 6 | -30.001±1.405 | **1.529**±0.072 | <0.0001 | <0.0001 | 0.991 |
| Passeriformes | 13 | -24.284±1.171 | **1.188**±0.061 | <0.0001 | <0.0001 | 0.972 |
| Psittaciformes | 11 | -23.931±1.264 | **1.183**±0.065 | <0.0001 | <0.0001 | 0.974 |
| Strigiformes | 7 | -22.131±3.078 | **1.114**±0.159 | 0.0008 | 0.0009 | 0.908 |
| Galliformes,<br>Anseriformes,<br>Accipitriformes,<br>Columbiformes,<br>Strigiformes | 32 | -23.000±0.896 | **1.160**±0.047 | <0.0001 | <0.0001 | 0.954 |
| Galliformes,<br>Anseriformes,<br>Accipitriformes,<br>Columbiformes,<br>Strigiformes,<br>Palaeognathae | 38 | -24.822±0.928 | **1.257**±0.048 | <0.0001 | <0.0001 | 0.950 |
| Falconiformes,<br>Passeriformes,<br>Psittaciformes | 27 | -24.822±1.025 | **1.223**±0.053 | <0.0001 | <0.0001 | 0.955 |
| Falconiformes,<br>Psittaciformes | 14 | -24.142±1.039 | **1.195**±0.053 | <0.0001 | <0.0001 | 0.977 |
| **Mammals** |  |  |  |  |  |  |
| All | 71 | -22.539±0.553 | **1.135**±0.028 | <0.0001 | <0.0001 | 0.961 |
| All non-primates | 58 | -24.586±0.495 | **1.245**±0.025 | <0.0001 | <0.0001 | 0.977 |
| All non-primates, non-eulipotyphlans | 53 | -24.111±0.478 | **1.223**±0.024 | <0.0001 | <0.0001 | 0.981 |
| Afrotheria | 6 | -21.115±1.027 | **1.079**±0.051 | <0.0001 | <0.0001 | 0.991 |
| Artiodactyla | 5 | -26.632±2.588 | **1.351**±0.118 | 0.0020 | 0.0014 | 0.978 |
| Carnivora | 8 | -21.332±2.667 | **1.095**±0.126 | 0.0002 | 0.0001 | 0.927 |
| Chiroptera | 13 | -23.582±1.610 | **1.183**±0.089 | <0.0001 | <0.0001 | 0.941 |
| Eulipotyphla | 5 | -21.165±1.520 | **1.028**±0.084 | 0.0008 | 0.0012 | 0.980 |
| Marsupialia | 10 | -23.421±0.746 | **1.186**±0.037 | <0.0001 | <0.0001 | 0.992 |
| Primata | 13 | -19.572±0.794 | **0.976**±0.036 | <0.0001 | <0.0001 | 0.985 |
| Glires | 10 | -26.462±1.406 | **1.349**±0.073 | <0.0001 | <0.0001 | 0.977 |
| Scandentia | 1 |  |  |  |  |  |
| All non-primates, non-eulipotyphlans,<br>Accipitriformes,<br>Anseriformes,<br>Columbiformes,<br>Galliformes,<br>Palaeognathae,<br>Strigiformes | 84 | -24.283±0.403 | **1.234**±0.021 | <0.0001 | <0.0001 | 0.978 |

**Table 7.**
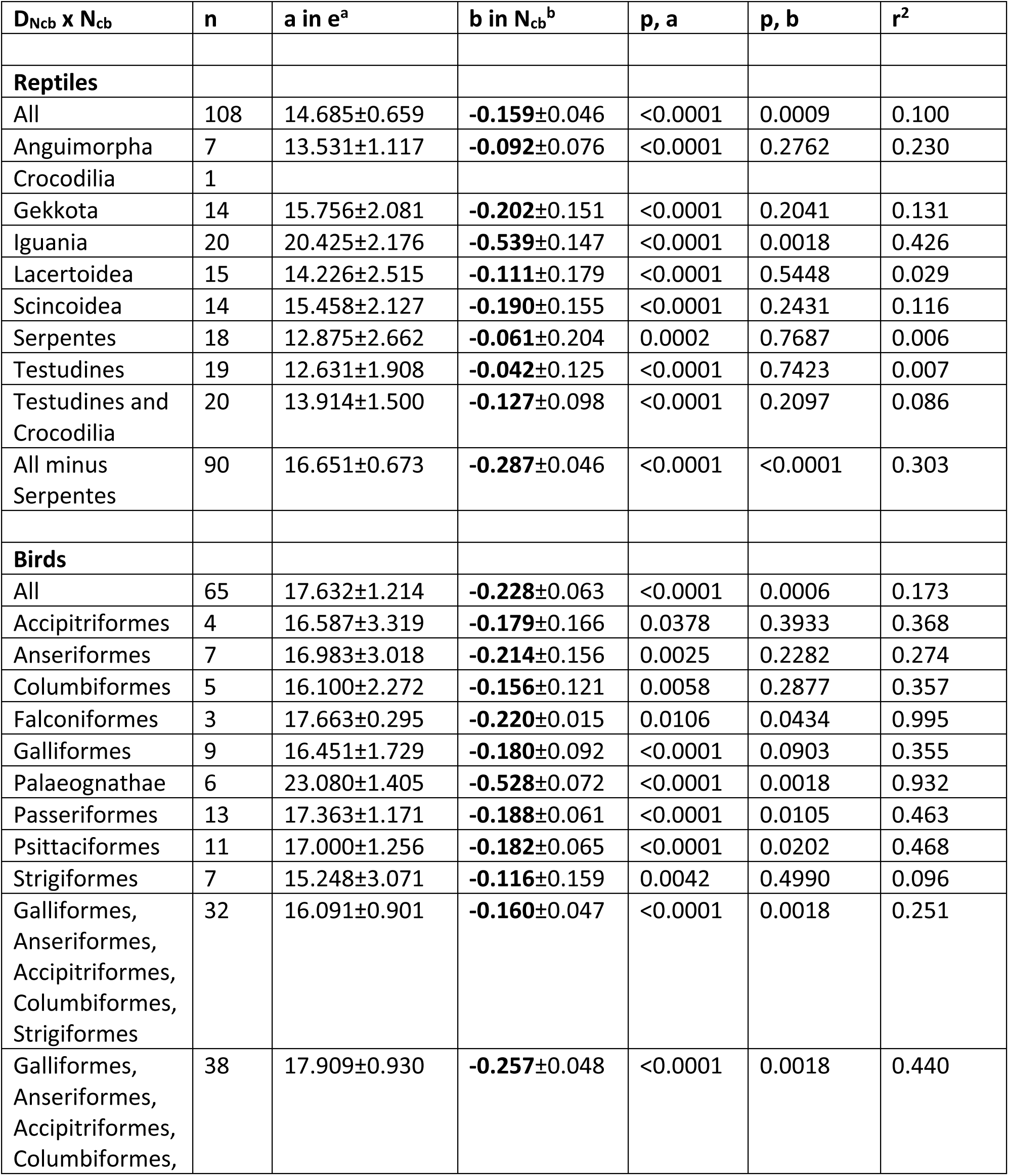
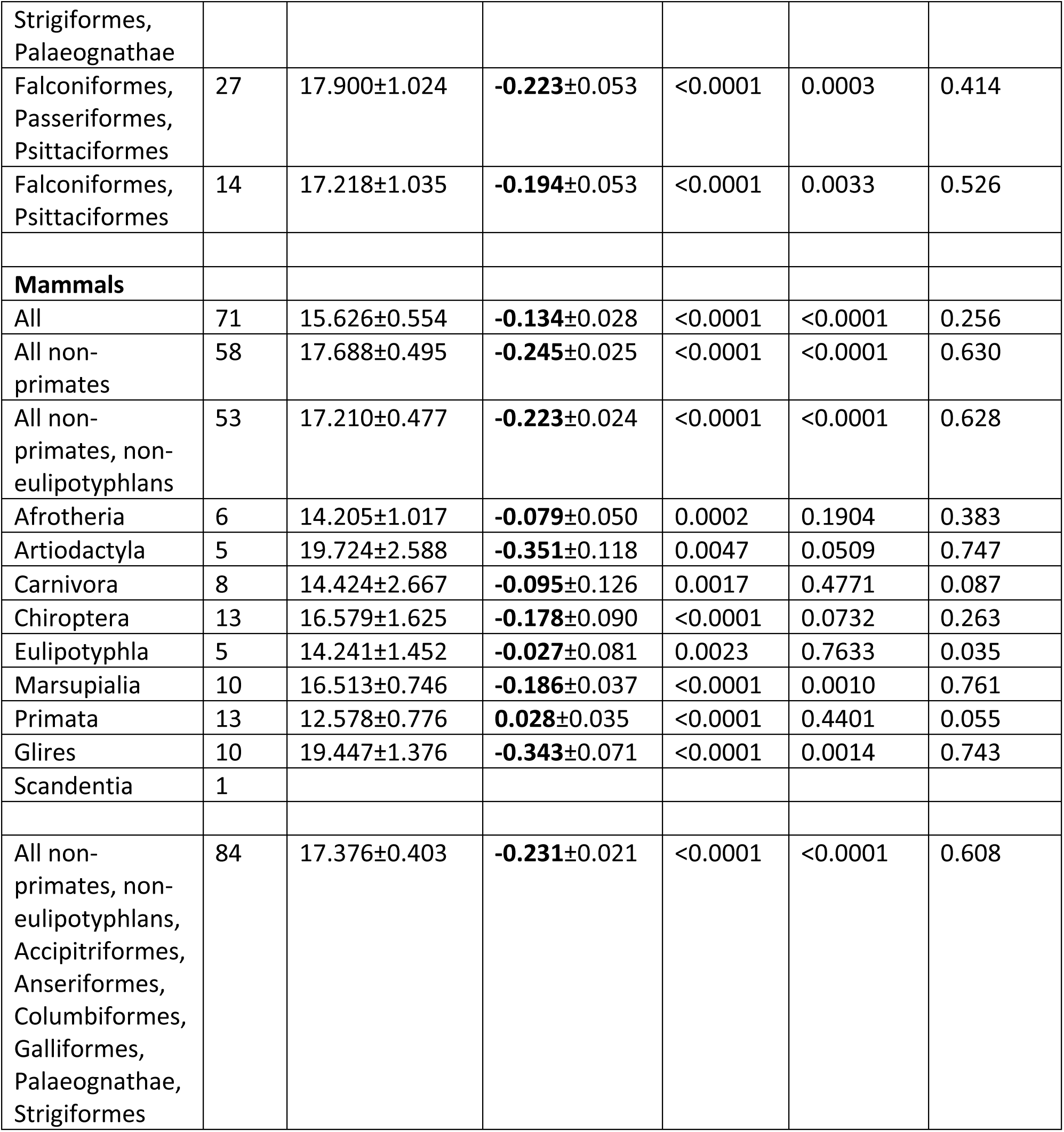
Scaling of neuronal density with varying numbers of neurons in the cerebellum of reptiles, birds and mammals. Values correspond to the equation DNcb = e^a±SE^ Ncb^b±SE^, the p-values for a and b, and the r^2^ for the power fit, where DNcb is neuronal density in the cerebellum and Ncb is the number of neurons in the cerebellum.

A power function with an exponent of 1.257±0.048, statistically indistinguishable from the reptilian exponent, applies to the cerebellum of early derived birds: Accipitriformes, Anseriformes, Columbiformes, Galliformes and Palaeognathae, and also to Strigiformes (Figure 13A, grey line), although this curve is shifted to the right with respect to reptiles, just as seen in the case of the telencephalon/pallium between Testudine and crocodilian reptiles and early-derived birds (Figure 12E). Remarkably, a similar power function to the one that applies to these six bird clades describes the scaling of the non-primate, non-eulipotyphlan mammalian cerebellum (Herculano-Houzel et al., 2007, 2014; Sarko et al., 2009; Jardim-Messeder et al., 2017), with a matching exponent of 1.223±0.024 (Figure 13A, red line; Figure 14A,B). Indeed, the scaling relationship that applies to the combination of these two groups of birds and mammals has an undistinguishable exponent of 1.234±0.021 (Figure 14A,B; Table 6). Additionally, the scaling exponent that applies to early-derived birds, plus Strigiformes, is virtually identical to the scaling exponent that applies to reptiles as a whole (1.160±0.047 and 1.159±0.046, respectively), with a distribution that is simply shifted to the right, with more neurons in a same structure mass (Figure 14C; Table 6). Across the other three bird clades (Falconiformes, Passeriformes including corvids, and Psittaciformes), yet another seemingly parallel power function relates cerebellar mass to number of neurons, with exponent 1.223±0.053 (Table 6), not significantly different from the others (Figure 13A, Figure 14). Strikingly, the neuronal scaling rules that apply to the cerebellum can be further distinguished across Falconiformes and Psittaciformes on the one hand, and Passeriformes on the other, with even higher neuronal densities in the cerebellum of the latter (Figure 14F).

The similar exponents that describe the scaling of cerebellar mass as it gains neurons across reptiles, birds, and most mammalian species stems from the similar slope of the decrease in cerebellar neuronal densities with increasing numbers of neurons across the three groups described above (Figure 13B, compare green, grey/red, and black lines), with exponents of around -0.25, with parallel power functions that apply to each clade (Figure 14F, Table 7). As described previously (Herculano-Houzel, 2014a), primates and eulipotyphlans are the only groups amongst mammals that clearly depart from these allometric relationships, with no significant change in neuronal density in the cerebellum as it gains neurons (Figure 13B), which translates into cerebellar mass that scales linearly with numbers of neurons (Figure 13A, 14B; Tables 6, 7).

Mapping these scaling relationships onto the phylogenetic tree of all species of amniotes in the dataset indicates that while there has been a major step increase in numbers of cerebellar neurons in the two groups of endotherms compared to reptiles, and apparently two further step increases at the origin of more recently derived birds, the allometric exponent that applies to the scaling of neuronal density with numbers of neurons in the cerebellum only diverged at the origin of primates and eulipotyphlans, amongst mammals; and at the origins of Serpentes, and Iguania amongst reptiles (Figure 7C). The significance of such step increases in the context of the evolution of endothermy is addressed separately (Herculano-Houzel, 2025).

### Neuronal scaling rules for the rest of brain are shared between reptiles and non-primate mammals

Similarly to the telencephalon, the neuronal scaling rules that apply to the rest of brain (brainstem, diencephalon and subpallium in mammals, or brainstem plus diencephalon in reptiles and birds; Kverkova et al., 2022) overlap across most reptiles and mammals (Figure 15A, Table 8), with the exception of primates, which diverge from other mammals as reported previously (Herculano-Houzel et al., 2014a). As shown in Figures 15A and 16A, the 95% prediction interval of the scaling relationship for reptiles between the mass of the rest of brain and its number of neurons includes the majority of non-primate mammals, including large artiodactyls and carnivorans, with scaling exponents that are not significantly different (non-primate mammals, 1.844±0.076; reptiles, 1.569±0.067, excluding Testudines and crocodile; combined sample, 1.673±0.035; Table 6). Increases in numbers of neurons in the rest of brain are thus accompanied by decreasing neuronal densities with a scaling relationship across reptiles (excluding Testudines and crocodile) that includes most mammalian species (Figure 16B, Table 8). In contrast, and as found for the telencephalon, the rest of brain of Testudines and the Nile crocodile is larger in mass than in other reptilian species with similar numbers of neurons in the structure (Figure 16, Table 8).

**Figure 15.**
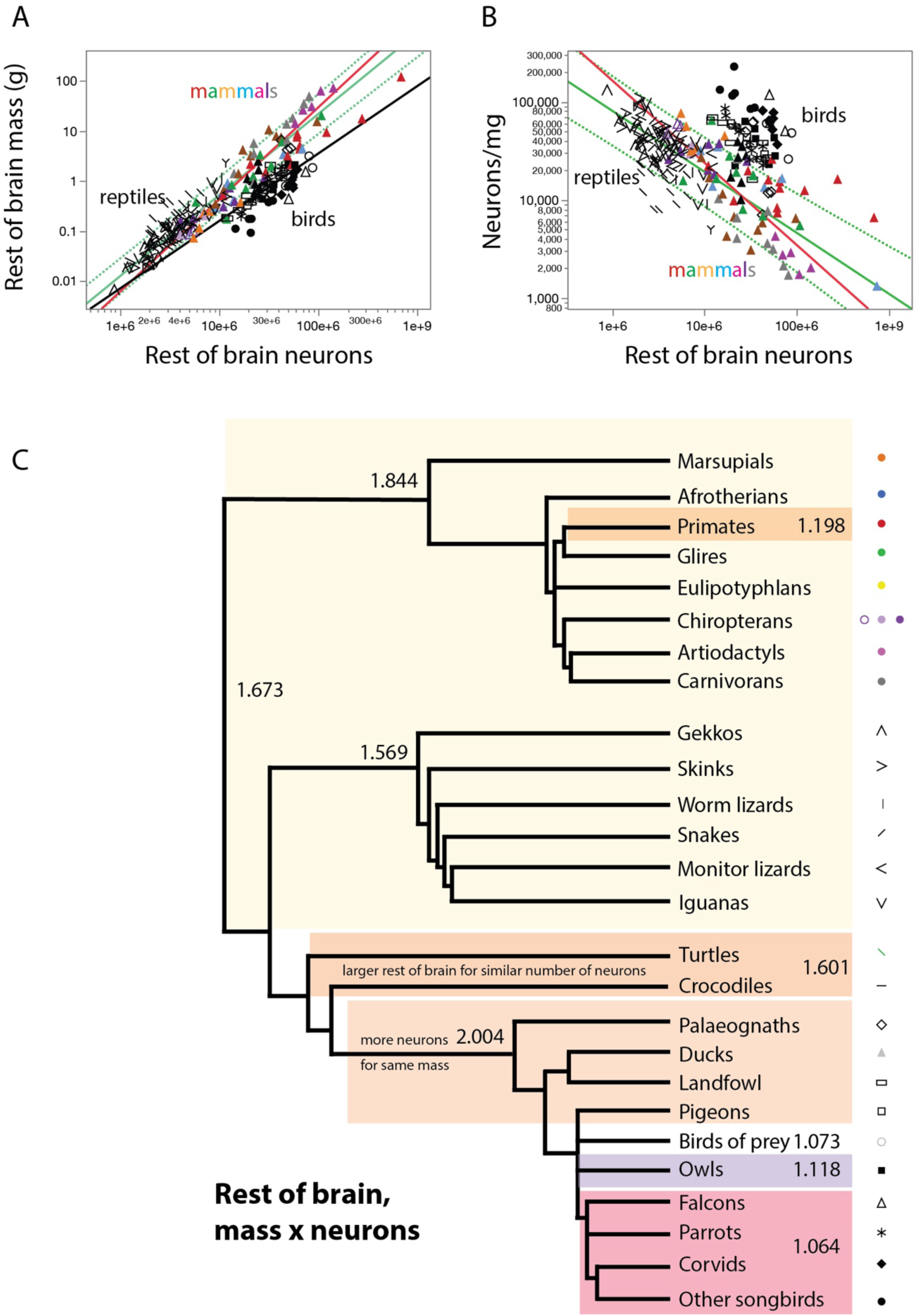
Reptiles and non-primate mammals share allometric exponents for the neuronal composition of the rest of brain, while the scaling relationship for birds is shifted to the right. **A**, scaling of rest of brain mass with numbers of neurons plotted for non-primate mammals (in red, exponent 1.844±0.076), all reptiles minus Testudines and crocodile (in green, exponent 1.569±0.067), and all birds (in black, exponent, 1.346±0.173). For the sake of clarity, the 95% prediction interval is only plotted for reptilian species. All power functions are listed in Table 8. **B**, Neuronal density in the rest of brain decreases steeply with increasing numbers of rest of brain neurons across reptilian and non-primate mammalian species, while birds have higher neuronal densities. Power functions plotted have exponents -0.573±0.067 (reptiles minus Testudines and crocodile, in green) and -0.845±0.076 (non-primate mammals, in red). Power functions for birds, listed in Table 9, are clade-specific and are not plotted for the sake of clarity. **C**, Putative cladogram for the evolution of the exponent that describes the scaling relationship between rest of brain mass and number of neurons, showing a shared exponent across most extant reptilian and mammalian species that can be inferred to have applied to ancestral amniotic species. Colored areas indicate clade-specific changes. Each data point corresponds to one species. The key used to depict each clade is shown to the right.

**Figure 16.**
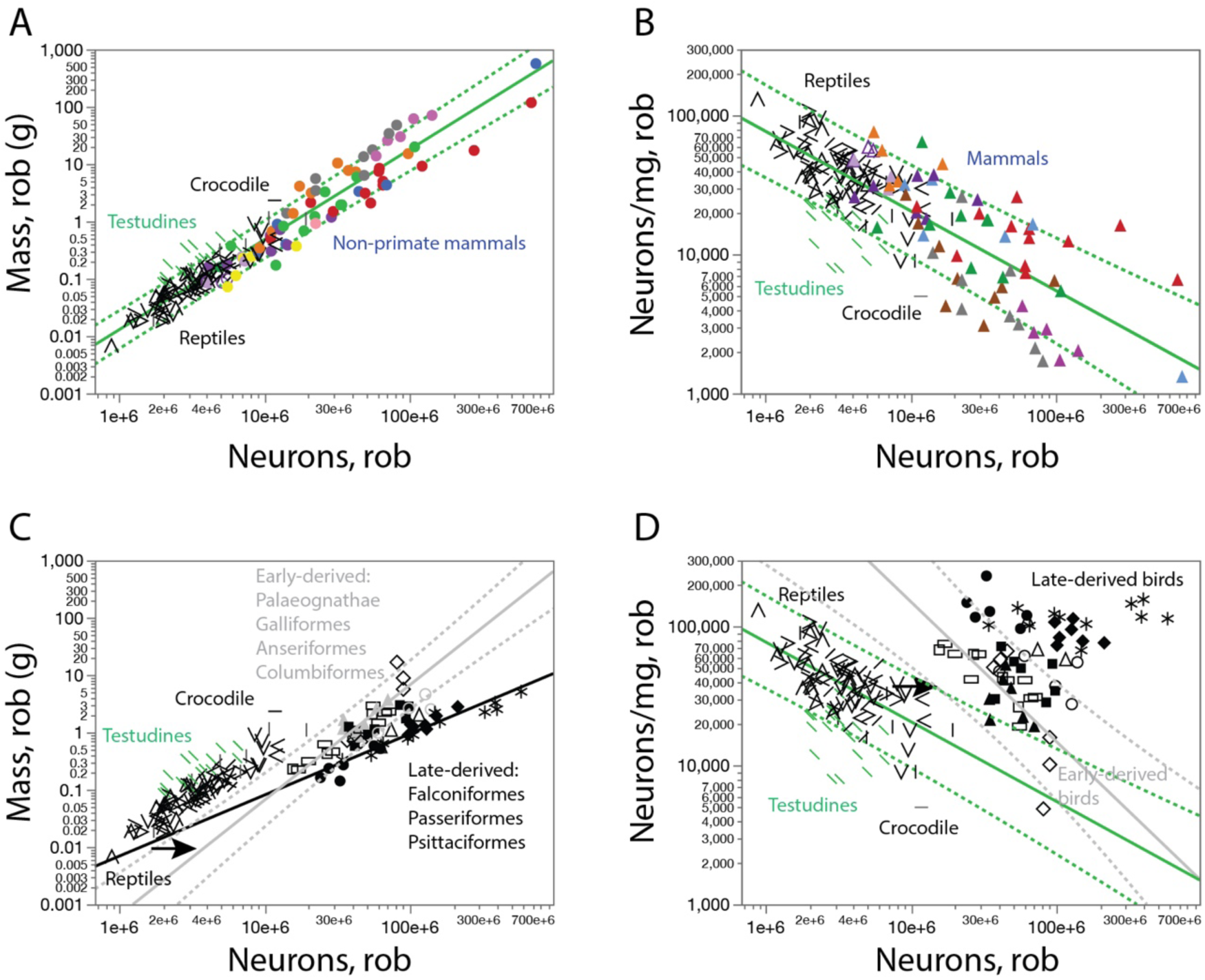
Neuronal scaling rules for the rest of brain of reptiles, mammals and birds. **A, C,** scaling of rest of brain mass with numbers of rest of brain neurons. **B, D,** scaling of neuronal density in the rest of brain with numbers of rest of brain neurons. **A, B,** The scaling rule that applies to Reptiles (minus Testudines and crocodile; plotted in green) includes most mammalian species in the dataset, in a continuum. **C, D,** parallel neuronal scaling rules with similar exponents apply to the rest of brain of reptiles and the early-derived bird clades. All power functions are listed in Tables 8 and 9.

**Table 8.**
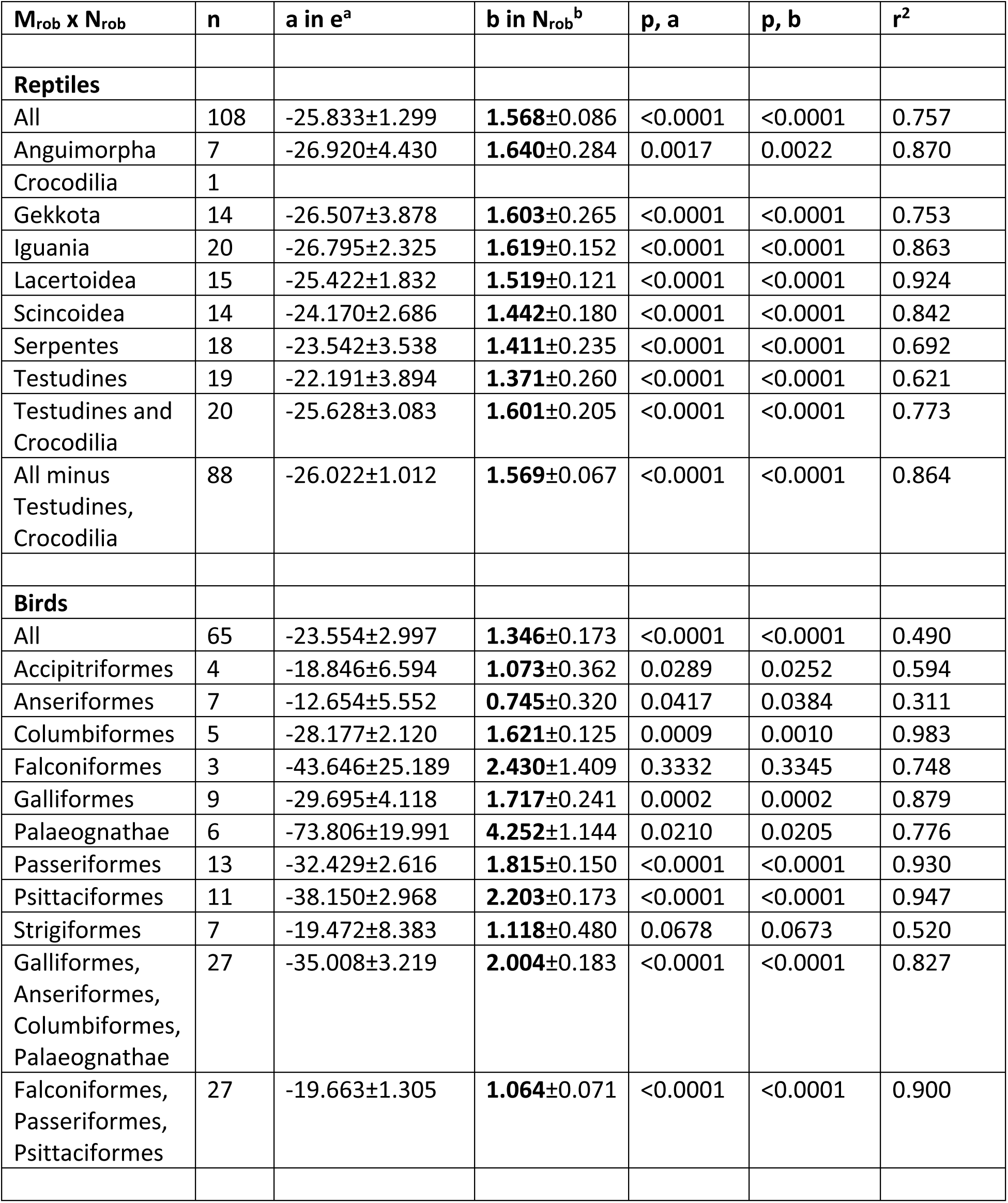
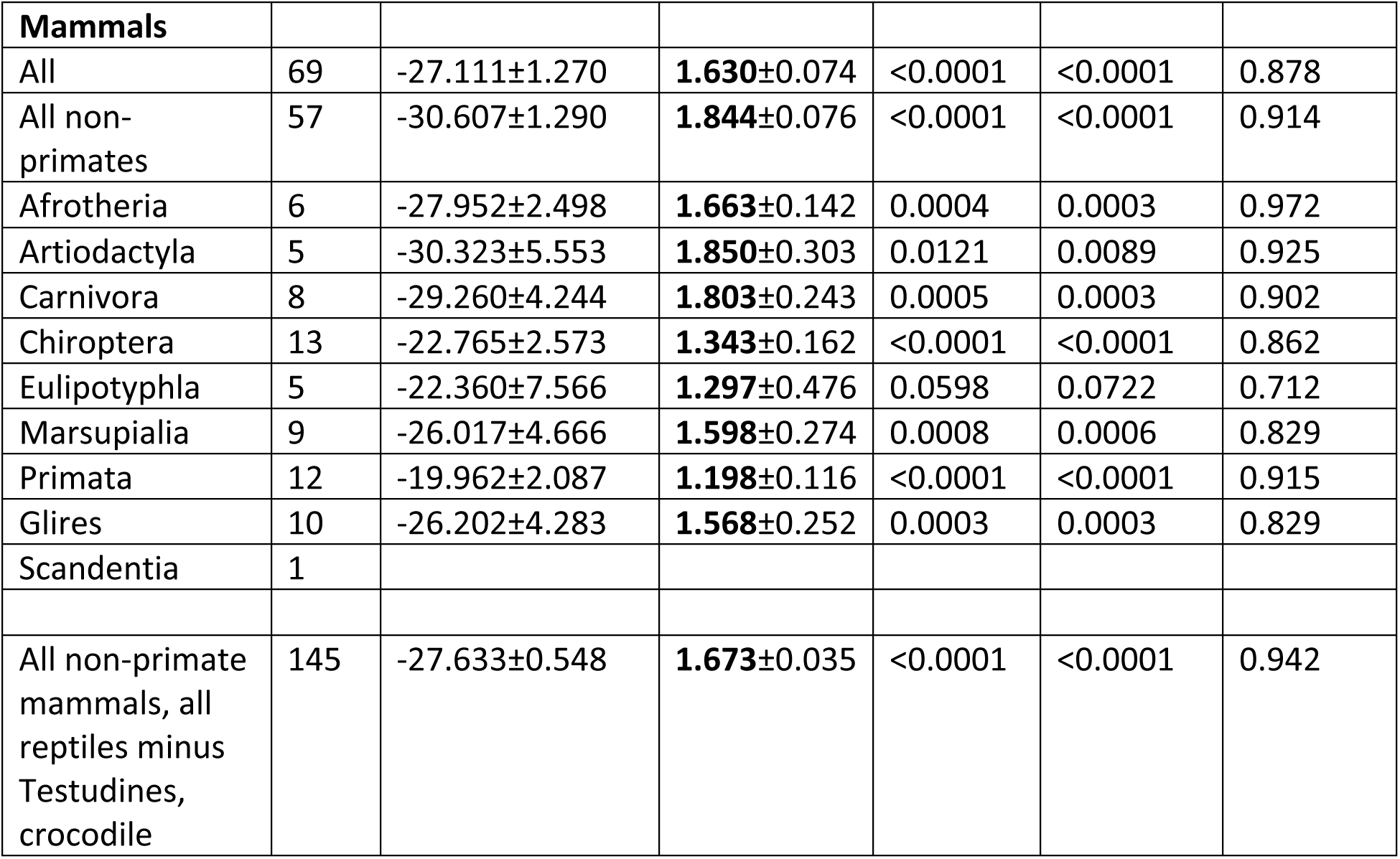
Neuronal scaling rules for the rest of brain of reptiles, birds and mammals. Values correspond to the equation Mrob = e^a±SE^ Nrob^b±SE^, the p-values for a and b, and the r^2^ for the power fit, where Mrob is mass of the rest of brain and Nrob is the number of neurons in the rest of brain. In reptiles and birds, rest of brain consists of the brainstem and diencephalon; in mammals, it additionally contains the striatum.

**Table 9.** Scaling of neuronal density with varying numbers of neurons in the rest of brain of reptiles, birds and mammals. Values correspond to the equation DNrob = e^a±SE^ Nrob^b±SE^, the p-values for a and b, and the r^2^ for the power fit, where DNrob is neuronal density in the rest of brain and Nrob is the number of neurons in the rest of brain. In reptiles and birds, rest of brain consists of the brainstem and diencephalon; in mammals, it additionally contains the striatum.

| $D_{N_{rob}} \times N_{rob}$ | n | a in $e^a$ | b in $N_{rob}^b$ | p, a | p, b | $r^2$ |
| --- | --- | --- | --- | --- | --- | --- |
| **Reptiles** |  |  |  |  |  |  |
| All | 108 | 18.988±1.306 | **-0.572±0.087** | <0.0001 | <0.0001 | 0.291 |
| Anguimorpha | 7 | 20.0455±4.455 | **-0.641±0.285** | 0.0064 | 0.0744 | 0.503 |
| Crocodylia | 1 |  |  |  |  |  |
| Gekkota | 14 | 19.654±3.864 | **-0.605±0.264** | 0.0003 | 0.0409 | 0.304 |
| Iguania | 20 | 19.971±2.353 | **-0.625±0.154** | <0.0001 | 0.0007 | 0.479 |
| Lacertoidea | 15 | 18.613±1.844 | **-0.526±0.122** | <0.0001 | 0.0008 | 0.589 |
| Scincoidea | 14 | 17.280±2.684 | **-0.443±0.180** | <0.0001 | 0.0299 | 0.336 |
| Serpentes | 18 | 16.789±3.585 | **-0.421±0.238** | 0.0002 | 0.0962 | 0.163 |
| Testudines | 19 | 15.473±3.996 | **-0.384±0.267** | 0.0012 | 0.1683 | 0.109 |
| Testudines and Crocodylia | 20 | 18.854±3.151 | **-0.610±0.209** | <0.0001 | 0.0092 | 0.321 |
| All minus Testudines, Crocodylia | 90 | 19.168±1.017 | **-0.573±0.067** | <0.0001 | <0.0001 | 0.456 |
| **Birds** |  |  |  |  |  |  |
| All | 65 | 16.611±2.991 | **-0.344±0.173** | <0.0001 | 0.0505 | 0.059 |
| Accipitriformes | 4 | 11.919±6.627 | **-0.072±0.364** | 0.1222 | 0.8489 | 0.007 |
| Anseriformes | 7 | 5.491±5.602 | **0.269±0.323** | 0.3463 | 0.4210 | 0.055 |
| Columbiformes | 5 | 21.172±2.138 | **-0.615±0.126** | 0.0022 | 0.0163 | 0.889 |
| Falconiformes | 3 | 36.340±25.388 | **-1.407±1.420** | 0.3882 | 0.5027 | 0.496 |
| Galliformes | 9 | 22.759±4.091 | **-0.715±0.239** | 0.0008 | 0.0202 | 0.561 |
| Palaeognathae | 6 | 66.530±19.270 | **-3.230±1.102** | 0.0260 | 0.0428 | 0.682 |
| Passeriformes | 13 | 25.522±2.616 | **-0.815±0.150** | <0.0001 | 0.0002 | 0.728 |
| Psittaciformes | 11 | 31.242±2.968 | **-1.203±0.173** | <0.0001 | <0.0001 | 0.843 |
| Strigiformes | 7 | 12.655±8.411 | **-0.123±0.481** | 0.1928 | 0.8084 | 0.013 |
| Galliformes,<br>Columbiformes,<br>Palaeognathae | 32 | 27.464±4.190 | **-0.990±0.244** | <0.0001 | 0.0007 | 0.479 |
| Accipitriformes,<br>Falconiformes,<br>Passeriformes | 20 | 25.948±2.948 | **-0.842±0.168** | <0.0001 | <0.0001 | 0.583 |
| **Mammals** |  |  |  |  |  |  |
| All | 69 | 20.163±1.272 | **-0.627±0.074** | <0.0001 | <0.0001 | 0.515 |
| All non-primates | 57 | 23.707±1.290 | **-0.845±0.076** | <0.0001 | <0.0001 | 0.690 |
| Afrotheria | 6 | 21.029±2.519 | **-0.662±0.144** | 0.0011 | 0.0099 | 0.842 |
| Artiodactyla | 5 | 23.412±5.552 | **-0.850±0.303** | 0.0244 | 0.0677 | 0.724 |
| Carnivora | 8 | 22.353±4.245 | **-0.803±0.243** | 0.0019 | 0.0163 | 0.646 |
| Chiroptera | 13 | 15.823±2.532 | **-0.340±0.160** | <0.0001 | 0.0564 | 0.292 |
| Eulipotyphla | 5 | 15.344±7.440 | **-0.290±0.468** | 0.1312 | 0.5789 | 0.114 |
| Marsupialia | 9 | 19.109±4.666 | **-0.598±0.274** | 0.0046 | 0.0654 | 0.405 |
| Primata | 12 | 12.734±2.035 | **-0.180±0.113** | <0.0001 | 0.1413 | 0.203 |
| Glires | 10 | 19.215±4.232 | **-0.563±0.249** | 0.0019 | 0.0535 | 0.390 |
| Scandentia | 1 |  |  |  |  |  |
| All non-primate<br>mammals, all<br>reptiles minus<br>Testudines,<br>crocodile | 145 | 20.717±0.549 | **-0.672±0.035** | <0.0001 | <0.0001 | 0.724 |

The conformity of most reptilian and mammalian species to the same scaling relationship between numbers of neurons in the rest of brain and average neuronal density in these structures, with continuity across these species in scaling (Figure 15), indicates that the neuronal scaling rule that they share also applied to their common ancestral amniote species (Figure 15C). Thus, as for the telencephalon, what differentiates the rest of brain of mammals from the majority of reptiles seems to be simply the vastly larger numbers of neurons in the former, but apparently added in conformity to the same scaling rules.

Birds, in contrast, have more neurons in the rest of brain than both reptiles and mammals with similar mass in these structures (Figure 15B, Figure 16C). The scaling relationship that applies to the rest of brain of the early-derived birds might also be simply shifted to the right compared to reptiles (as found for the cerebellum), with an exponent of 2.004±0.183 across Galliformes, Anseriformes, Columbiformes and Palaeognathae that is statistically indistinguishable from the exponent of 1.601±0.205 that applies to Testudines and crocodile (Figure 16C). Amongst birds, however, different relationships between numbers of neurons and neuronal density apply to the early-derived Palaeognathae, Galliformes, Anseriformes and Collumbiformes versus the later-derived Falconiformes, Passeriformes and Psittaciformes (Tables 8 and 9; Figure 16D).

### Relationships of numbers of neurons to body mass

In contrast to the marked continuity in the relationship between structure mass and numbers of neurons across amniote clades pointed out above, the numbers of neurons that compose the telencephalon/pallium (Figure 17A), cerebellum (Figure 17B) and rest of brain (Figure 17C) are not universally related to body mass across all amniotes. Across reptiles, while the telencephalon of all species conforms to the same scaling relationship between numbers of neurons and body mass (Figure 17A), Serpentes deviate markedly from all other reptilian clades in their much shallower scaling of numbers of neurons in the cerebellum with body mass (Figure 17B), and Serpentes, Testudines and Crocodilians together differ from other reptilian clades in how numbers of neurons in the rest of brain scales with body mass (Figure 17C; Table 10).

**Figure 17.**
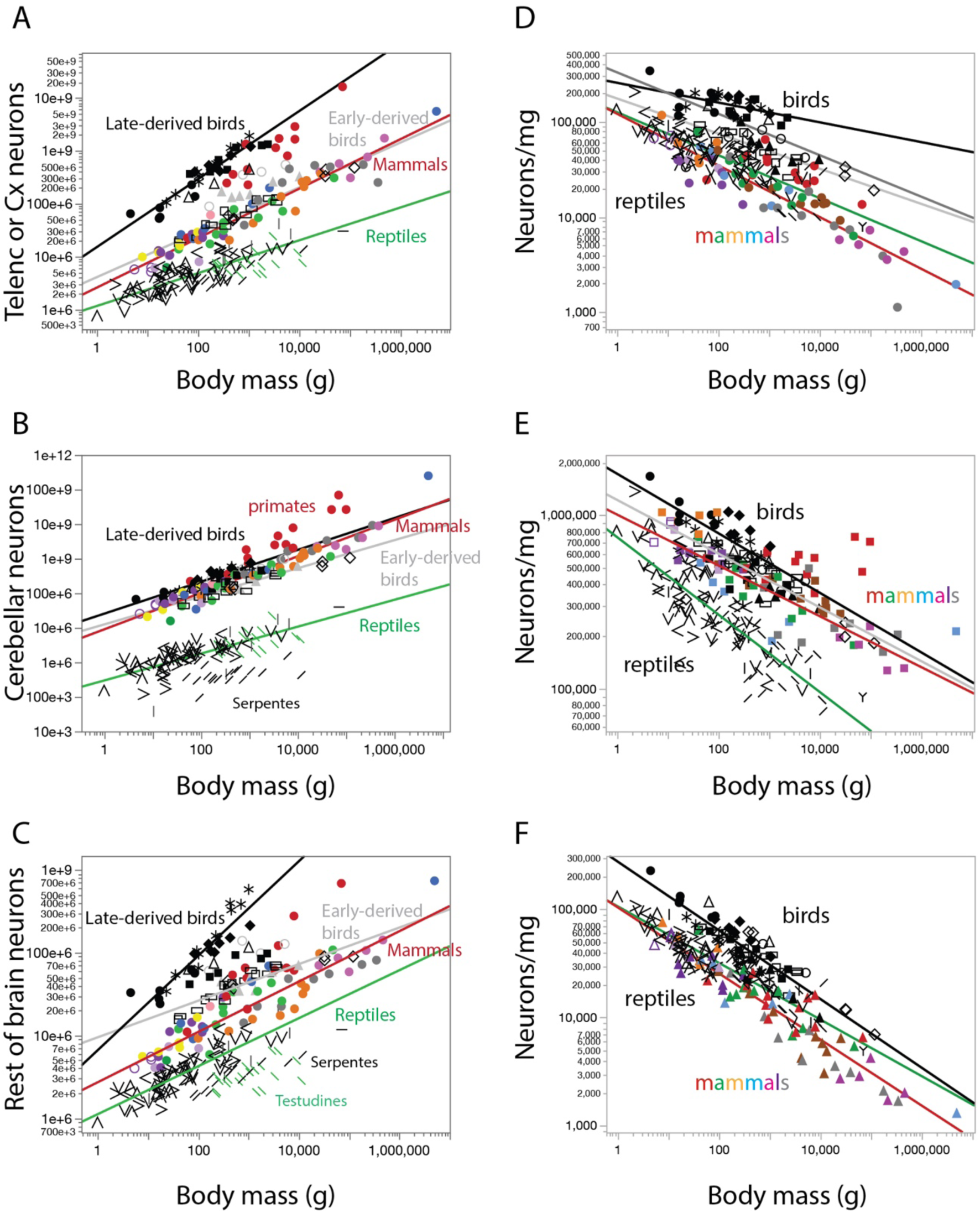
Clade-specific scaling of numbers of neurons and neuronal density in the different brain structures with body mass across mammals, reptiles and birds. A-C, scaling of numbers of neurons in the telencephalon or pallium (A), cerebellum (B) and rest of brain (C) with body mass. D-F, scaling of neuronal density in the telencephalon or pallium (D), cerebellum (E) and rest of brain (F) with body mass. Power functions plotted apply to reptiles (green, minus Testudines and crocodile in **A**, minus Serpentes in **B**, and minus Serpentes, Testudines and crocodile in **C**), non-primate mammals (red), early-derived birds (light gray: Palaeognathae, Galliformes and Columbiformes in **A**; Anseriformes, Palaeognathae, Galliformes and Columbiformes in **B** and **C**), and late-derived birds (black: Passeriformes, Psittaciformes and Strigiformes in **A**, Accipitriformes, Falconiformes, Passeriformes and Psittaciformes in **B**, and Passeriformes and Psittaciformes in **C**). Power functions are listed in Table 10.

**Table 10.** Scaling of numbers of neurons in the different brain structures of reptiles, birds and mammals with varying body mass. Values correspond to the equation Nstr = e^a±SE^ Mbd ^b±SE^, the p-values for a and b, and the r^2^ for the power fit, where Nstr is the number of neurons in a given brain structure and Mbd is the average body mass for the species.

| $N_{str} \times M_{bd}$ | n | a in $e^a$ | b in $M_{bd}^b$ | p, a | p, b | $r^2$ |
| --- | --- | --- | --- | --- | --- | --- |
| **Cortex/pallium/telencephalon** |  |  |  |  |  |  |
| **Reptiles** |  |  |  |  |  |  |
| All | 108 | 14.063±0.107 | **0.278**±0.020 | <0.0001 | <0.0001 | 0.649 |
| Anguimorpha | 7 | 13.561±0.406 | **0.444**±0.062 | <0.0001 | 0.0008 | 0.912 |
| Crocodylia | 1 |  |  |  |  |  |
| Gekkota | 14 | 13.803±0.276 | **0.414**±0.093 | <0.0001 | 0.0008 | 0.623 |
| Iguania | 20 | 13.531±0.148 | **0.339**±0.030 | <0.0001 | <0.0001 | 0.880 |
| Lacertoidea | 15 | 13.866±0.219 | **0.412**±0.055 | <0.0001 | <0.0001 | 0.811 |
| Scincoidea | 14 | 13.942±0.252 | **0.390**±0.066 | <0.0001 | <0.0001 | 0.743 |
| Serpentes | 18 | 13.780±0.328 | **0.345**±0.054 | <0.0001 | <0.0001 | 0.719 |
| Testudines | 19 | 14.752±0.551 | **0.154**±0.078 | <0.0001 | 0.0639 | 0.188 |
| Testudines and Crocodylia | 20 | 14.298±0.460 | **0.222**±0.063 | <0.0001 | 0.0023 | 0.412 |
| All minus Testudines, Crocodylia | 88 | 13.956±0.116 | **0.311**±0.024 | <0.0001 | <0.0001 | 0.658 |
| All minus Serpentes | 90 | 14.083±0.110 | **0.286**±0.021 | <0.0001 | <0.0001 | 0.676 |
| **Birds** |  |  |  |  |  |  |
| All | 65 | 17.803±0.452 | **0.225**±0.073 | <0.0001 | 0.0029 | 0.133 |
| Accipitriformes | 4 | 15.879±0.947 | **0.518**±0.134 | 0.0035 | 0.0611 | 0.882 |
| Anseriformes | 7 | 17.231±0.673 | **0.243**±0.094 | <0.0001 | 0.0487 | 0.573 |
| Columbiformes | 5 | 14.937±0.480 | **0.503**±0.085 | <0.0001 | 0.0095 | 0.922 |
| Falconiformes | 3 | 16.913±0.183 | **0.437**±0.033 | 0.0069 | 0.0478 | 0.994 |
| Galliformes | 9 | 15.097±0.210 | **0.432**±0.032 | <0.0001 | <0.0001 | 0.963 |
| Palaeognathae | 6 | 15.194±0.304 | **0.427**±0.035 | <0.0001 | 0.0002 | 0.975 |
| Passeriformes | 13 | 16.398±0.331 | **0.657**±0.071 | <0.0001 | <0.0001 | 0.885 |
| Psittaciformes | 11 | 16.139±0.200 | **0.748±0.038** | <0.0001 | <0.0001 | 0.978 |
| Strigiformes | 7 | 17.920±0.566 | **0.405±0.092** | <0.0001 | 0.0070 | 0.795 |
| Passeriformes,<br>Psittaciformes,<br>Strigiformes | 31 | 16.534±0.217 | **0.645±0.041** | <0.0001 | <0.0001 | 0.893 |
| Galliformes,<br>Columbiformes,<br>Palaeognathae | 20 | 15.217±0.147 | **0.425±0.021** | <0.0001 | <0.0001 | 0.959 |
| Accipitriformes,<br>Falconiformes | 7 | 16.941±0.611 | **0.390±0.095** | <0.0001 | 0.0092 | 0.773 |
| **Mammals** |  |  |  |  |  |  |
| All | 70 | 14.772±0.283 | **0.519±0.038** | <0.0001 | <0.0001 | 0.737 |
| All non-primates | 58 | 14.728±0.141 | **0.472±0.019** | <0.0001 | <0.0001 | 0.917 |
| Afrotheria | 6 | 15.046±0.187 | **0.484±0.023** | <0.0001 | <0.0001 | 0.991 |
| Artiodactyla | 5 | 14.815±2.420 | **0.468±0.208** | 0.0088 | 0.1097 | 0.628 |
| Carnivora | 8 | 16.617±1.258 | **0.287±0.129** | <0.0001 | 0.0673 | 0.453 |
| Chiroptera | 13 | 14.787±0.344 | **0.417±0.093** | <0.0001 | 0.0009 | 0.647 |
| Eulipotyphla | 5 | 15.175±0.221 | **0.410±0.063** | <0.0001 | 0.0075 | 0.933 |
| Marsupialia | 9 | 13.797±0.671 | **0.565±0.080** | <0.0001 | 0.0002 | 0.876 |
| Primata | 12 | 14.195±0.755 | **0.825±0.097** | <0.0001 | <0.0001 | 0.878 |
| Glires | 11 | 14.556±0.344 | **0.469±0.052** | <0.0001 | <0.0001 | 0.902 |
| Scandentia | 1 |  |  |  |  |  |
| **Cerebellum** |  |  |  |  |  |  |
| **Reptiles** |  |  |  |  |  |  |
| All | 108 | 12.701±0.228 | **0.305±0.042** | <0.0001 | <0.0001 | 0.328 |
| Anguimorpha | 7 | 8.846±1.324 | **0.925±0.201** | 0.0011 | 0.0059 | 0.808 |
| Crocodylia | 1 |  |  |  |  |  |
| Gekkota | 14 | 12.240±0.298 | **0.567±0.100** | <0.0001 | 0.0001 | 0.728 |
| Iguania | 20 | 13.494±0.189 | **0.273±0.038** | <0.0001 | <0.0001 | 0.744 |
| Lacertoidea | 15 | 12.673±0.354 | **0.370±0.089** | <0.0001 | 0.0012 | 0.568 |
| Scincoidea | 14 | 12.259±0.556 | **0.402±0.146** | <0.0001 | 0.0174 | 0.387 |
| Serpentes | 18 | 12.156±0.573 | **0.144±0.094** | <0.0001 | 0.1457 | 0.128 |
| Testudines | 19 | 13.501±0.815 | **0.245±0.115** | <0.0001 | 0.0479 | 0.211 |
| Testudines and<br>Crocodylia | 20 | 12.728±0.691 | **0.361±0.063** | <0.0001 | 0.0012 | 0.451 |
| All minus<br>Testudines,<br>Crocodylia,<br>Serpentes | 70 | 12.534±0.197 | **0.413±0.044** | <0.0001 | <0.0001 | 0.559 |
| All minus<br>Serpentes | 90 | 12.607±0.168 | **0.389**±0.032 | <0.0001 | <0.0001 | 0.625 |
| **Birds** |  |  |  |  |  |  |
| All | 65 | 17.441±0.212 | **0.312**±0.034 | <0.0001 | <0.0001 | 0.571 |
| Accipitriformes | 4 | 17.497±0.455 | **0.367**±0.065 | 0.0007 | 0.0296 | 0.942 |
| Anseriformes | 7 | 16.545±0.610 | **0.398**±0.085 | <0.0001 | 0.0054 | 0.814 |
| Columbiformes | 5 | 15.502±0.539 | **0.585**±0.095 | <0.0001 | 0.0086 | 0.927 |
| Falconiformes | 3 | 16.766±0.226 | **0.513**±0.041 | 0.0086 | 0.0502 | 0.994 |
| Galliformes | 9 | 15.698±0.384 | **0.494**±0.059 | <0.0001 | <0.0001 | 0.910 |
| Palaeognathae | 6 | 16.455±0.182 | **0.374**±0.021 | <0.0001 | <0.0001 | 0.988 |
| Passeriformes | 13 | 16.882±0.209 | **0.524**±0.045 | <0.0001 | <0.0001 | 0.925 |
| Psittaciformes | 11 | 16.620±0.306 | **0.540**±0.058 | <0.0001 | <0.0001 | 0.907 |
| Strigiformes | 7 | 16.942±0.380 | **0.399**±0.062 | <0.0001 | 0.0013 | 0.893 |
| Accipitriformes,<br>Falconiformes,<br>Passeriformes,<br>Psittaciformes | 21 | 15.933±0.276 | **0.477**±0.042 | <0.0001 | <0.0001 | 0.872 |
| Anseriformes,<br>Galliformes,<br>Columbiformes,<br>Palaeognathae | 27 | 16.343±0.203 | **0.406**±0.028 | <0.0001 | <0.0001 | 0.891 |
| **Mammals** |  |  |  |  |  |  |
| All | 72 | 15.871±0.223 | **0.585**±0.029 | <0.0001 | <0.0001 | 0.854 |
| All non-primates | 59 | 15.953±0.145 | **0.535**±0.019 | <0.0001 | <0.0001 | 0.931 |
| Afrotheria | 6 | 14.729±0.485 | **0.733**±0.059 | <0.0001 | 0.0002 | 0.975 |
| Artiodactyla | 5 | 16.527±2.072 | **0.463**±0.178 | 0.0041 | 0.0801 | 0.693 |
| Carnivora | 8 | 16.179±0.551 | **0.522**±0.056 | <0.0001 | <0.0001 | 0.935 |
| Chiroptera | 13 | 16.496±0.252 | **0.448**±0.068 | <0.0001 | <0.0001 | 0.797 |
| Eulipotyphla | 5 | 15.049±0.306 | **0.873**±0.088 | <0.0001 | 0.0022 | 0.970 |
| Marsupialia | 9 | 15.422±0.317 | **0.577**±0.037 | <0.0001 | <0.0001 | 0.967 |
| Primata | 12 | 15.656±0.622 | **0.754**±0.073 | <0.0001 | <0.0001 | 0.906 |
| Glires | 11 | 15.583±0.459 | **0.540**±0.069 | <0.0001 | <0.0001 | 0.872 |
| Scandentia | 1 |  |  |  |  |  |
| **Rest of brain** |  |  |  |  |  |  |
| **Reptiles** |  |  |  |  |  |  |
| All | 108 | 14.242±0.101 | **0.168**±0.019 | <0.0001 | <0.0001 | 0.429 |
| Anguimorpha | 7 | 13.564±0.555 | **0.325**±0.084 | <0.0001 | 0.0120 | 0.748 |
| Crocodylia | 1 |  |  |  |  |  |
| Gekkota | 14 | 13.806±0.177 | **0.292**±0.060 | <0.0001 | 0.0004 | 0.667 |
| Iguania | 20 | 14.086±0.165 | **0.262±0.033** | <0.0001 | <0.0001 | 0.781 |
| Lacertoidea | 15 | 14.003±0.165 | **0.317±0.042** | <0.0001 | <0.0001 | 0.816 |
| Scincoidea | 14 | 13.791±0.252 | **0.313±0.066** | <0.0001 | 0.0005 | 0.651 |
| Serpentes | 18 | 14.192±0.306 | **0.142±0.050** | <0.0001 | 0.0120 | 0.334 |
| Testudines | 19 | 13.927±0.430 | **0.152±0.061** | <0.0001 | 0.0223 | 0.271 |
| Testudines and Crocodilia | 20 | 13.529±0.363 | **0.212±0.049** | <0.0001 | 0.0004 | 0.506 |
| Testudines, Crocodilia, Serpentes | 38 | 14.035±0.226 | **0.153±0.033** | <0.0001 | <0.0001 | 0.370 |
| All minus Testudines, Crocodilia, Serpentes | 70 | 13.924±0.085 | **0.290±0.019** | <0.0001 | <0.0001 | 0.770 |
| **Birds** |  |  |  |  |  |  |
| All | 65 | 17.185±0.290 | **0.140±0.047** | <0.0001 | 0.0038 | 0.125 |
| Accipitriformes | 4 | 17.123±0.869 | **0.189±0.123** | 0.0026 | 0.2646 | 0.541 |
| Anseriformes | 7 | 16.543±0.481 | **0.157±0.067** | <0.0001 | 0.0668 | 0.522 |
| Columbiformes | 5 | 15.296±0.222 | **0.353±0.039** | <0.0001 | 0.0026 | 0.967 |
| Falconiformes | 3 | 16.994±0.215 | **0.224±0.038** | 0.0080 | 0.1083 | 0.971 |
| Galliformes | 9 | 15.374±0.202 | **0.329±0.031** | <0.0001 | <0.0001 | 0.942 |
| Palaeognathae | 6 | 16.599±0.176 | **0.154±0.020** | <0.0001 | 0.0015 | 0.937 |
| Passeriformes | 13 | 16.180±0.224 | **0.434±0.048** | <0.0001 | <0.0001 | 0.881 |
| Psittaciformes | 11 | 14.941±0.298 | **0.749±0.056** | <0.0001 | <0.0001 | 0.952 |
| Strigiformes | 7 | 16.449±0.521 | **0.247±0.085** | <0.0001 | 0.0328 | 0.631 |
| Passeriformes, Psittaciformes | 24 | 15.707±0.251 | **0.571±0.051** | <0.0001 | <0.0001 | 0.853 |
| Anseriformes, Galliformes, Columbiformes, Palaeognathae | 27 | 16.065±0.145 | **0.221±0.020** | <0.0001 | <0.0001 | 0.826 |
| Accipitriformes, Falconiformes | 7 | 17.158±0.356 | **0.188±0.055** | <0.0001 | 0.0191 | 0.699 |
| **Mammals** |  |  |  |  |  |  |
| All | 70 | 14.785±0.162 | **0.330±0.022** | <0.0001 | <0.0001 | 0.775 |
| All non-primates | 58 | 14.793±0.119 | **0.307±0.016** | <0.0001 | <0.0001 | 0.869 |
| Afrotheria | 6 | 14.771±0.294 | **0.375±0.036** | <0.0001 | 0.0005 | 0.965 |
| Artiodactyla | 5 | 15.569±1.320 | **0.235±0.113** | 0.0013 | 0.1294 | 0.590 |
| Carnivora | 8 | 14.909±0.412 | **0.269±0.042** | <0.0001 | 0.0007 | 0.872 |
| Chiroptera | 13 | 14.343±0.271 | **0.431±0.073** | <0.0001 | 0.0001 | 0.759 |
| Eulipotyphla | 5 | 14.610±0.435 | **0.383±0.125** | <0.0001 | 0.0548 | 0.758 |
| Marsupialia | 9 | 14.159±0.589 | **0.349±0.071** | <0.0001 | 0.0017 | 0.778 |
| Primata | 12 | 14.037±0.691 | **0.525±0.089** | <0.0001 | 0.0002 | 0.777 |
| Glires | 11 | 14.538±0.408 | **0.363±0.061** | <0.0001 | 0.0002 | 0.796 |
| Scandentia | 1 |  |  |  |  |  |

Most importantly, it is not the case that there was a simple, one-time grade shift in both mammals and birds with the advent of endothermy towards more brain neurons for a given body mass, as suggested recently (Kverkova et al., 2022). Instead, different clades of endothermic amniotes have different relationships between body mass and numbers of neurons in each brain structure (Figure 17, Table 10). The pallium of non-primate mammals and the early-derived bird clades does scale similarly in numbers of neurons together with increasing body mass (Figure 17A), but Accipitriformes and Falconiformes, then the late-derived Passeriformes, Psittaciformes and Strigiformes show a shift towards more pallial neurons in a similar body mass (Figures 17A, 18D), as do primates compared to other mammals (Herculano-Houzel et al., 2014a). At the same time, neuronal densities in the pallium/telencephalon remained similar across reptiles, early-derived birds and mammalian species of a similar body mass (Figure 17D). Likewise, the cerebellum of birds and mammals scales similarly in number of neurons with increasing body mass, but with a pronounced shift to more than 10 times as many cerebellar neurons for a given body mass (Figure 17B), with similarly increased neuronal densities in both groups (Figure 17E).

Importantly, the scaling of numbers of neurons in the “rest of brain” differs markedly between early-derived birds and mammals (Figure 11C). As a result, numbers of neurons in this collection of structures, which are directly involved in operating the body, vary by about 50-fold across reptiles, birds, and mammals of similar body mass. This discrepancy is consistent with recent reports that body mass and brain mass are quite free to vary independently of each other in mammalian and avian evolution (Ksepka et al., 2020; Bertrand et al., 2022) and that body mass is not a determinant of numbers of brain neurons (Herculano-Houzel et al., 2015b). Nevertheless, the striking finding remains that the parallel, independent shifts to endothermy in birds and mammals occurred with increased numbers of neurons in all brain structures compared to ancestral species of a similar body mass.

### Scaling relationships of mass ratios across structures distinguish reptiles from birds and mammals

The non-telencephalic, non-cerebellar rest of brain comprises the brain structures that operate the body. These “rest of brain” structures, which are readily comparable in organization and function across clades of amniotes, provide the anatomical and circuit-related scaffolding on top of which the pallium and the cerebellum function. It is also composed of structures derived from the near entirety of the cranial neural tube, minus the first two neuromeres (Puelles et al., 2013). Thus, I use the rest of brain as an anchor to which I compare the scaling of the other brain structures across clades of amniotes. This procedure allows for a measurement of relative allocation of neurons across brain structures that is completely irrespective of body mass or brain mass.

Despite the 100,000-fold range of variation in the mass of the rest of brain across the 242 species in the dataset, the pallium (or whole telencephalon, in reptiles) scales in mass faster than the rest of brain in a uniform, shared manner across all amniotes, with the notable exceptions of primates and owls (Strigiformes), with a shared exponent of 1.189±0.013, significantly above unity (Figure 18A; Table 11). As a consequence, the mass ratio between pallium (or telencephalon) and rest of brain increases together with the absolute mass of the rest of the brain in a continuous manner across reptiles, mammals, and birds (exponent, 0.189±0.013; Figure 18B; Table 12) minus primates and owls, which show a step increase away from the shared pattern, with a larger pallium than expected for an amniote of similar mass of the rest of brain (Figure 18B).

**Figure 18.**
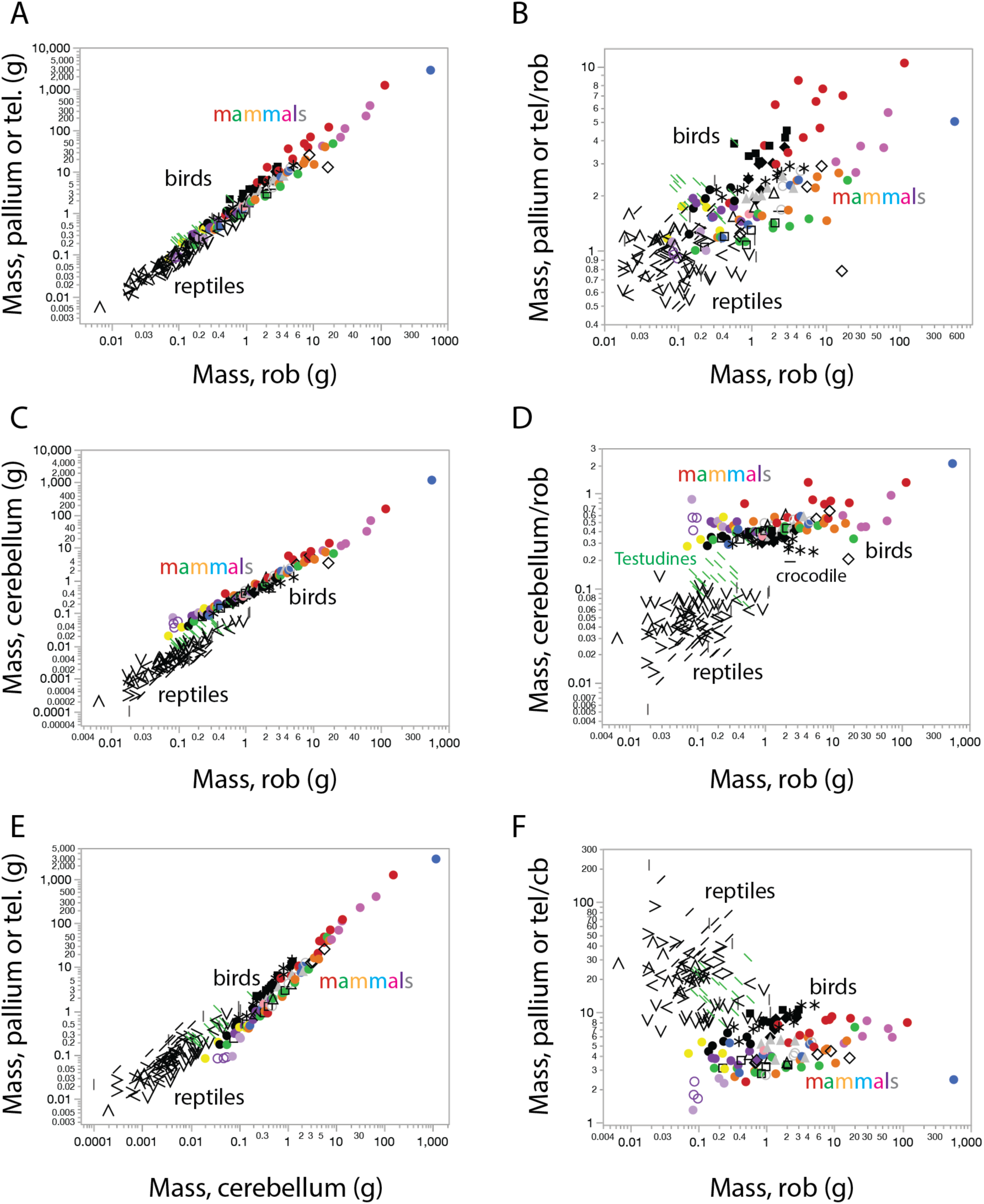
Mass scaling relationships across brain structures in mammals, reptiles and birds. **A, C, E,** scaling of structure mass across the structures indicated. **B, D, F,** scaling of the mass ratio across the brain structures indicated, as a function of the mass of the rest of brain. Each data point represents one species, using the same symbols as in all other figures. All applicable power functions, not plotted for clarity, are listed in Tables 11 and 12.

**Table 11.** Scaling of the mass of the telencephalon or pallium (Mtel; reptilian telencephalon or bird or mammalian pallium), mass of the cerebellum (Mcb), and mass of the rest of brain (Mrob; reptilian or bird brainstem and diencephalon, or mammalian brainstem, diencephalon plus striatum) as a function of one another in the different groups. Values correspond to the equation Mstructure1 = e^a±SE^ Mstructure2 ^b±SE^, the p-values for a and b, and the r^2^ for the power fit.

| $M_{tel} \times M_{rob}$ | n | a in $e^a$ | b in $N_{rob}^b$ | p, a | p, b | $r^2$ |
| --- | --- | --- | --- | --- | --- | --- |
| **Reptiles** |  |  |  |  |  |  |
| All | 108 | 0.341±0.092 | **1.112±0.038** | 0.0003 | <0.0001 | 0.891 |
| Testudines and crocodile | 20 | 0.614±0.115 | **0.952±0.067** | <0.0001 | <0.0001 | 0.918 |
| All minus Testudines and crocodile | 88 | 0.041±0.087 | **1.035±0.034** | 0.6410 | <0.0001 | 0.917 |
| **Mammals** |  |  |  |  |  |  |
| All | 69 | 0.640±0.051 | **1.231±0.024** | <0.0001 | <0.0001 | 0.975 |
| Primates | 12 | 1.137±0.144 | **1.294±0.069** | <0.0001 | <0.0001 | 0.972 |
| All minus primates | 57 | 0.533±0.037 | **1.193±0.017** | <0.0001 | <0.0001 | 0.988 |
| **All reptiles and mammals minus Testudines and crocodiles, and primates** | 145 | 0.457±0.032 | **1.189±0.013** | <0.0001 | <0.0001 | 0.983 |
| **Birds** |  |  |  |  |  |  |
| Anseriformes, Accipitriformes, Falconiformes | 14 | 0.571±0.049 | **1.227±0.065** | <0.0001 | <0.0001 | 0.968 |
| Palaeognathae, Galliformes, Columbiformes | 20 | 0.236±0.059 | **1.107±0.051** | 0.0009 | <0.0001 | 0.963 |
| Passeriformes, Strigiformes | 20 | 1.020±0.048 | **1.301±0.053** | <0.0001 | <0.0001 | 0.971 |
| Psittaciformes | 11 | 0.848±0.016 | **1.132**±0.018 | <0.0001 | <0.0001 | 0.998 |
| **M<sub>cb</sub> x M<sub>rob</sub>** | **n** | **a in e<sup>a</sup>** | **b in N<sub>rob</sub><sup>b</sup>** | **p, a** | **p, b** | **r<sup>2</sup>** |
| **Reptiles** |  |  |  |  |  |  |
| All | 108 | -2.225±0.139 | **1.318**±0.057 | <0.0001 | <0.0001 | 0.835 |
| Testudines and crocodile | 20 | -1.856±0.172 | **1.111**±0.100 | <0.0001 | <0.0001 | 0.872 |
| All minus Testudines and crocodile | 88 | -2.623±0.143 | **1.215**±0.056 | <0.0001 | <0.0001 | 0.848 |
| **Mammals** |  |  |  |  |  |  |
| All (minus elephant) | 67 | -0.712±0.038 | **1.080**±0.019 | <0.0001 | <0.0001 | 0.981 |
| Primates | 11 | -0.481±0.145 | **1.123**±0.067 | 0.0091 | <0.0001 | 0.969 |
| All minus primates, elephant | 56 | -0.762±0.035 | **1.055**±0.018 | <0.0001 | <0.0001 | 0.985 |
| **Birds** |  |  |  |  |  |  |
| All | 65 | -1.004±0.027 | **1.054**±0.028 | <0.0001 | <0.0001 | 0.958 |
| All minus Psittaciformes | 54 | -0.961±0.026 | **1.084**±0.027 | <0.0001 | <0.0001 | 0.970 |
| Psittaciformes | 11 | -1.198±0.032 | **0.913**±0.036 | <0.0001 | <0.0001 | 0.986 |
| **M<sub>tel</sub> x M<sub>cb</sub>** | **n** | **a in e<sup>a</sup>** | **b in N<sub>rob</sub><sup>b</sup>** | **p, a** | **p, b** | **r<sup>2</sup>** |
| **Reptiles** |  |  |  |  |  |  |
| All | 108 | 1.634±0.190 | **0.730**±0.036 | <0.0001 | <0.0001 | 0.799 |
| Testudines and crocodile | 20 | 1.905±0.275 | **0.772**±0.075 | <0.0001 | <0.0001 | 0.855 |
| All minus Testudines and crocodile | 88 | 1.407±0.265 | **0.694**±0.047 | <0.0001 | <0.0001 | 0.719 |
| **Mammals** |  |  |  |  |  |  |
| All (minus elephant) | 67 | 1.444±0.039 | **1.143**±0.018 | <0.0001 | <0.0001 | 0.983 |
| Primates | 11 | 1.651±0.139 | **1.149**±0.066 | <0.0001 | <0.0001 | 0.971 |
| All minus primates, elephant | 56 | 1.395±0.040 | **1.124**±0.019 | <0.0001 | <0.0001 | 0.984 |
| **Birds** |  |  |  |  |  |  |
| Anseriformes,<br>Accipitriformes,<br>Falconiformes | 14 | 1.460±0.066 | **1.007**±0.087 | <0.0001 | <0.0001 | 0.917 |
| Palaeognathae,<br>Galliformes,<br>Columbiformes | 20 | 1.304±0.023 | **1.060**±0.016 | <0.0001 | <0.0001 | 0.996 |
| Passeriformes,<br>Psittaciformes,<br>Strigiformes | 31 | 2.280±0.041 | **1.226**±0.029 | <0.0001 | <0.0001 | 0.984 |

**Table 12.**
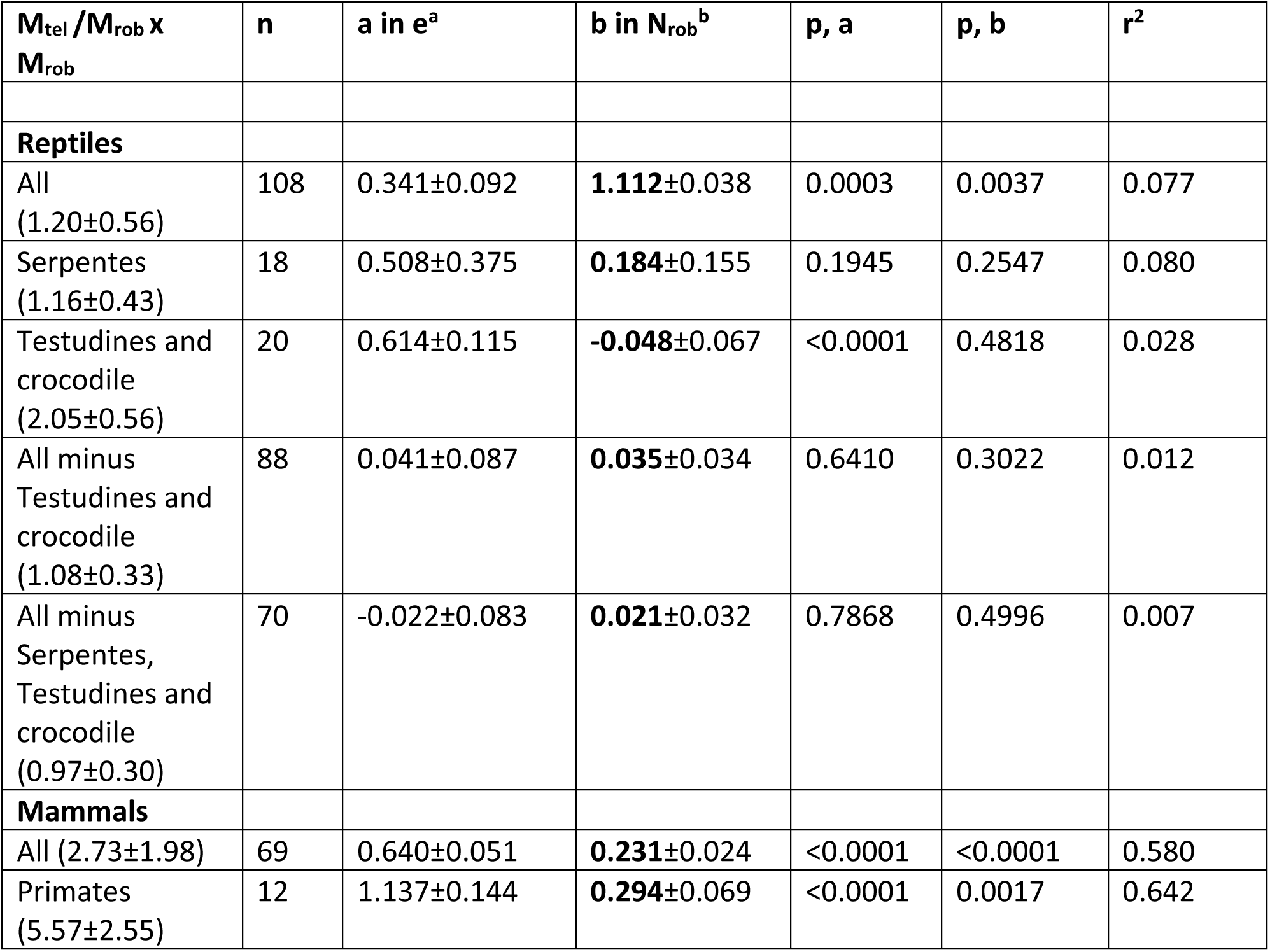
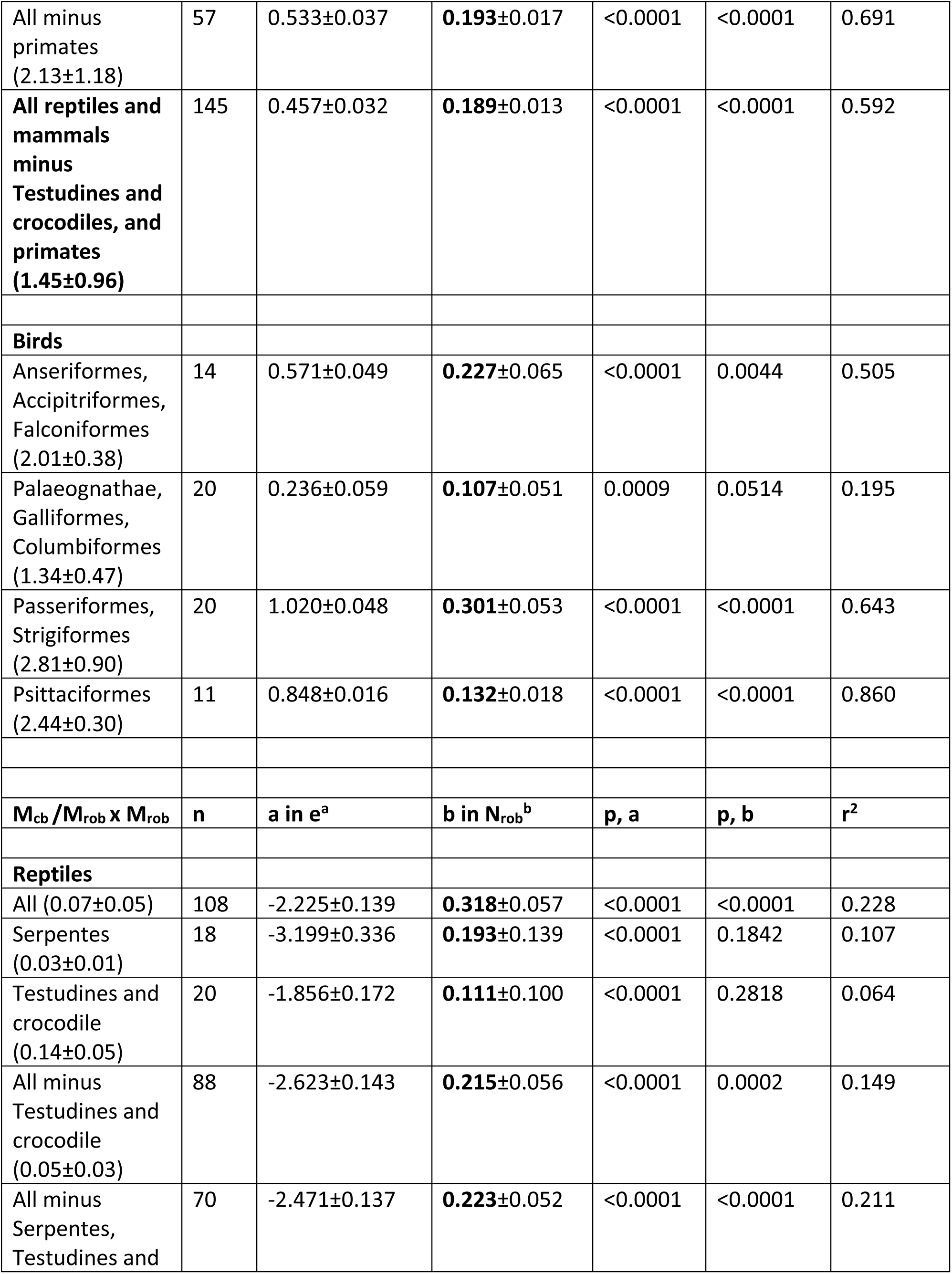
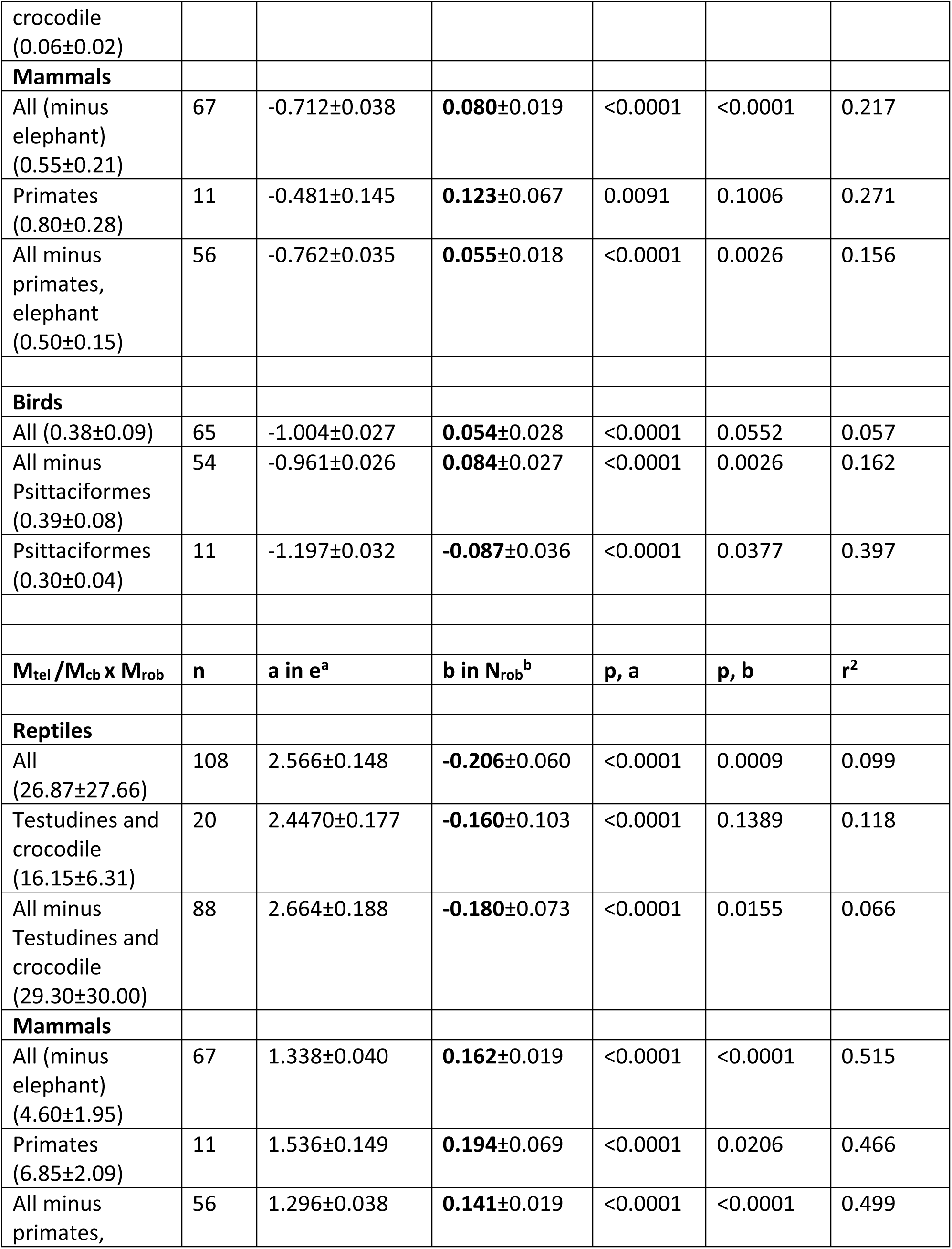
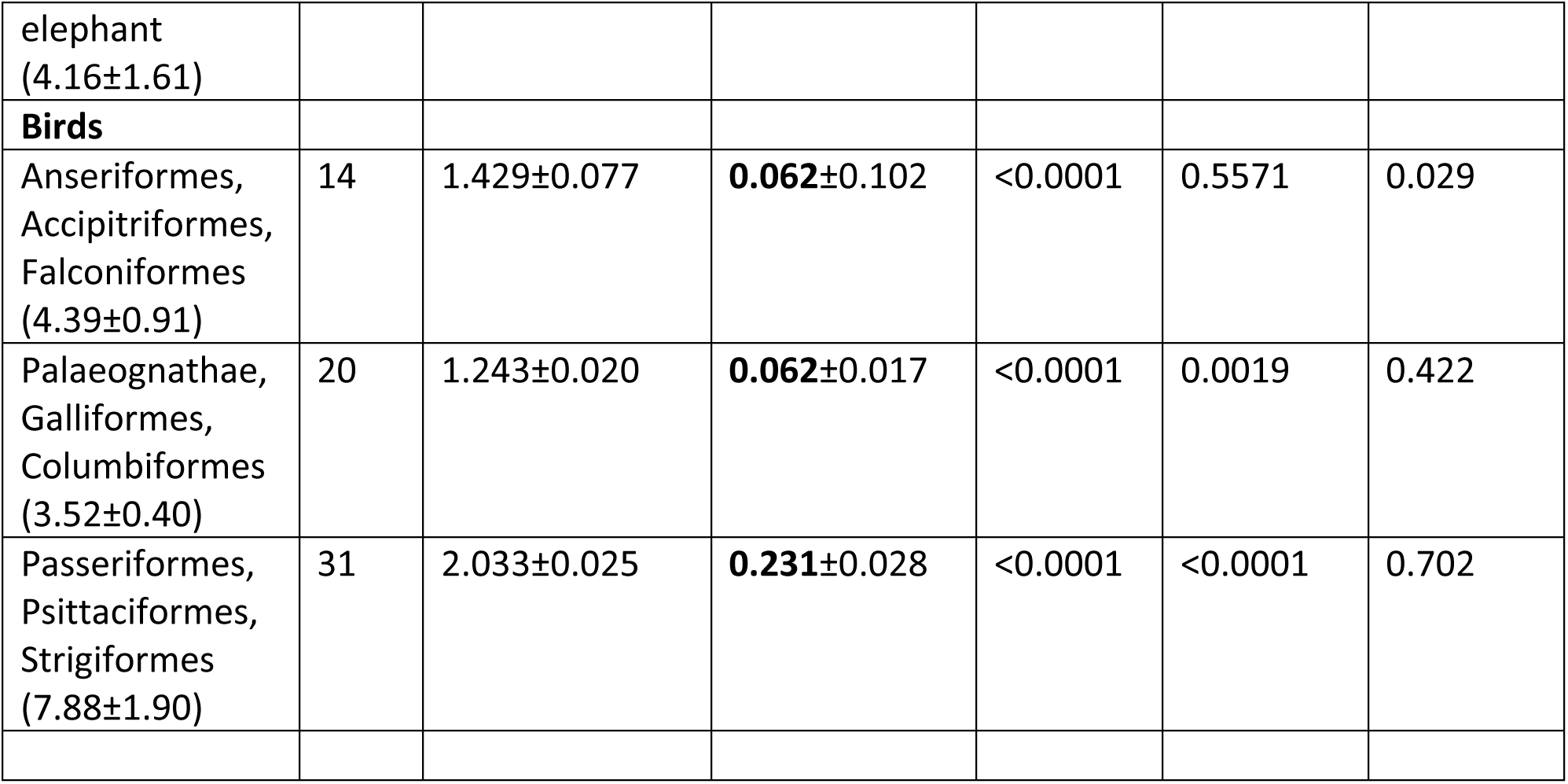
Scaling of the mass ratios between structures as a function of the mass of the rest of the brain in the different groups. Average ratios across species within each group ± SD are provided in parentheses. Mtel, mass of the reptilian telencephalon or bird or mammalian pallium; Mcb, mass of the cerebellum; Mrob, mass of the rest of brain, which is the reptilian or bird brainstem and diencephalon, or the mammalian brainstem, diencephalon plus striatum. Values correspond to the equation Mstructure1 = e^a±SE^ Mstructure2 ^b±SE^, the p-values for a and b, and the r^2^ for the power fit.

In contrast, the cerebellum of mammals and birds, the endothermic amniotes, shows a marked difference from the scaling relationship in mass with the rest of brain that applies to the non-avian reptiles in the dataset, with ca. 10-fold larger cerebellar mass in the warm-blooded amniotes compared to reptiles of similar rest of brain mass (Figure 18C). As a result, while the cerebellum is on average only about 1/20^th^ the mass of the rest of brain in reptiles, it jumps to a mass equivalent to ca. 1/3 to ½ the mass of the rest of brain in birds and mammals (Figure 18D, Table 12). Strikingly, the mass scaling relationship between cerebellum and rest of brain is shared across most avian and mammalian species (Figure 18C-D), although the cerebellum appears relatively smaller in parrots than in mammals of similar rest of brain mass (30% of the mass of the rest of brain or less in parrots vs. 50% in non-primate mammals; Table 12). As a consequence, the ratio between the mass of the pallium (or telencephalon) and of the cerebellum is very large in reptiles, whose telencephalons may be from 6 to over 100 times larger than the cerebellum, but this ratio is 4-10 in birds, and only 3-5 in most mammals (Figure 18E-F, Table 12). Thus, examination of the scaling relationships that apply to structure mass across the three main divisions of amniote brains suggests that a distinctive signature of endothermic amniotes is increased relative mass of the cerebellum, in a mass proportionality to the rest of brain that is shared by the vast majority of the evolutionarily unrelated mammals and birds. Remarkably, the shift to endothermy per se does not appear to have changed the proportionality with which telencephalic mass (whether cortical, pallial, or telencephalic as a whole) is added on top of the rest of brain, which remains shared across most amniotes. In terms of scaling of brain structure mass, the main difference in how bird and mammalian brains are built is that there is slightly more pallial mass relative to the cerebellum in songbirds, parrots and owls than in other birds and in non-primate mammals (Figure 18F) – but otherwise, bird and non-primate mammalian brains are built to fairly similar proportions across their main structures. These proportions are illustrated in Figure 19.

**Figure 19.**
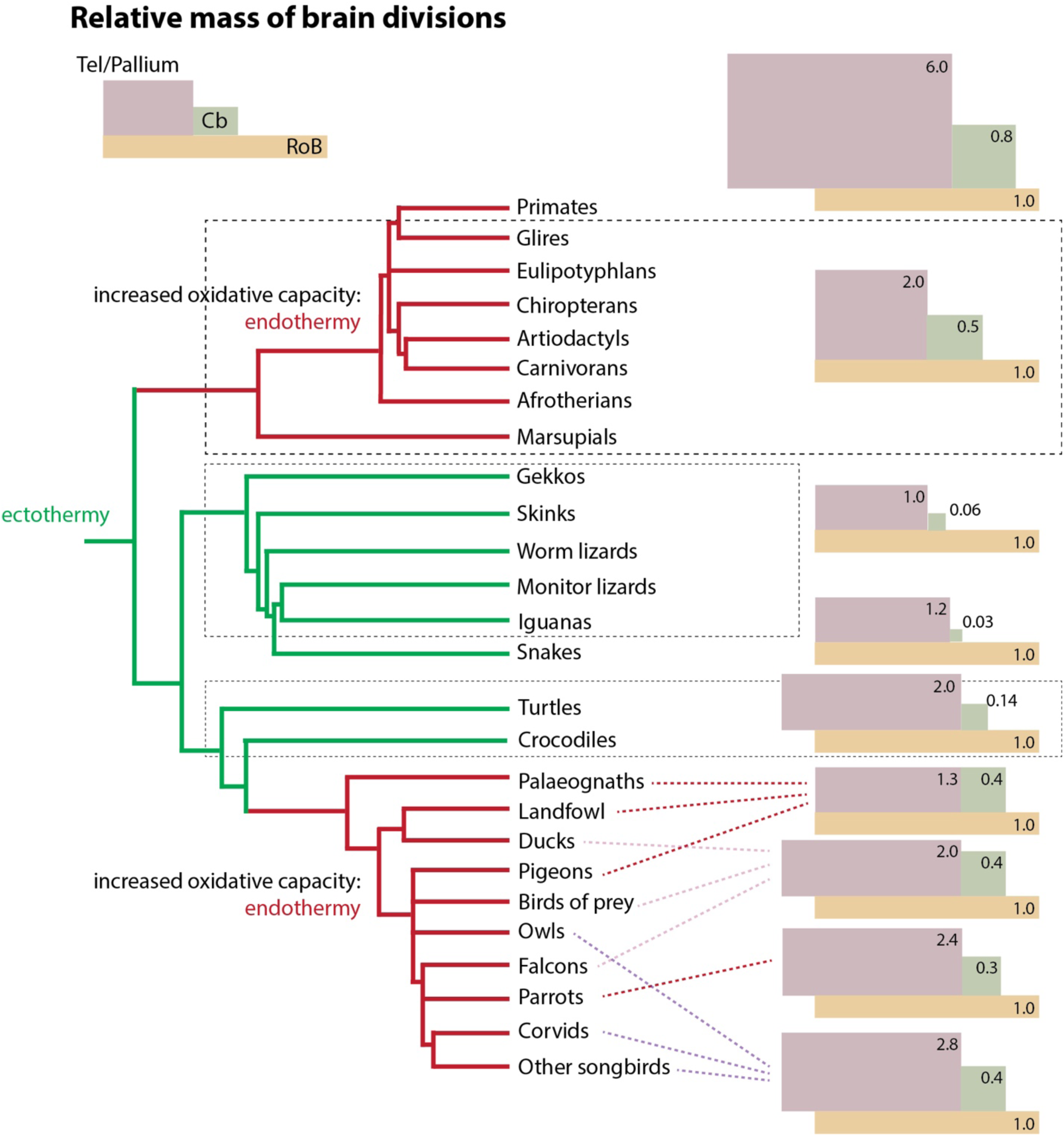
A relatively enlarged cerebellum is a signature of the brain of endothermic amniotes. The non-overlapping rectangles of varying sizes represent the relative mass of the telencephalon or pallium (red) and cerebellum (green) in proportion to the rest of brain (tan rectangle). The surface area of each rectangle is proportional to the mass of that brain structure relative to the rest of brain. The schematics illustrates the finding that while relative enlargement of the telencephalon over the rest of brain does not distinguish endothermic from ectothermic clades, the two independently evolved endothermic clades (mammals and avians) are characterized by a marked increase in mass of the cerebellum over the rest of brain compared to reptiles. Note that the distribution of relative brain structure mass can be shared across unrelated groups among birds.

### Mass relationships across structures differ from scaling of numbers of neurons

Despite the enormous diversity in absolute numbers of neurons composing the brains of the different amniotes, striking patterns emerge in how they vary across structures within each group of amniotes. While the scaling of structure mass does not distinguish between bird and mammalian brains, the scaling of numbers of neurons across structures does, and indeed characterizes each of the three groups of amniotes, as I show next.

The mass relationships across brain structures do not inform about the scaling of numbers of neurons across brain structures in amniotes. The shared scaling of pallial (or telencephalic) mass as a function of the mass of the rest of brain across most amniotes in the dataset (Figure 18A,B), with the exception of owls and primates, conceals the fact that artiodactyls, carnivorans and large marsupials, besides owls and primates, have increased numbers of pallial neurons compared to the prediction for a generic amniote of their numbers of neurons in the rest of brain (Figure 20A; Table 13). As a result, these clades have markedly larger ratios of numbers of neurons in the pallium over the rest of brain that set them apart from all other amniotes (Figure 20B).

**Figure 20.**
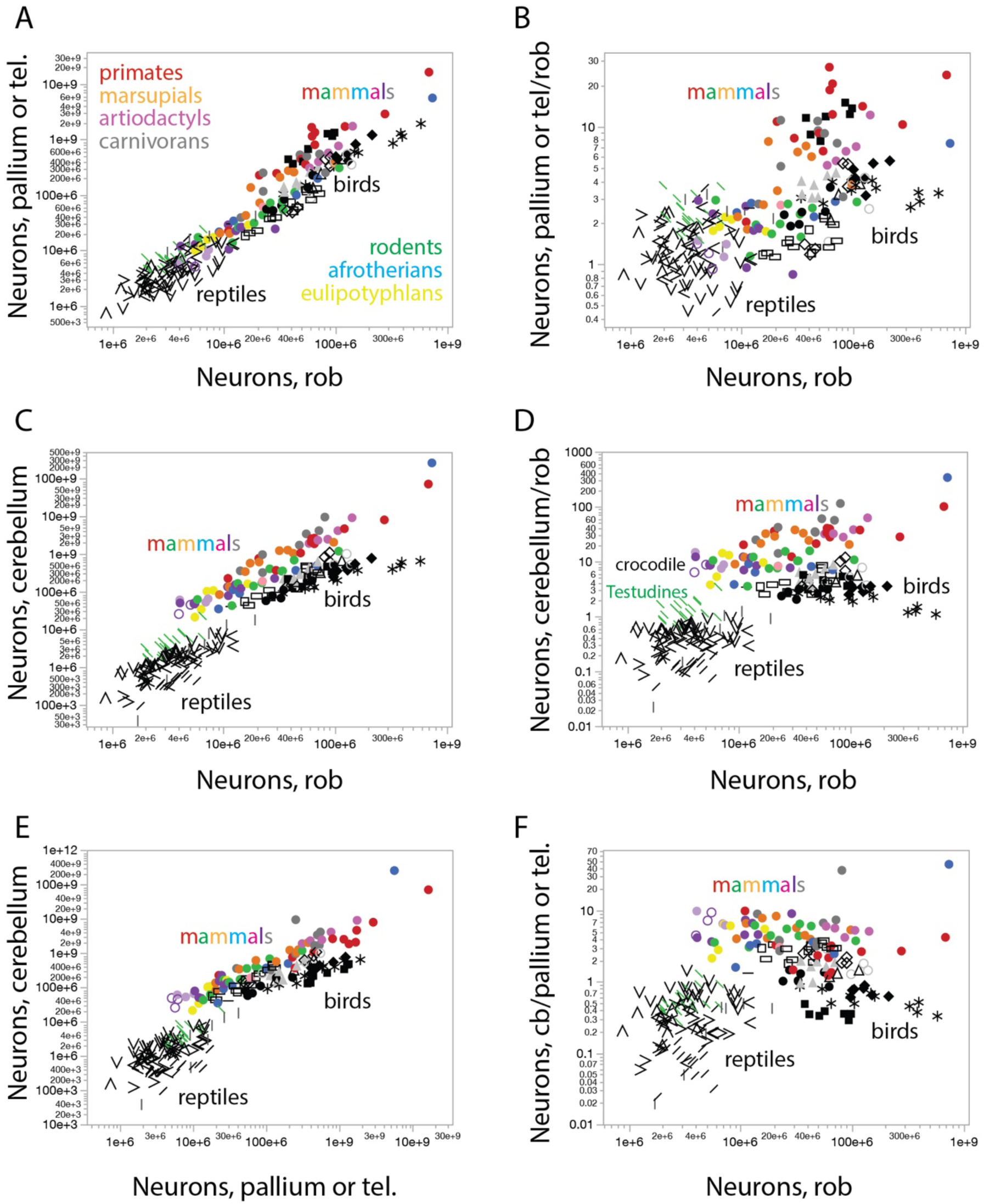
Scaling of numbers of neurons across structures defines clades of amniotes. **A, B,** the pallium or telencephalon gains neurons coordinately with the rest of brain across reptiles, birds, and all small mammals, while some large mammals, with larger numbers of neurons in the rest of brain, have significantly increased numbers of pallial neurons over the rest of brain (primates, red; artiodactyls, lavender; carnivorans, gray; marsupials, orange). Power functions and clade averages are listed in Tables 13 and 16. **C**, **D,** the cerebellum gains neurons in clade-specific relationships with the rest of brain across reptiles, birds, and mammals, while some large mammals, with larger numbers of neurons in the rest of brain, have significantly increased numbers of cerebellar neurons. Power functions and clade averages are listed in Tables 14 and 17. **E**, **F:** as a result of the parallels between the distributions shown in (**A, B**) and (**C, D**), the ratio between cerebellar and pallial (or telencephalic) neurons does not scale significantly with numbers of neurons in the rest of brain across reptiles and birds, and is fairly constant at a higher value across mammalian species. Interestingly, this ratio is higher in early-derived birds than in later-derived birds. Power functions and clade averages are listed in Tables 15 and 18.

**Table 13.** Scaling of neurons in the reptilian telencephalon or bird/mammalian pallium (Ntel) as a function of numbers of neurons in the rest of brain (Nrob; reptilian or bird brainstem and diencephalon, or mammalian brainstem, diencephalon plus striatum). Values correspond to the equation NTel = e^a±SE^ Nrob^b±SE^, the p-values for a and b, and the r^2^ for the power fit.

| $N_{\text{Tel}} \times N_{\text{rob}}$ | n | a in $e^a$ | b in $N_{\text{rob}}^b$ | p, a | p, b | $r^2$ |
| --- | --- | --- | --- | --- | --- | --- |
| **Reptiles** |  |  |  |  |  |  |
| All | 108 | -0.168±1.252 | **1.035**±0.083 | 0.8935 | <0.0001 | 0.594 |
| Anguimorpha | 7 | -1.508±3.300 | **1.145**±0.211 | 0.6668 | 0.0029 | 0.854 |
| Crocodylia | 1 |  |  |  |  |  |
| Gekkota | 14 | -3.634±3.081 | **1.271**±0.211 | 0.2612 | <0.0001 | 0.752 |
| Iguania | 20 | -1.524±1.977 | **1.087**±0.129 | 0.4508 | <0.0001 | 0.797 |
| Lacertoidea | 15 | -2.045±2.602 | **1.149**±0.172 | 0.4461 | <0.0001 | 0.775 |
| Scincoidea | 14 | 0.078±2.390 | **1.023**±0.160 | 0.9744 | <0.0001 | 0.772 |
| Serpentes | 18 | 0.846±5.039 | **0.970**±0.335 | 0.8688 | 0.0106 | 0.344 |
| Testudines and Crocodylia | 20 | 0.473±1.937 | **1.025**±0.129 | 0.8099 | <0.0001 | 0.779 |
| All minus Serpentes, Testudines, Crocodylia | 70 | -0.542±1.275 | **1.050**±0.084 | 0.6722 | <0.0001 | 0.694 |
| **Birds** |  |  |  |  |  |  |
| All | 65 | -5.027±1.826 | **1.341**±0.101 | 0.0077 | <0.0001 | 0.736 |
| Accipitriformes | 4 | -15.931±12.473 | **1.921**±0.677 | 0.3298 | 0.1049 | 0.801 |
| Anseriformes | 7 | -5.344±4.309 | **1.376**±0.244 | 0.2699 | 0.0024 | 0.864 |
| Columbiformes | 5 | -5.907±3.841 | **1.365**±0.222 | 0.2216 | 0.0086 | 0.927 |
| Falconiformes | 3 | -14.767±8.525 | **1.870**±0.468 | 0.3333 | 0.1561 | 0.941 |
| Galliformes | 9 | -4.157±2.056 | **1.259**±0.118 | 0.0830 | <0.0001 | 0.942 |
| Palaeognathae | 6 | -28.708±5.017 | **2.654**±0.280 | 0.0046 | 0.0007 | 0.957 |
| Passeriformes | 13 | -7.678±1.235 | **1.491**±0.068 | <0.0001 | <0.0001 | 0.977 |
| Psittaciformes | 11 | 1.707±0.991 | **0.973**±0.053 | 0.1192 | <0.0001 | 0.974 |
| Strigiformes | 7 | -4.614±3.495 | **1.392**±0.195 | 0.2440 | 0.0008 | 0.911 |
| All minus Strigiformes | 58 | -5.559±1.220 | **1.362**±0.068 | <0.0001 | <0.0001 | 0.879 |
| **Mammals** |  |  |  |  |  |  |
| All | 70 | 11.563±0.103 | **-0.304**±0.014 | <0.0001 | <0.0001 | 0.881 |
| All non-primates | 58 | 11.592±0.099 | **-0.315**±0.013 | <0.0001 | <0.0001 | 0.913 |
| Afrotheria | 6 | 11.372±0.306 | **-0.265**±0.306 | <0.0001 | 0.0020 | 0.927 |
| Artiodactyla | 5 | 9.726±1.751 | **-0.161±0.150** | 0.0115 | 0.3630 | 0.276 |
| Carnivora | 8 | 10.916±0.385 | **-0.271±0.039** | <0.0001 | 0.0005 | 0.888 |
| Chiroptera | 13 | 11.256±0.228 | **-0.235±0.062** | <0.0001 | 0.0029 | 0.570 |
| Eulipotyphla | 5 | 11.689±0.502 | **-0.285±0.144** | 0.0002 | 0.1425 | 0.566 |
| Marsupialia | 9 | 11.628±0.540 | **-0.330±0.065** | <0.0001 | 0.0014 | 0.788 |
| Primata | 12 | 10.780±0.407 | **-0.171±0.052** | <0.0001 | 0.0085 | 0.516 |
| Glires | 10 | 11.735±0.410 | **-0.314±0.059** | <0.0001 | 0.0007 | 0.778 |
| Scandentia | 1 |  |  |  |  |  |
| All mammals, all reptiles | 177 | 11.725±0.060 | **-0.301±0.010** | <0.0001 | <0.0001 | 0.850 |

**Table 14.** Scaling of neurons in the cerebellum (Ncb) as a function of numbers of neurons in the rest of brain (Nrob; reptilian or bird brainstem and diencephalon, or mammalian brainstem, diencephalon plus striatum). Values correspond to the equation Ncb = e^a±SE^ Nrob^b±SE^, the p-values for a and b, and the r^2^ for the power fit.

| $N_{Cb} \times N_{rob}$ | n | a in $e^a$ | b in $N_{rob}^b$ | p, a | p, b | $r^2$ |
| --- | --- | --- | --- | --- | --- | --- |
| **Reptiles** |  |  |  |  |  |  |
| All | 108 | -7.329±2.201 | **1.429**±0.146 | 0.0012 | <0.0001 | 0.475 |
| Anguimorpha | 7 | -25.867±6.062 | **2.597**±0.388 | 0.0080 | 0.0011 | 0.900 |
| Crocodylia | 1 |  |  |  |  |  |
| Gekkota | 14 | -9.390±4.080 | **1.587**±0.279 | 0.0401 | 0.0001 | 0.729 |
| Iguania | 20 | -0.185±1.526 | **0.977**±0.100 | 0.9049 | <0.0001 | 0.842 |
| Lacertoidea | 15 | -3.750±3.205 | **1.172**±0.211 | 0.2631 | <0.0001 | 0.703 |
| Scincoidea | 14 | -8.557±3.160 | **1.492**±0.212 | 0.0190 | <0.0001 | 0.805 |
| Serpentes | 18 | -5.528±4.058 | **1.233**±0.270 | 0.1920 | 0.0003 | 0.566 |
| Testudines and Crocodylia | 20 | -8.934±2.882 | **1.612**±0.191 | 0.0062 | <0.0001 | 0.798 |
| All minus Serpentes, Testudines, Crocodylia | 70 | -7.008±1.653 | **1.405**±0.110 | <0.0001 | <0.0001 | 0.708 |
| **Birds** |  |  |  |  |  |  |
| All | 65 | 4.487±1.462 | **0.822**±0.081 | 0.0032 | <0.0001 | 0.620 |
| Accipitriformes | 4 | -2.245±10.904 | **1.209**±0.591 | 0.8559 | 0.1776 | 0.676 |
| Anseriformes | 7 | -13.063±6.875 | **1.837**±0.389 | 0.1159 | 0.0053 | 0.817 |
| Columbiformes | 5 | -8.072±5.515 | **1.549**±0.318 | 0.2395 | 0.0166 | 0.888 |
| Falconiformes | 3 | -20.443±10.178 | **2.196**±0.559 | 0.2941 | 0.1586 | 0.939 |
| Galliformes | 9 | -5.606±4.027 | **1.399**±0.230 | 0.2065 | 0.0005 | 0.841 |
| Palaeognathae | 6 | -21.326±5.267 | **2.287**±0.294 | 0.0155 | 0.0015 | 0.938 |
| Passeriformes | 13 | -1.796±1.077 | **1.160**±0.060 | 0.1237 | <0.0001 | 0.972 |
| Psittaciformes | 11 | 6.063±1.284 | **0.709**±0.068 | 0.0011 | <0.0001 | 0.923 |
| Strigiformes | 7 | -1.128±5.866 | **1.141±0.327** | 0.8551 | 0.0174 | 0.709 |
| All minus Psittaciformes | 54 | -2.085±1.627 | **1.196±0.091** | 0.2056 | <0.0001 | 0.768 |
| **Mammals** |  |  |  |  |  |  |
| All | 70 | 11.563±0.103 | **-0.304±0.014** | <0.0001 | <0.0001 | 0.881 |
| All non-primates | 58 | 11.592±0.099 | **-0.315±0.013** | <0.0001 | <0.0001 | 0.913 |
| Afrotheria | 6 | 11.372±0.306 | **-0.265±0.306** | <0.0001 | 0.0020 | 0.927 |
| Artiodactyla | 5 | 9.726±1.751 | **-0.161±0.150** | 0.0115 | 0.3630 | 0.276 |
| Carnivora | 8 | 10.916±0.385 | **-0.271±0.039** | <0.0001 | 0.0005 | 0.888 |
| Chiroptera | 13 | 11.256±0.228 | **-0.235±0.062** | <0.0001 | 0.0029 | 0.570 |
| Eulipotyphla | 5 | 11.689±0.502 | **-0.285±0.144** | 0.0002 | 0.1425 | 0.566 |
| Marsupialia | 9 | 11.628±0.540 | **-0.330±0.065** | <0.0001 | 0.0014 | 0.788 |
| Primata | 12 | 10.780±0.407 | **-0.171±0.052** | <0.0001 | 0.0085 | 0.516 |
| Glires | 10 | 11.735±0.410 | **-0.314±0.059** | <0.0001 | 0.0007 | 0.778 |
| Scandentia | 1 |  |  |  |  |  |
| All mammals, all reptiles | 177 | 11.725±0.060 | **-0.301±0.010** | <0.0001 | <0.0001 | 0.850 |

**Table 15.** Scaling of neurons in the cerebellum (N_cb_) as a function of numbers of neurons in the telencephalon or pallium (N_tel_; reptilian telencephalon or bird or mammalian pallium). Values correspond to the equation N_cb_ = e^a±SE^ N_tel_^b±SE^, the p-values for a and b, and the r^2^ for the power fit.

| $N_{cb} \times N_{tel}$ | n | a in $e^a$ | b in $N_{rob}^b$ | p, a | p, b | $r^2$ |
| --- | --- | --- | --- | --- | --- | --- |
| **Reptiles** |  |  |  |  |  |  |
| All | 108 | -1.222±1.767 | **1.000±0.114** | 0.4904 | <0.0001 | 0.419 |
| Anguimorpha | 7 | -19.526±5.283 | **2.091±0.323** | 0.0141 | 0.0013 | 0.894 |
| Crocodylia | 1 |  |  |  |  |  |
| Gekkota | 14 | -1.849±3.082 | **1.047±0.206** | 0.5597 | 0.0003 | 0.683 |
| Iguania | 20 | 2.548±1.187 | **0.809±0.079** | 0.0457 | <0.0001 | 0.855 |
| Lacertoidea | 15 | 0.378±2.560 | **0.888±0.167** | 0.8848 | 0.0001 | 0.686 |
| Scincoidea | 14 | -3.054±4.078 | **1.092±0.266** | 0.4684 | 0.0015 | 0.584 |
| Serpentes | 18 | 7.087±3.523 | **0.384±0.228** | 0.0614 | 0.1120 | 0.150 |
| Testudines and Crocodilia | 20 | -6.533±2.718 | **1.375±0.171** | 0.0272 | <0.0001 | 0.782 |
| All minus Serpentes, Testudines, Crocodilia | 70 | -0.005±1.759 | **0.928±0.115** | 0.9977 | <0.0001 | 0.490 |
| **Birds** |  |  |  |  |  |  |
| All | 65 | 9.310±1.006 | **0.522±0.052** | <0.0001 | <0.0001 | 0.611 |
| Accipitriformes | 4 | 7.638±3.477 | **0.637±0.178** | 0.1592 | 0.0703 | 0.864 |
| Anseriformes | 7 | -1.860±6.735 | **1.120±0.355** | 0.7934 | 0.0253 | 0.665 |
| Columbiformes | 5 | -0.881±3.526 | **1.107±0.199** | 0.8189 | 0.0114 | 0.912 |
| Falconiformes | 3 | -3.120±0.088 | **1.176±0.005** | 0.0179 | 0.0025 | 0.999 |
| Galliformes | 9 | -1.401±2.108 | **1.134±0.118** | 0.5277 | <0.0001 | 0.930 |
| Palaeognathae | 6 | 3.352±0.923 | **0.865±0.049** | 0.0221 | <0.0001 | 0.987 |
| Passeriformes | 13 | 4.395±0.840 | **0.767±0.043** | 0.0003 | <0.0001 | 0.966 |
| Psittaciformes | 11 | 4.876±1.224 | **0.727±0.061** | 0.0032 | <0.0001 | 0.940 |
| Strigiformes | 7 | 1.172±2.350 | **0.893±0.115** | 0.6390 | 0.0006 | 0.923 |
| Anseriformes, Accipitriformes, Galliformes, Columbiformes, Palaeognathae | 31 | 3.991±1.100 | **0.826±0.059** | 0.0011 | <0.0001 | 0.869 |
| Passeriformes, Psittaciformes | 24 | 5.600±0.874 | **0.698±0.044** | <0.0001 | <0.0001 | 0.918 |
| **Mammals** |  |  |  |  |  |  |
| All | 70 | 11.563±0.103 | **-0.304±0.014** | <0.0001 | <0.0001 | 0.881 |
| All non-primates | 58 | 11.592±0.099 | **-0.315±0.013** | <0.0001 | <0.0001 | 0.913 |
| Afrotheria | 6 | 11.372±0.306 | **-0.265±0.306** | <0.0001 | 0.0020 | 0.927 |
| Artiodactyla | 5 | 9.726±1.751 | **-0.161±0.150** | 0.0115 | 0.3630 | 0.276 |
| Carnivora | 8 | 10.916±0.385 | **-0.271±0.039** | <0.0001 | 0.0005 | 0.888 |
| Chiroptera | 13 | 11.256±0.228 | **-0.235±0.062** | <0.0001 | 0.0029 | 0.570 |
| Eulipotyphla | 5 | 11.689±0.502 | **-0.285±0.144** | 0.0002 | 0.1425 | 0.566 |
| Marsupialia | 9 | 11.628±0.540 | **-0.330±0.065** | <0.0001 | 0.0014 | 0.788 |
| Primata | 12 | 10.780±0.407 | **-0.171±0.052** | <0.0001 | 0.0085 | 0.516 |
| Glires | 10 | 11.735±0.410 | **-0.314±0.059** | <0.0001 | 0.0007 | 0.778 |
| Scandentia | 1 |  |  |  |  |  |
| All mammals, all reptiles | 177 | 11.725±0.060 | **-0.301±0.010** | <0.0001 | <0.0001 | 0.850 |

**Table 16.**
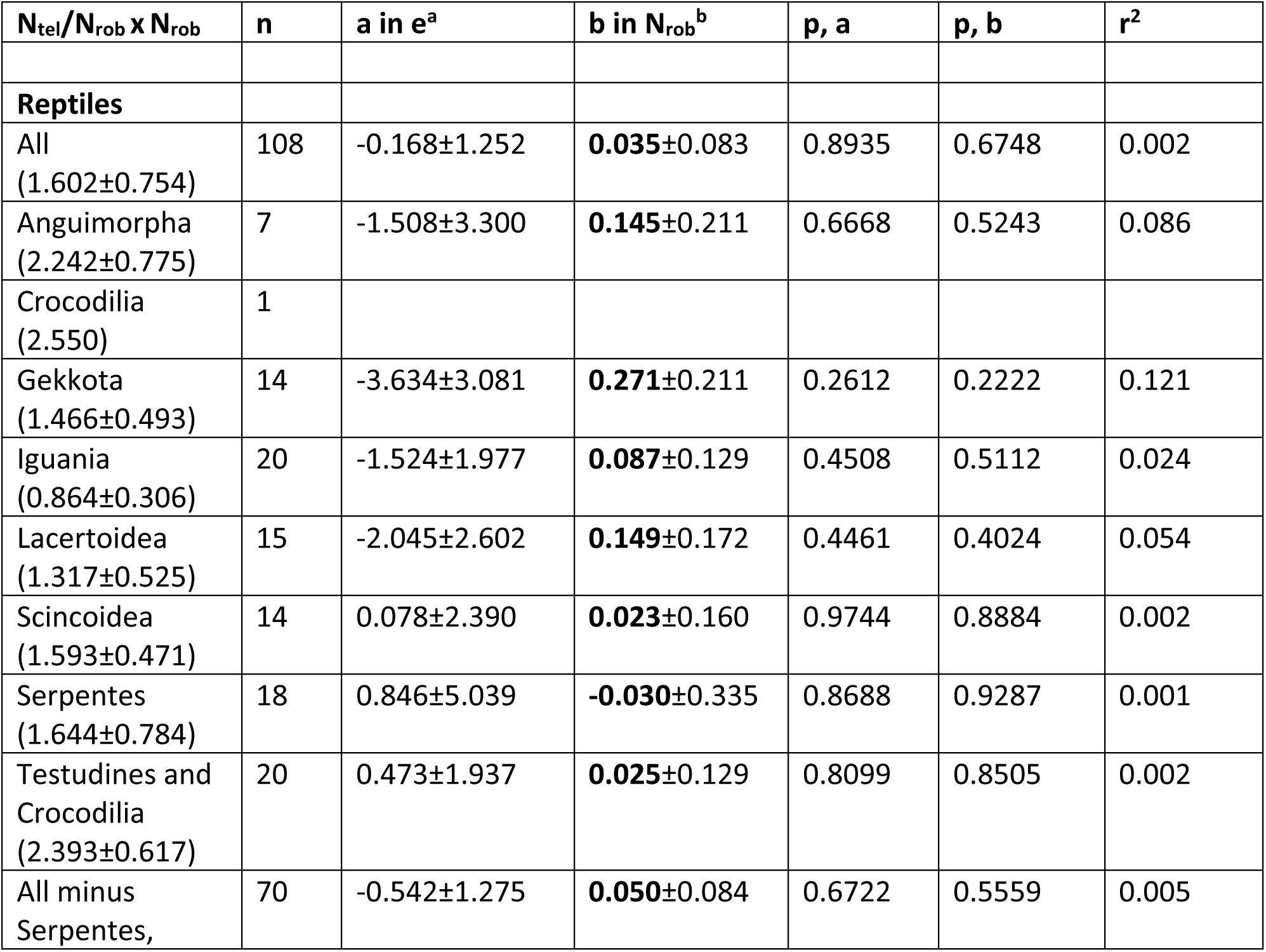
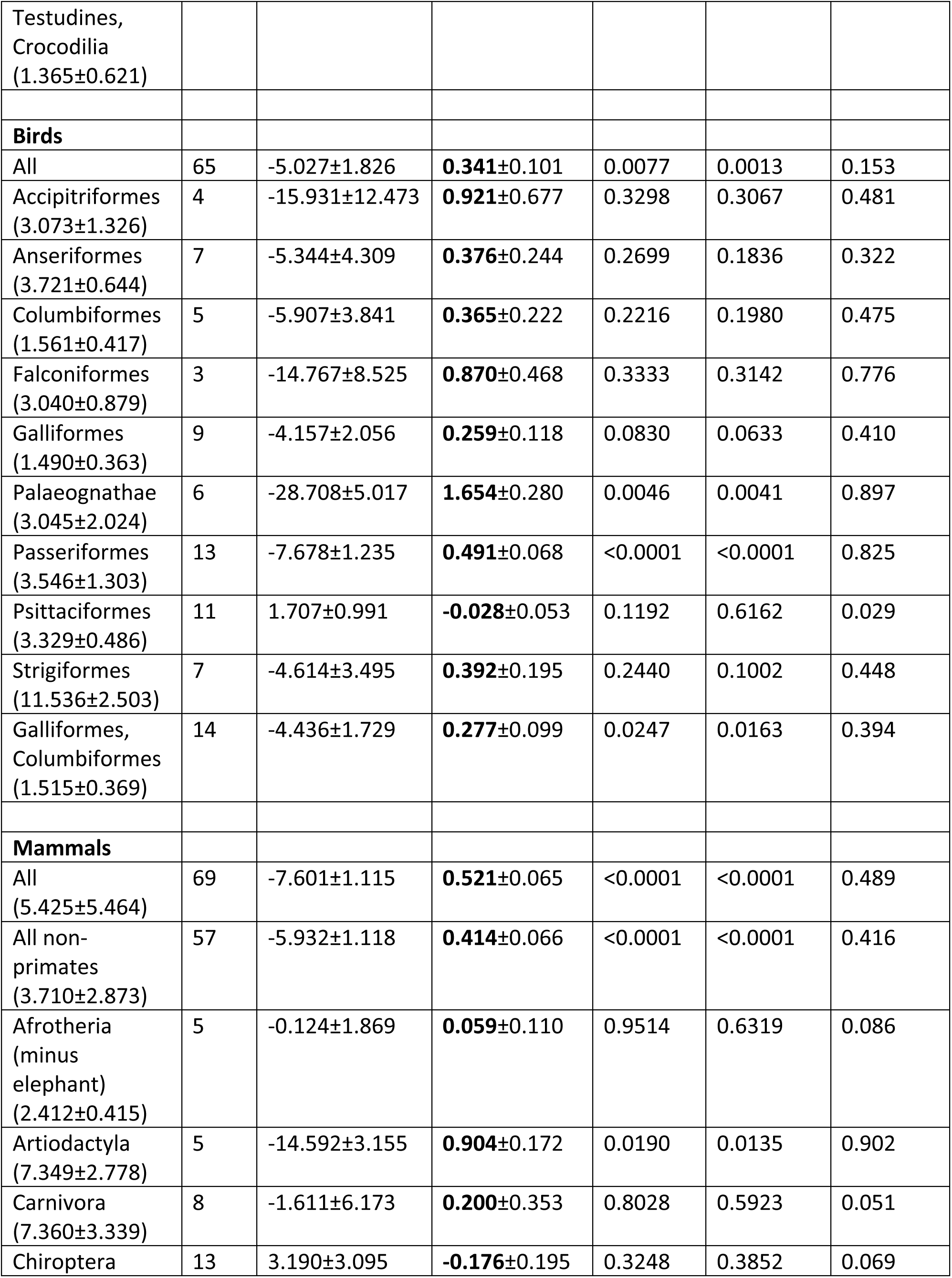
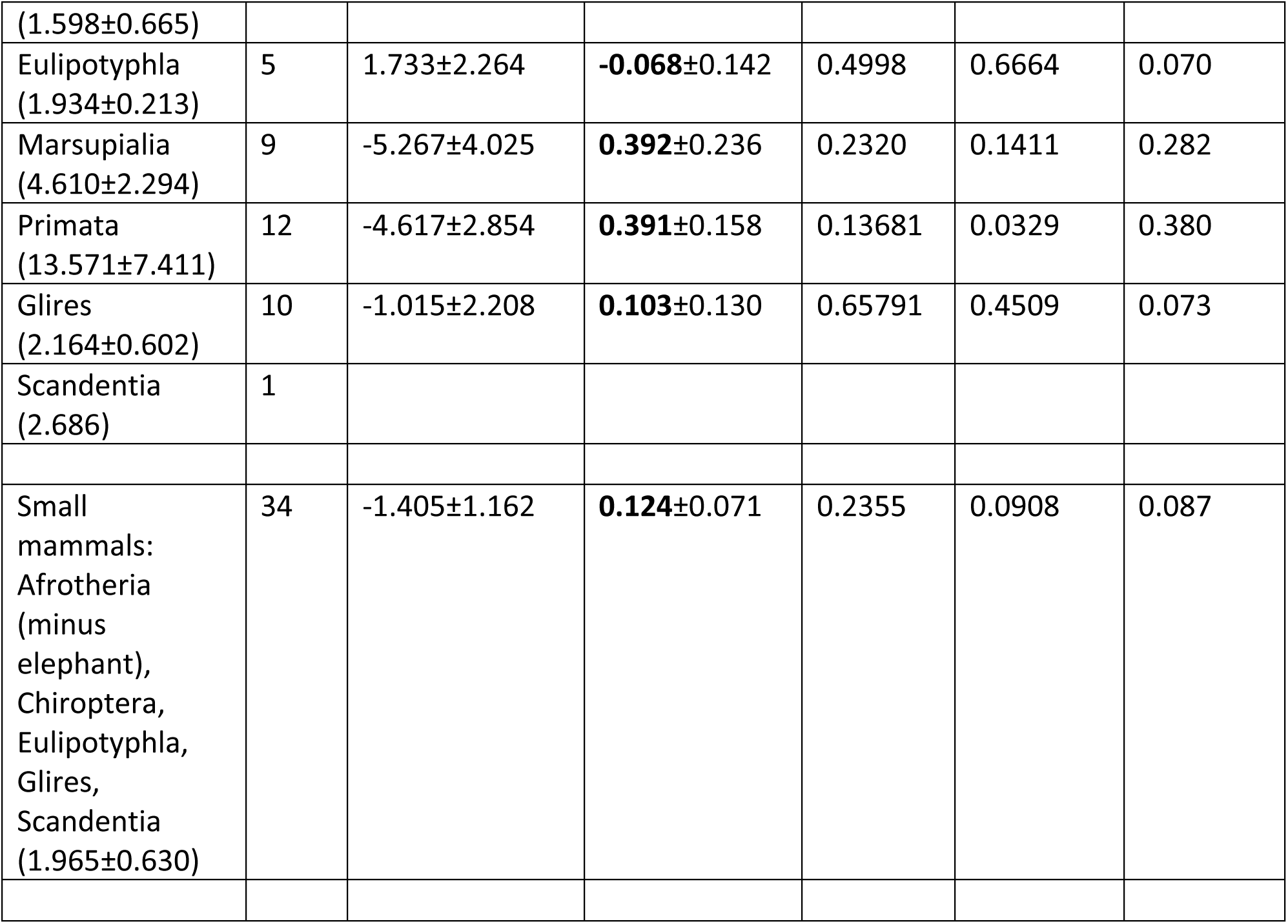
Scaling of the ratio Ntel/Nrob between neurons in the telencephalon or pallium (Ntel; reptilian telencephalon or bird or mammalian pallium) and in the rest of brain (Nrob; reptilian or bird brainstem and diencephalon, or mammalian brainstem, diencephalon plus striatum) as a function of numbers of neurons in the rest of brain (Nrob). Values correspond to the equation Ntel/Nrob = e^a±SE^ Nrob^b±SE^, the p-values for a and b, and the r^2^ for the power fit. Values in parentheses are the average ratio Ntel/Nrob for the clade ± standard deviation.

**Table 17.**
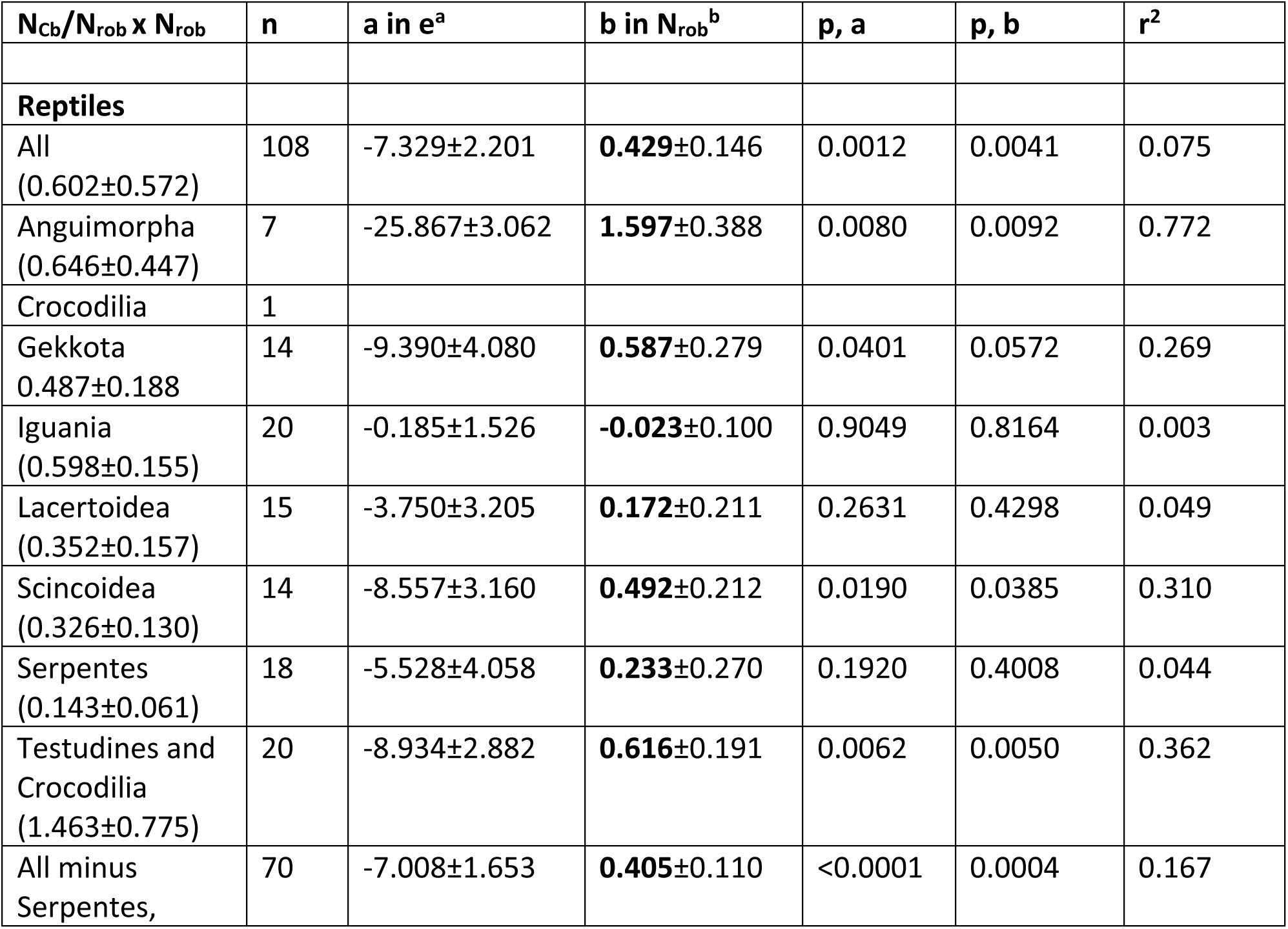
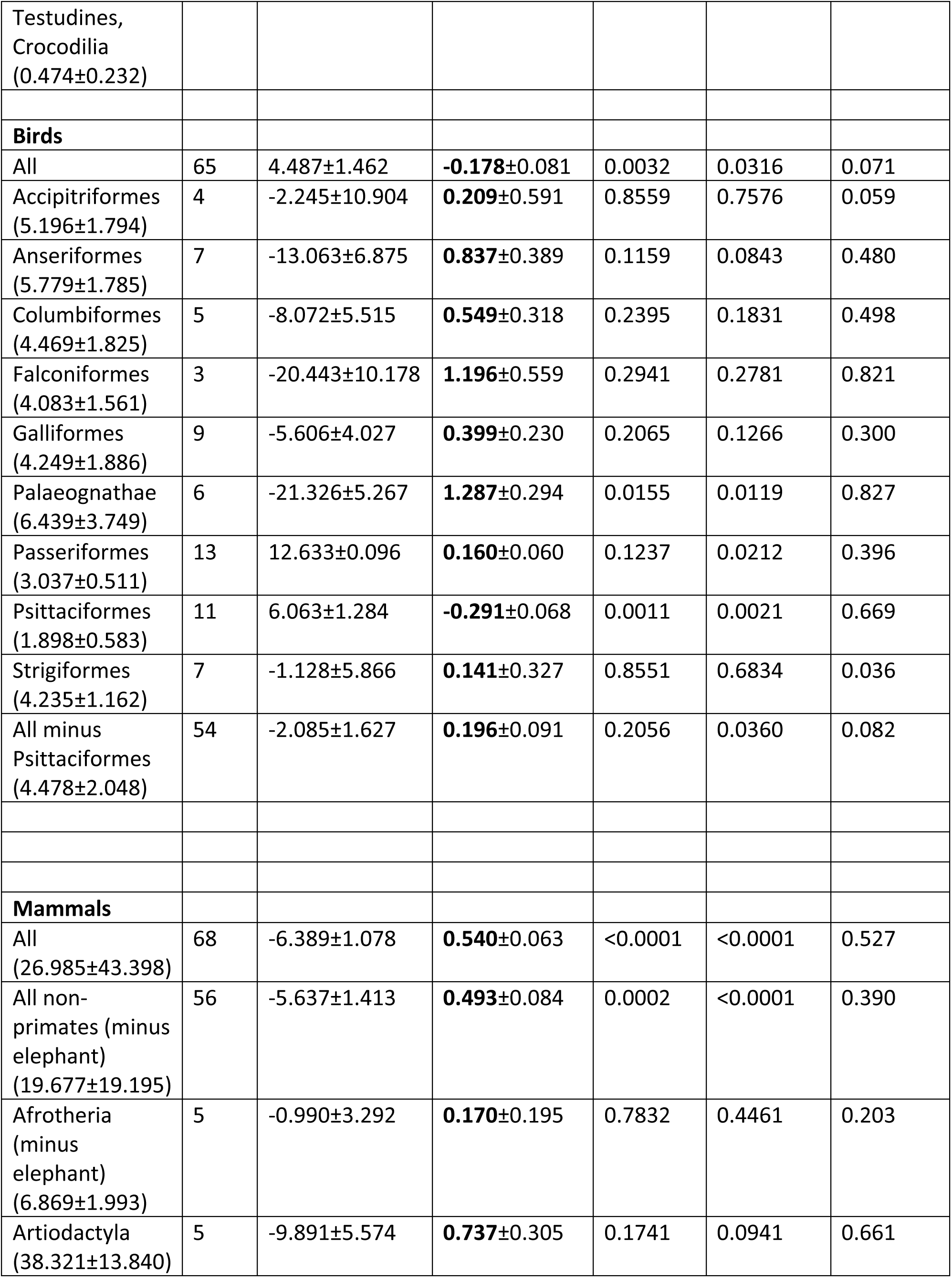
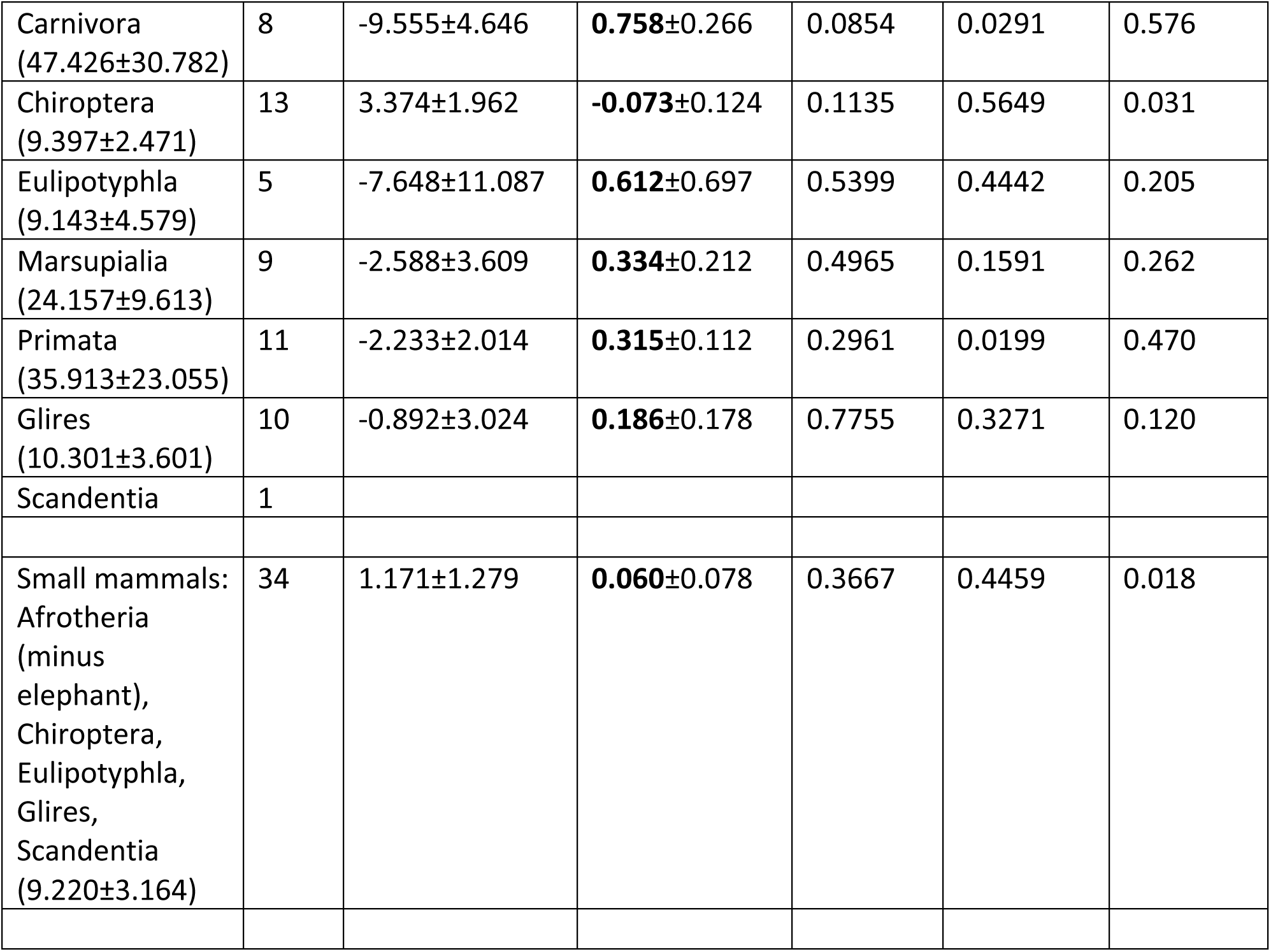
Scaling of the ratio Ncb/Nrob between neurons in the cerebellum (Ncb) and in the rest of brain (Nrob; reptilian or bird brainstem and diencephalon, or mammalian brainstem, diencephalon plus striatum) as a function of numbers of neurons in the rest of brain (Nrob). Values correspond to the equation Ncb/Nrob = e^a±SE^ Nrob^b±SE^, the p-values for a and b, and the r^2^ for the power fit. Values in parentheses are the average ratio Ncb/Nrob for the clade ± standard deviation.

**Table 18.**
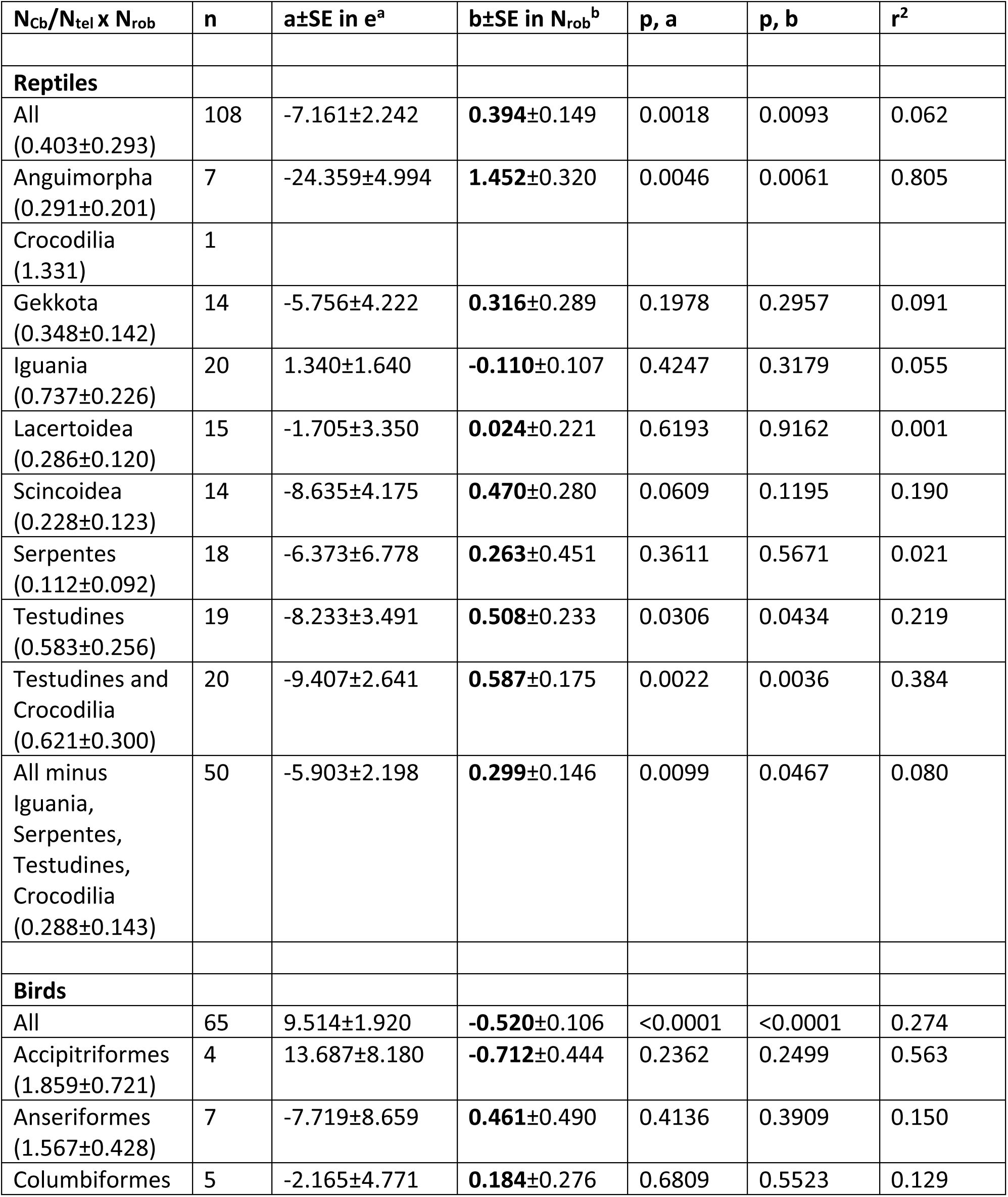
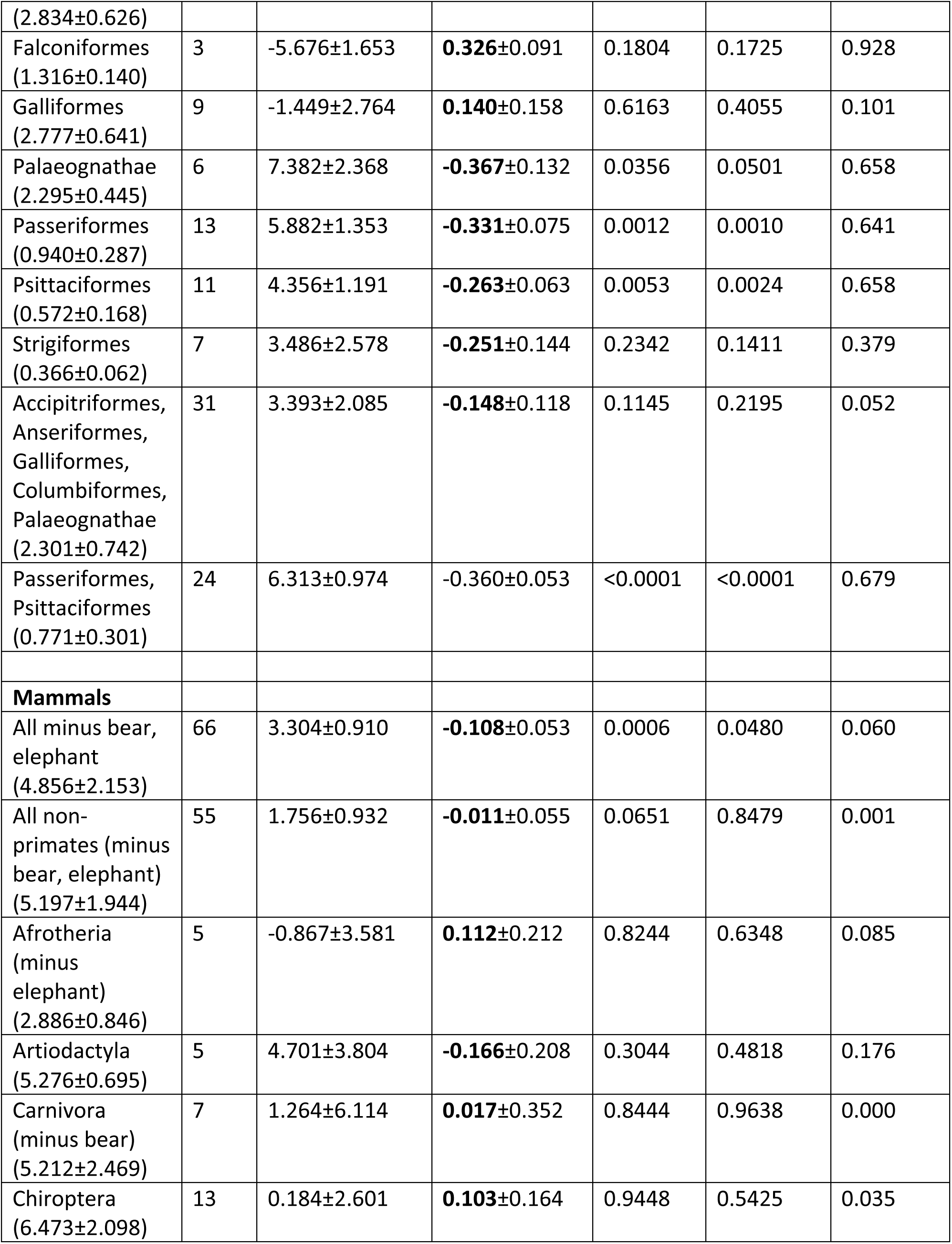
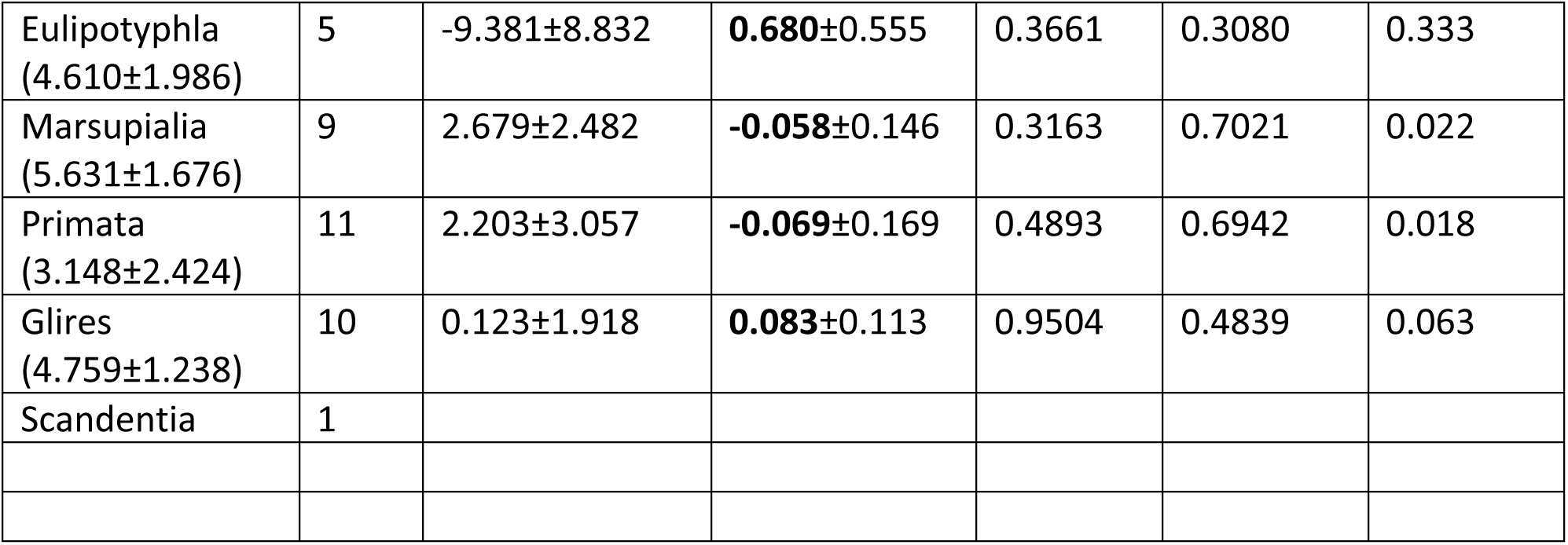
Scaling of the ratio N_cb_/N_tel_ between neurons in the cerebellum (N_cb_) and telencephalon (N_tel_; reptilian telencephalon or bird or mammalian pallium) as a function of numbers of neurons in the rest of brain (N_rob_; reptilian or bird brainstem and diencephalon, or mammalian brainstem, diencephalon plus striatum). Values correspond to the equation N_cb_/N_tel_ = e^a±SE^ N_rob_^b±SE^, the p-values for a and b, and the r^2^ for the power fit. Values in parentheses are the average ratio N_cb_/N_tel_ for the clade ± standard deviation.

Likewise, the shared scaling in mass of the cerebellum with the rest of brain across birds and mammals (Figure 18C), the endothermic amniotes, contrasts with the very diverse relationships between numbers of neurons in the cerebellum and the rest of brain across birds, small mammals, and large mammals (Figure 20C; Table 14). As a consequence, mammals and birds of similar mass ratios between cerebellum and rest of brain (Figure 18D) have ratios of neurons between the two structures that differ by up to a factor of ten, with much larger and variable ratios in mammals than in birds (Figure 20D). Finally, the much larger mass of the telencephalon relative to the cerebellum in reptiles compared to birds and mammals (Figure 18E,F) is no indication of the fact that the proportion of neurons between the cerebellum and the telencephalon (or pallium) is similar between many reptiles and birds (Figure 20E,F; Table 15).

### Mammals are not defined by expansion of pallial neurons over the rest of brain

All reptiles in the dataset gain neurons coordinately between the telencephalon and the rest of brain (Figure 20A), with ratios of numbers of neurons between the structures that vary from just below 1.0 in Iguania to 2.4 neurons in the telencephalon to every neuron in the rest of brain of Testudines and the Nile crocodile (Figure 20B; Table 16). On average, reptiles other than Testudines and the Nile crocodile have 1.4±0.6 neurons in the telencephalon to every neuron in the rest of brain; Testudines and the Nile crocodile stand out amongst reptiles with 2.4±0.6 neurons in the telencephalon to every neuron in the rest of brain (Figure 21, Table 16).

**Figure 21.**
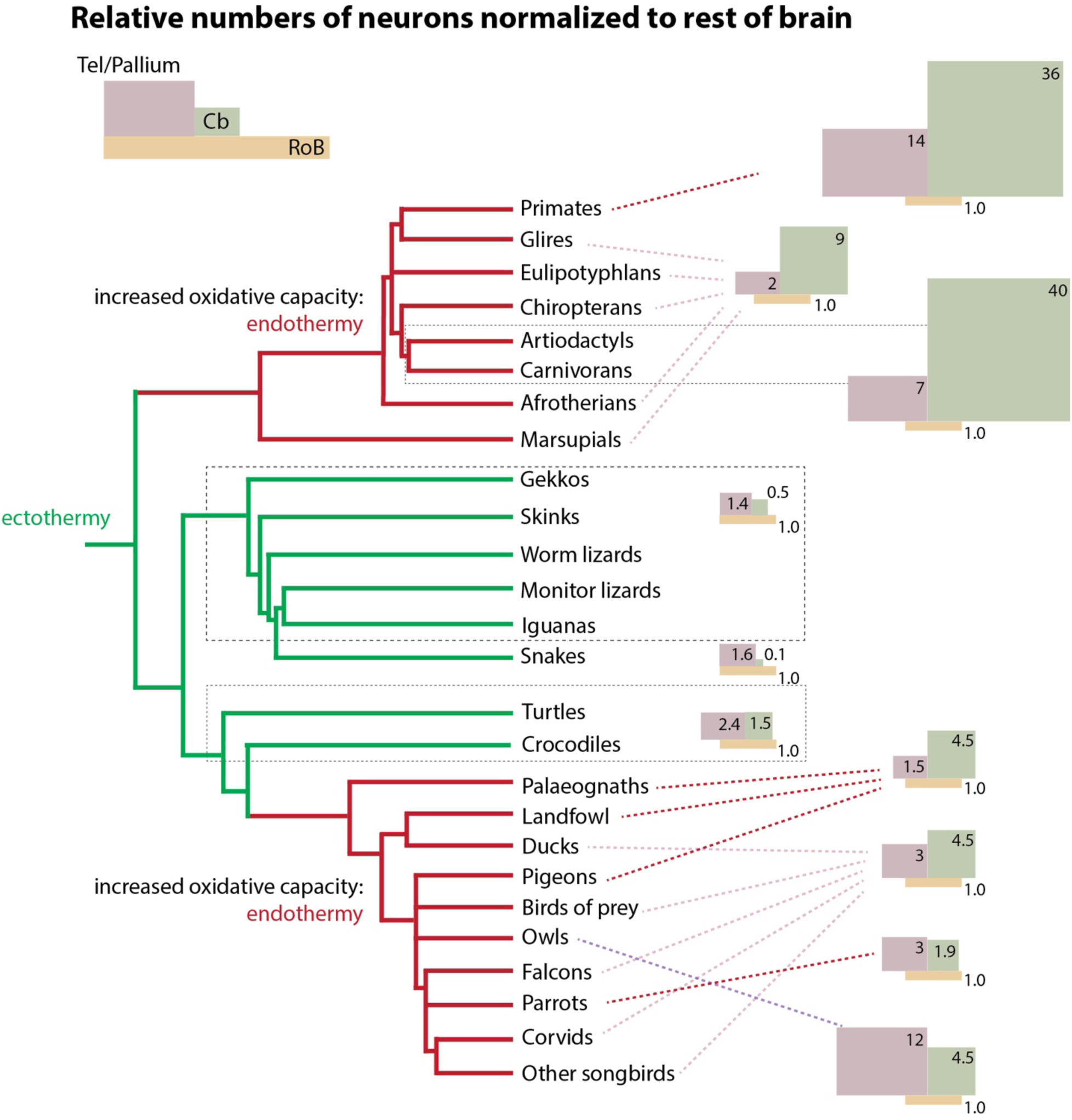
Signature ratios of numbers of neurons between the telencephalon/pallium and rest of brain describe the brains of different amniote clades. The schematics in varying sizes represent the rest of brain (tan rectangle), telencephalon (red) and cerebellum (green). All schematics are thus scaled to a similar number of neurons in the rest of brain; in this way, the surface area of each brain part represents the relative number of neurons in that structure compared to the rest of brain (which has a value of 1.0). The schematics illustrate that reptilian brains are characterized by modestly larger numbers of neurons in the telencephalon compared to the rest of brain, and fewer neurons in the cerebellum (green lines). Birds (pink lines), with increased oxidative capacity, diverged away from reptiles with a step increase to a larger ratio of 4.5 neurons in the cerebellum for every neuron in the rest of brain, followed in most bird clades (other than the early derived Galliformes, as well as Columbiformes) by an increase to on average 3 neurons in the pallium to every neuron in the rest of brain, and reaching a peak in Strigiformes. Independently increased oxidative capacity occurred in mammals with a different brain signature of an increased average of 4 cerebellar neurons for every pallial neuron in all extant mammals, while the ratio of pallial/rest of brain neurons remained low at ca. 2 in small species, but increased in carnivorans, artiodactyls, large marsupials, and reached peak values in primates.

Strikingly, and despite the common narratives of cortical expansion in mammalian evolution (Rowe et al., 2011; Bertrand et al., 2022), mammals do not have the expected matching increase in the ratio of neurons between the cortex and rest of brain that would set them apart from reptiles. Small mammals (eulipotyphlans, chiropterans, rodents, small afrotherians, and South American marsupials) have ca. 2 neurons in the cerebral cortex for every neuron in the rest of brain, as described before (Herculano-Houzel et al., 2014a), which is only slightly more than most reptiles, and fewer than found in turtles and the Nile crocodile (Figure 20B; Figure 21). It is only the large mammals – carnivorans, Australian marsupials, primates, and artiodactyls – that have greatly increased ratios of neurons in the cerebral cortex over the rest of brain, which average just over 7 in artiodactyls and carnivorans, and can exceed 20 in primates (Figure 20B, Figure 21; Table 16). These data show that compared to the ancestor shared with modern reptiles, and to small mammals, there has been a marked expansion of cortical neurons over the rest of the neural tube in the evolution of *large* mammalian species – but not at the origin of mammals.

Interestingly, the early-derived Galliformes and Columbiformes, along with the smallest Palaeognathes, share with reptiles a low ratio of 1.5 neurons in the pallium to every neuron in the rest of brain (Figure 20B, Table 16). In contrast, most bird species in the dataset have a significantly larger ratio of ca. 3 neurons in the pallium to every neuron in the rest of brain (Table 16, Figure 21). The striking outliers are the Strigiformes (owls), with on average 12 neurons in the pallium for every neuron in the rest of brain, on par with primate species in the dataset, and well above passerines and parrots (Figure 21, Table 16), with which owls share a similar range of absolute numbers of pallial neurons (Figure 20A).

The similarities across reptiles, small mammals and the early-derived birds in the ratios in numbers of neurons between the pallium and rest of brain indicate that while the advent of endothermy was associated with vastly increased absolute numbers of neurons in the brain, it did not lead on its own to any relative expansion of “cortical processing power” over the rest of brain (see Herculano-Houzel, 2025). That increase happened later, and separately, in primates, artiodactyls, and carnivorans.

### Expansion of cerebellar neurons over the rest of brain defines endothermic amniotes

The analysis of the scaling of numbers of neurons across structures reveals that the ratio at which neurons are added to the cerebellum over the rest of brain is particularly well conserved across reptilian species, with on average 0.5 neurons in the cerebellum to every neuron in the rest of brain in other reptiles (Figure 20C,D, Table 17). The exceptions are snakes, which have only about 1 neuron in the cerebellum for every 10 neurons in the rest of brain, and, in the opposite direction, Testudines and the Nile crocodile, with 1.5 cerebellar neurons for every neuron in the rest of brain (Table 17; Figure 21).

Compared to other reptiles, birds seem to have experienced a step increase to an average of 4.5±2.0 neurons in the cerebellum to every neuron in the rest of brain (Figure 20D, Figure 21; Table 17). The only exception is Psittaciformes, with lower ratios of 1 to 3 cerebellar neurons to every neuron in the rest of brain, which decrease with increasing numbers of neurons in the latter structure (Table 17).

Mammals as a whole, the other endothermic amniotes, also exhibit an expansion of numbers of cerebellar neurons over the rest of brain, which in this case is even more pronounced than in birds (Figure 20D). Across small mammals, there are on average 9 neurons in the cerebellum for every neuron in the rest of brain; larger mammals have increasing ratios of neurons in the cerebellum for every neuron in the rest of brain, reaching 30 to 40 neurons in the cerebellum to every neuron in the rest of brain in primates, artiodactyls and carnivorans (Figure 21; Table 17).

### Ratios of cerebellar to pallial neurons define clades of amniotes

The cerebellum and pallium develop from entirely different progenitor cell populations in the brain. It is thus not surprising that while numbers of neurons in these two structures tend to vary together across species of reptiles (Figure 20E, Table 18), they occur at no particular ratio: that is, the cerebellum/telencephalon ratio of numbers of neurons is neither shared across reptilian species, nor scales systematically as reptilian brains vary in size (Figure 20F, Table 18). Rather, while almost all reptilian clades have more neurons in the telencephalon than in the cerebellum, different clades have different characteristic ratios of numbers of neurons between the cerebellum and telencephalon, with Serpentes on the low end (with a ratio of 0.1, amounting to almost ten neurons in the telencephalon to every neuron in the cerebellum), Iguania (0.7) and Testudines and Crocodilia (0.6) on the high end (slightly more than one neuron in the telencephalon for every neuron in the cerebellum), and most other reptilian clades in between (Table 18). These ratios are illustrated by the size of the circles in Figure 22.

**Figure 22.**
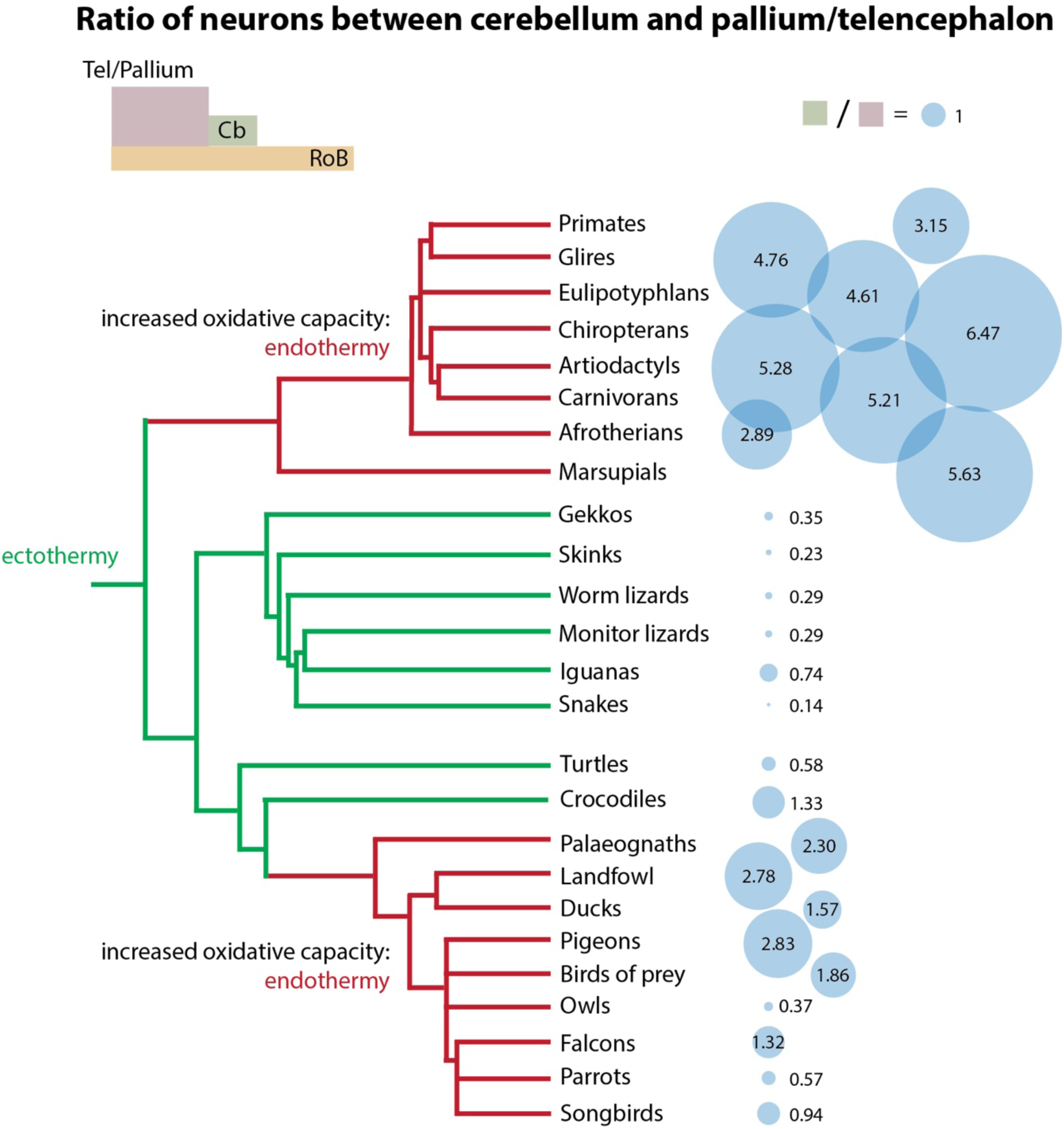
Signature ratios of numbers of neurons between the cerebellum and the telencephalon/pallium describe the brains of different amniote clades. The diameter of the circles in varying sizes represents the ratio of numbers of neurons between the pallium or telencephalon and the cerebellum. The schematics illustrate that reptilian brains, with the exception of Crocodilians, are characterized by ratios smaller than 1, that is, by having more neurons in the telencephalon than in the cerebellum. The endothermic, early-derived birds have around 2 neurons in the cerebellum for every neuron in the pallium, whereas the more recent groups of birds show decreased ratios again, due to the independent expansion of numbers of pallial neurons, which is most extreme in Strigiformes (owls). In contrast, the independently endothermic mammals have a characteristic signature of an increased average of 4 cerebellar neurons for every pallial neuron in all extant mammals, ranging from 3 to 6 (Table 18).

While all birds have in common their increased absolute numbers of neurons in the cerebellum compared to reptiles (Figure 20C), even when matched for cerebellar mass (Figure 14C), different bird clades have characteristic ratios of numbers of neurons between the cerebellum and the pallium. As summarized in Figure 22, early-derived birds have a predominance of neurons in the cerebellum over the pallium, akin to mammals, which is maintained in Falconiformes; Passeriformes have similar numbers of neurons in the cerebellum and pallium; Psittaciformes have a prevalence of pallial over cerebellar neurons; and Strigiformes, like reptiles, have several times more neurons in the telencephalon than in the cerebellum (Table 18). Importantly, the predominance of pallial over cerebellar neurons in Psittaciformes and Strigiformes is due not to a “reversal” to the reptilian pattern, but rather to a relative decrease in numbers of cerebellar neurons in parrots compared to the avian norm given the number of neurons in the rest of brain, and to the separate and striking increase in absolute and relative numbers of neurons in the pallium of owls over the rest of brain (Figure 20).

Mammals, on the other hand, are characterized by a fairly constant average of 3-6 neurons in the cerebellum for every neuron in the cerebral cortex (Figure 22). This is the consequence of the striking similarity between the scaling of numbers of neurons in the mammalian pallium against the rest of brain, and the scaling of numbers of neurons in the cerebellum against the rest of brain (Figure 20F; Table 18).

### Evolution of signature ratios of numbers of neurons in birds and mammals

Taking it all together, each of the three groups of amniotes, and many of their main clades, can be defined by the 3D space they occupy in the relationship between the absolute number of neurons in the rest of brain; the proportion of neurons in the pallium (or telencephalon) over the rest of brain; and the proportion of neurons in the cerebellum over the rest of brain. Strikingly, while the equivalent mass relationships separate reptiles from the endothermic amniotes, they fail to distinguish among birds and mammals (Figure 23). Thus, what clearly distinguishes the two clades of endothermic amniotes from each other and from the ectothermic amniotes is the distribution of numbers of neurons across brain structures. Combined with the increased numbers of neurons that accompanied the evolution of endothermy, the three groups of amniotes can be separated using the diagnostic features illustrated in Figure 23A.

**Figure 23.**
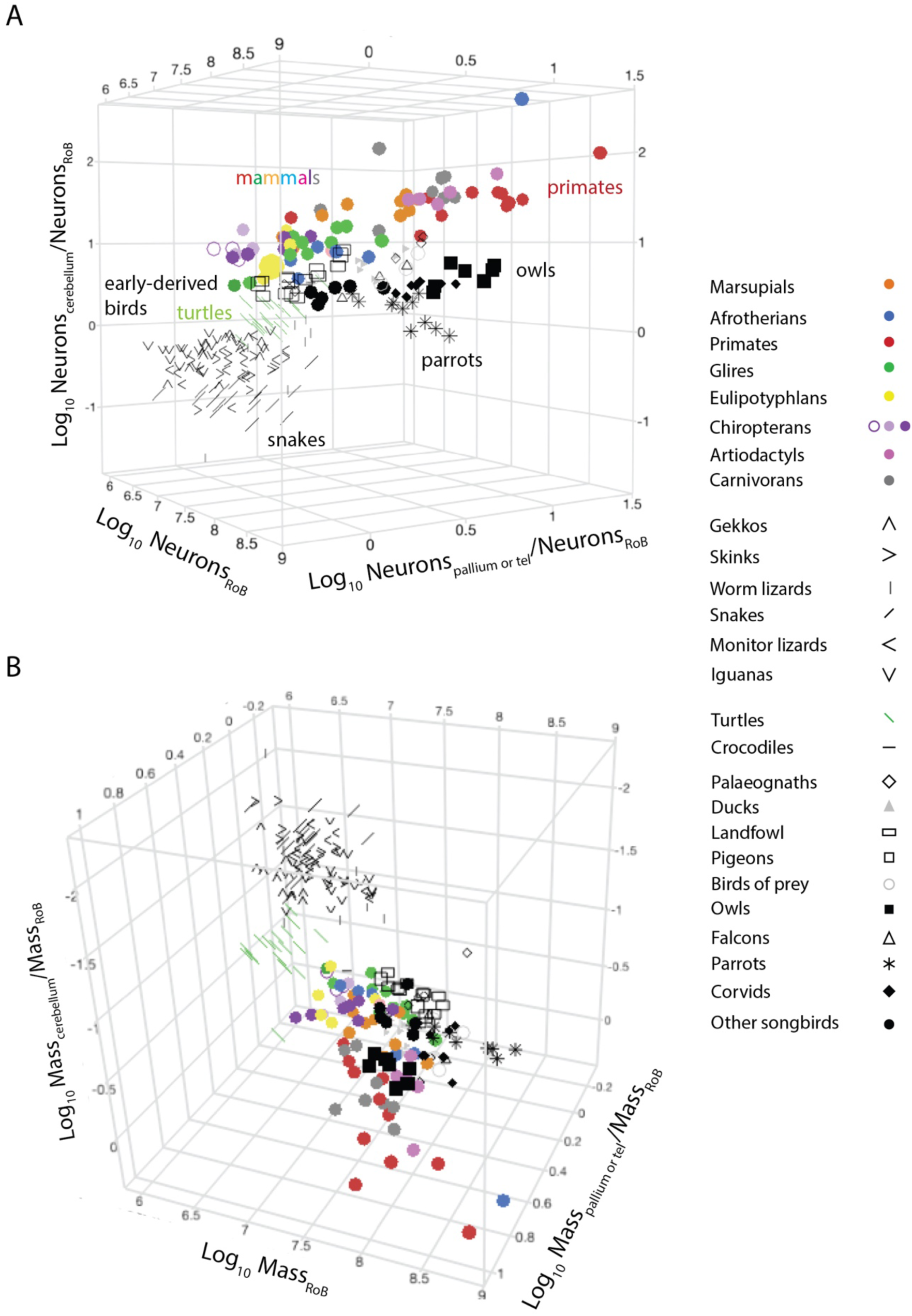
Each group of amniotes, and several clades within them, can be defined by the 3D space they occupy in the relationships between absolute numbers of neurons in the rest of brain and the normalized numbers of neurons in the pallium (or telencephalon) and cerebellum (A). In contrast, the corresponding mass relationships across the structures single out reptiles, but fail to separate birds and mammals (**B**). Each data point corresponds to one species. The key used to depict each clade is shown to the right.

Reptiles are the amniotes with few neurons in the rest of brain as well as proportionately few neurons in the cerebellum and the telencephalon, normalized to the rest of brain. Modern reptiles have brains with fewer than 10 million neurons in the rest of brain, typically half as many in the cerebellum, and a slight (1.4-fold) predominance of neurons in the telencephalon over the rest of brain.

Birds deviate from reptiles with a shift to a higher, non-overlapping range of numbers of neurons in the rest of brain together with a jump to an also higher, non-overlapping range of numbers of neurons in the cerebellum. Compared to the closest living non-avian reptilian relatives, turtles and crocodiles, birds appeared with a 3-fold step increase in numbers of neurons in the cerebellum for every neuron in the rest of brain, shifting to a new ratio of 4.5 neurons in the cerebellum to every neuron in the rest of brain that remains a stable signature of the bird brain. The sole known exception so far is Psittaciformes, in which this ratio drops progressively with increasing numbers of neurons, resulting in an average 1.9 neurons in the cerebellum to every neuron in the rest of brain (Figure 21). Developmentally, the cerebellum is part of the hindbrain (Puelles et al., 2013), which is a key component of the “rest of brain” here. Thus, I propose that birds arose with a modification(s) in the developmental program of the hindbrain that resulted in the generation of 3-fold larger numbers of neurons in the cerebellum. Mechanistically, that could be implemented through two doublings of the numbers of early progenitor cells that give rise to the cerebellum in quantal developmental units, followed by apoptosis (Herrup, 1986).

While the higher cerebellum/rest of brain neuronal ratio remains steady, more recent bird clades are defined by an increased ratio of pallial neurons over the rest of brain, ranging from a reptile-like 1.5 in early-derived birds to 12 in owls, but averaging 3 pallial neurons per neuron in the rest of brain in most bird clades, including Passeriformes and Psittaciformes. As a result, the ratio of neurons between the cerebellum and the pallium is characteristic of each clade of birds (Figure 22). Mechanistically, this means that the pallium, which originates from the dorsal cap of the second neuromere (Puelles et al., 2013), gains neurons fairly independently from both the cerebellum and the rest of brain in birds. This freedom in the addition of numbers of neurons between pallium and cerebellum in bird species is consistent with the lack of major projections linking the two structures in birds, as the bird cerebellum is predominantly connected to brainstem structures instead of the pallium (Gutiérrez-Ibáñez et al., 2018). The lack of major connection pathways between the cerebellum and the pallium in birds together with the shared developmental hindbrain identity of the cerebellum and most of the rest of brain is presumably sufficient to account for the concerted addition of numbers of neurons to the cerebellum and rest of brain across most bird species. It remains to be determined whether the strong link between numbers of neurons formed in the rest of brain and cerebellum of birds is associated with the advent of any major bird-specific neuronal pathways between the two parts of the brain, compared to reptiles, or simply constitutes a quantitative variation, made possible by increased oxidative and cardiovascular capacity in birds, of a qualitatively similar organization to the rest of brain and cerebellum of reptiles.

Birds can thus be defined as the group of amniotes with more than 10 million neurons in the rest of brain and in the cerebellum, with ratios between the two that favor the cerebellum by nearly 2 to 4.5 times. In contrast, the smallest mammalian species still share with reptiles numbers of neurons in the rest of brain below 10 million; what defines mammals is an enormously increased ratio of numbers of neurons in the cerebellum over the rest of brain (Figure 23A), which occurs in coordination with increased numbers of neurons in the cerebral cortex, such that a fairly steady average ratio of 4 neurons in the cerebellum to every neuron in the pallium applies to most species, as described previously.

While the mammalian cerebellum is a hindbrain structure derived from neuromeres r0-r1 exactly like in birds (Puelles et al., 2013), mammals differ from birds in that the former exhibit massive connectivity between the cerebral cortex and cerebellum, mediated by the pontine nuclei of the hindbrain, and back to the cortex through the ventral thalamus (Ramnani, 2006). I thus propose that a key defining innovation at the evolutionary origin of mammals was the formation of the dominant cortico-pontine-cerebellar pathway, reciprocated by cerebello-thalamo-cortical projections, which, combined with the increased oxidative and cardiovascular capacities that allowed for increased numbers of neurons to be supported (as well as for the evolution of endothermy; see the following chapter, Herculano-Houzel 2025), led to the concerted scaling of numbers of neurons in the cerebral cortex and cerebellum that I postulate to define mammals. This hypothesis cannot at present differentiate between the possibilities that increased numbers of neurons in the cortex drove increased numbers of neurons in the cerebellum, or that conversely, increased numbers of neurons in the cerebellum drove increased numbers of neurons in the cerebral cortex. However, since numbers of cerebellar neurons are only established after birth, once adult numbers of neurons in the cerebral cortex have already been attained (Goldowitz and Hamre, 1998), it seems more likely that, if numbers of neurons in one structure drive the formation of proportional numbers of neurons in the others, the driver is the cerebral cortex – which only occurs in mammals, not in birds, for the latter lack strong connectivity between cerebral cortex and cerebellum.

The proposition that the advent of a strong anatomical and functional connection between cerebral cortex and cerebellum lies at the origin of mammalian brains is compatible with the finding that while the expansion of the olfactory bulb is a very obvious and diagnostic development in the evolution of modern mammals (Rowe et al., 2011), it is accompanied by expansion of the lateral hemispheres of the cerebellum (Kielan-Jaworowska et al., 2004). Because the predicted increased in numbers of cerebellar neurons in coordination with increased cerebral cortical neurons at the origin of mammals (Figure 21) is accompanied by an increase in cerebellar mass compared to both the mass of the rest of brain and the mass of the cerebral cortex (Figure 19), relative cerebellar mass defined in this manner can be used as a diagnostic feature of endothermic, mammalian brains. The development of segmentation methods that will make it possible to estimate the volume of the cerebellum in endocasts of fossil species should allow for this new line of investigation to be explored.

### Clade-specific extremes: snakes and owls

While similarities and shared differences across clades reveal patterns that are informative and even diagnostic of what constitutes a reptilian, bird, or mammalian brain, clade-specific features are also enlightening of the mechanisms of biology and evolution. In the present dataset, Serpentes are the most obvious outlier group amongst reptiles, with fewer cerebellar neurons than expected for both their body mass and for their numbers of neurons in the rest of brain. The obvious explanation for the fewer cerebellar neurons than expected for body mass is that Serpentes are characterized by their extremely elongated body, in which case the more biologically correct statement would be that Serpentes have evolved a larger body mass than expected for both the size of their brain structures (Figures 1-4) and their numbers of neurons in the cerebellum and rest of brain (Figure 17). However, the finding that Serpentes do have fewer cerebellar neurons than expected for a generic reptile with their numbers of rest of brain and pallial neurons (Figure 20C-F) calls for a clade-specific feature that would have either driven a reduction of numbers of cerebellar neurons, or allowed it to suffice. One possibility is the loss of limbs, which should lift the imposition on cerebellar processing that otherwise mediates the stabilization of the sensory surfaces of the limbs through fine motor control. This possibility is consistent with the other extreme in neuronal composition that is the cerebellum of the African elephant, which harbors 12 times the predicted number of neurons for a generic mammal, which we have speculated that might be associated to fine sensorimotor processing of the uniquely large appendage, the trunk (Herculano-Houzel et al., 2014b), mediated by a massively enlarged trigeminal ganglion (Pukart et al., 2022). Comparison with other animals of similarly elongated bodies and reduced limbs, such as the caecilian amphibians, should shed light on this possibility. Owls (Strigiformes) are another interesting outlier group, with the largest ratios of pallial neurons per neuron in the rest of brain amongst birds, averaging 12, on par with some primates (Figure 21), which seems to be due to an expansion of the pallium that while remaining in conformity to the neuronal scaling rules that apply to other late-derived bird clades (Figure 11). As a consequence, the exceptionally large ratio of pallial/rest of brain neurons in Strigiformes is reflected in a similarly exceptionally large ratio of pallial/rest of brain mass (Figure 18B). Importantly, whatever the number of neurons in the rest of brain, the large numbers of pallial neurons mean owls have a presumably enlarged signal processing capacity that should be on par with that of crows (Kabadayi et al., 2016), which we have recently showed to be associated with large numbers of associative pallial neurons rather than pallial mass (Ströckens et al., 2022). Indeed, a comparative studies of behavioral flexibility in the form of innovative behavior across the present bird clades confirms that increased numbers of pallial neurons correlate with increased cognitive capacity (Sol et al., 2022).

### Predictions for early reptiles and mammaliaformes

Under the principle of parsimony, the similarities and continuities in the neuronal scaling rules observed between most reptilian and mammalian species must reflect shared scaling rules that also applied to stem amniote species ancestral to synapsids and diapsids (sauropsids) (Figures 11, 15). The data above show that extant reptiles and mammals share not only the neuronal scaling relationships for the telencephalon (Figure 11) and rest of brain (Figure 15), but, importantly for predictive purposes, also the scaling relationship between the mass (Figure 18A) and numbers of neurons (Figure 20A) in these two structures. Additionally, assuming that ancestral amniotes were ectothermic like modern reptiles (Nespolo et al., 2011), it is also reasonable to assume that the scaling rules that apply to the cerebellum of the latter also applied to ancestral amniotes. Thus, using the present findings, one can predict that the neuronal scaling rules that apply similarly to the telencephalon, and to the rest of brain, of modern mammals and reptiles (Figures 11C and 15C), and separately to the cerebellum of modern reptiles (Figure 13C), can be extended to their shared ancestors.

The neuronal scaling rules that putatively applied to ancestral amniotes can be expressed further as a function of brain mass, which allows the estimation of the numbers of neurons that composed those early brains since the advent of high-resolution 3D computerized tomography has allowed for the size of the brain in fossil species to be estimated from their endocasts (Macrini, 2006). Figure 24 shows the neuronal scaling relationships that apply jointly to the cerebral cortex of small extant mammalian species (with brain mass below 100g, excluding primates) and to the telencephalon of extant reptiles excluding turtles and crocodiles. Across these 122 species in the dataset, the number of cortical (or telencephalic) neurons can be predicted from brain mass according to the equation N_tel_ = e^16.609±0.040^ M_br_^0.719±0.021^ (p<0.0001) with an r^2^ value of 0.906 (Figure 24A). That is, over 90% of the variation in the number of cortical (in mammals) or telencephalic (in reptiles) neurons in extant species as distinct as ectothermic reptiles and endothermic mammals can be predicted from brain mass. Similarly, across the same 122 species, the number of neurons in the rest of brain can be predicted from brain mass according to the equation N_rob_ = e^16.087±0.030^ M_br_^0.570±0.016^ (p<0.0001) with an r^2^ value of 0.913 (Figure 24B).

**Figure 24.**
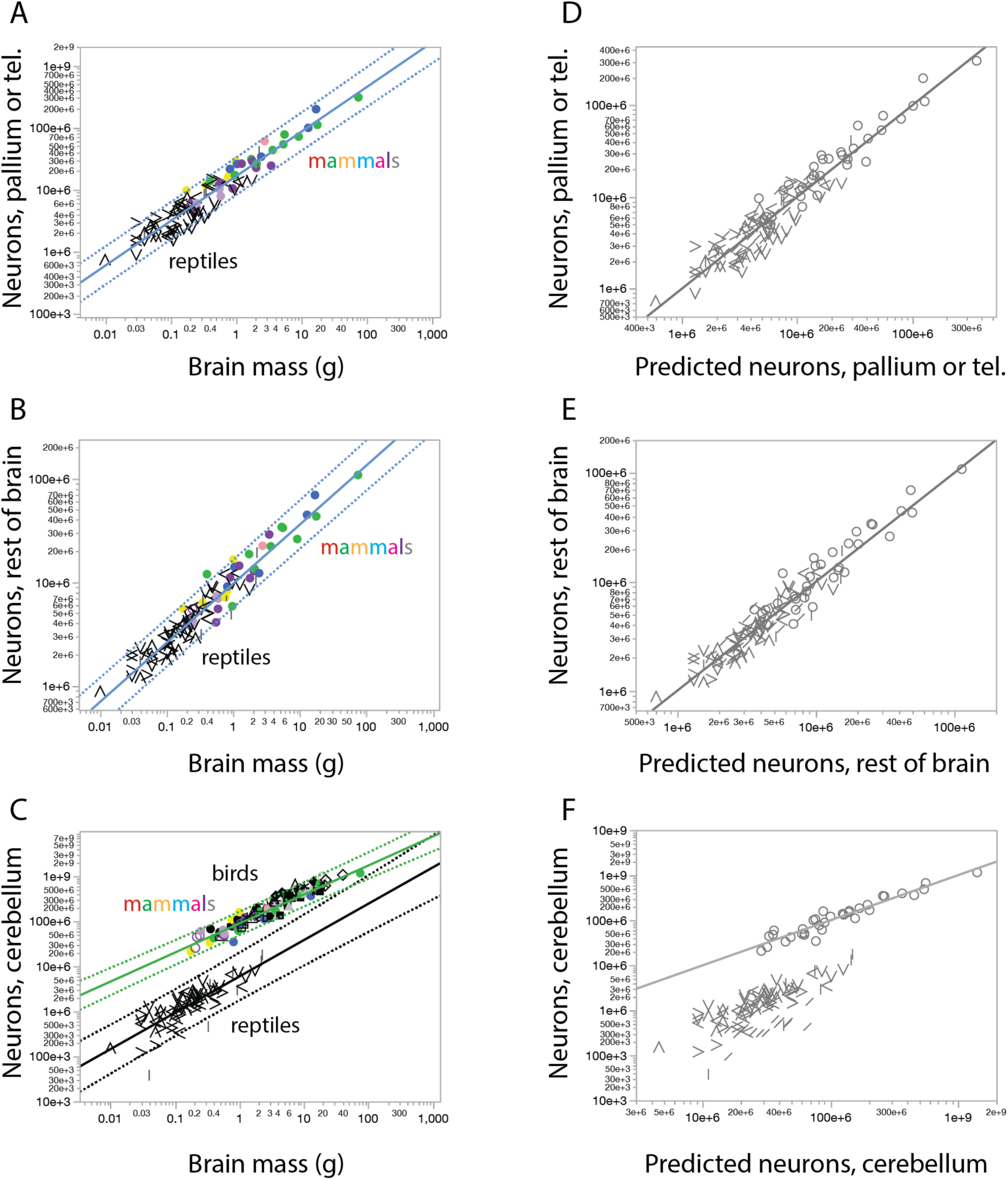
Predictive scaling relationships for numbers of neurons in the different brain structures of ancestral amniotes. A-C, power functions describing numbers of neurons in the telencephalon or pallium (A), rest of brain (B) and cerebellum (C) as a function of whole brain mass (minus the olfactory bulbs). Predictive power functions are Ntel = e^16.609±0.040^ M_br_ ^0.719±0.021^ (r^2^=0.906, p<0.0001), N_rob_ = e^16.087±0.030^ M_br_ ^0.570±0.016^ (r^2^=0.913, p<0.0001) and N_cb_ = e^15.593±0.133^ M_br_ ^0.807±0.064^ (across 70 ectothermic reptilian species, excluding Serpentes, Testudine and the Nile crocodile; p<0.0001, r^2^ = 0.699) or Ncb = e^18.294±0.052^ M ^0.644±0.035^ (across the 34 small mammalian species in the dataset; p<0.0001, r^2^ = 0.913). D-F, comparison between measured and predicted numbers of neurons in the telencephalon (D), rest of brain (E) and cerebellum (F, using the scaling rule for small mammalian species). All graphs are fit with linear functions with slope = 1 and intercept = 0.

In contrast, the relationship between numbers of cerebellar neurons and brain mass differs strikingly between ectothermic reptiles and endothermic mammals (as well as birds). Here, two markedly different scaling relationships apply across 70 ectothermic reptiles (minus Serpentes, Testudine and Nile crocodile) and the mammalian clades comprised of 34 small species in the dataset: N_cb_ = e^15.593±0.133^ M_br_^0.807±0.064^ (p<0.0001, r^2^ = 0.699) and N_cb_ = e^18.294±0.052^ M_br_^0.644±0.035^ (p<0.0001, r^2^ = 0.913), respectively (Figure 20C). The relationship between predicted and measured numbers of neurons across the same species is depicted in Figure 20D-F, which shows that brain mass can be used to make accurate predictions of numbers of neurons in the main divisions of the brain of living reptiles and small mammals.

It follows from the excellent fit between actual numbers of brain neurons and estimates predicted from brain mass (Figure 24D-F) using the scaling functions above that the numbers of neurons that composed the telencephalon and rest of brain of early amniotes can be predicted directly from their brain mass estimated from reconstructed endocasts of fossil skulls. As to the cerebellum, however, the clear shift in the neuronal scaling rules that apply between mammals and reptiles means that inferring the neuronal scaling rules that applied to early mammaliaformes, which may or may not have already been endothermic, requires further evidence to infer which neuronal scaling rules applied: whether the allometry that describes ectotherms like the modern reptiles, or endotherms like modern mammals. Importantly, which allometric rules applied to the cerebellum of fossil early amniote species can be inferred from the relationship between body mass and cerebellar mass (Figure 3) and/or by the different scaling relationship between the mass of the cerebellum and that of the whole brain (Figure 6) across extant reptiles and mammals. For the sake of the current estimates, I will use the assumption that early mammaliaformes, with their relatively enlarged cerebella (Kielan-Jaworowska et al., 2004), already had numbers of neurons that conform to the scaling rules that apply to modern, endotherm mammalian species.

The equations above, depicted in Figure 24, can be used to predict the numbers of neurons that composed the brain of early amniote species by applying them to data on estimated brain volume (assumed to equal brain mass, using a specific density of brain tissue of 1 g/cm^3^) from endocast reconstructions that excluded the olfactory bulbs in mammaliaformes, or that had barely distinguishable olfactory bulbs in cynodonts and other early Therapsid taxa. Brain mass and body mass values in fossil mammaliaformes and stem amniote species were compiled from the literature (Benoit et al., 2017a,b; Boonstra, 1968; Laas, 2015; Castanhinha et al., 2013; Laass and Kaestner, 2017; Jerison, 1973; Hopson, 1979; Rowe et al., 2011; Sigurdsen et al., 2012; Rodrigues et al., 2014; Quiroga, 1980; Luo et al., 2001; Macrini, 2006; Macrini et al., 2007; Bertrand et al., 2019, 2020, 2022; Cameron et al., 2018; Maugoust and Orliac, 2021), and are listed in Table 18. The results are shown in Figure 25A-C, plotted against estimated body mass for each species solely for illustration purposes. Analysis of the estimated numbers of neurons that composed the brains of early amniotes indicates, for a given body mass, a clear shift from modern reptile-like absolute numbers of neurons in the telencephalon (Figure 25A), cerebellum (Figure 25B) and rest of brain (Figure 25C) of early therapsid species in the Permian and Triassic to modern mammal-like numbers of neurons in the Eocene, with intermediate numbers in species living during the Cretaceous and Paleocene. Of note, *Hadrocodium wui*, considered the earliest mammaliaform (Luo et al., 2001), from the Jurassic period, had an estimated 14.03 million neurons in the brain, 1.6 million neurons of which located in the telencephalon, fewer than in living mammals and on par with the similarly small Sumatran flying dragon (*Draco sumatranus*, which however has far fewer neurons in the cerebellum and thus in the brain as a whole, barely exceeding 5 million; Kverkova et al., 2022). *Morganucodon*, a larger mammaliaform that also lived during the Jurassic, had an estimated 52.32 million brain neurons, 6.9 million of which in the telencephalon, comparable to extant microbats in the dataset – although *Morganucodon* had a body mass estimated at ca. 5 times the body mass of small extant bats with similarly small numbers of cortical neurons (Herculano-Houzel et al., 2020). The largest Jurassic mammaliaform in the dataset, *Triconodon mordax*, had the body mass of a hamster but the predicted number of cortical neurons, 13 million, found in a laboratory mouse of ¼ the body mass (Herculano-Houzel et al., 2006).

**Figure 25.**
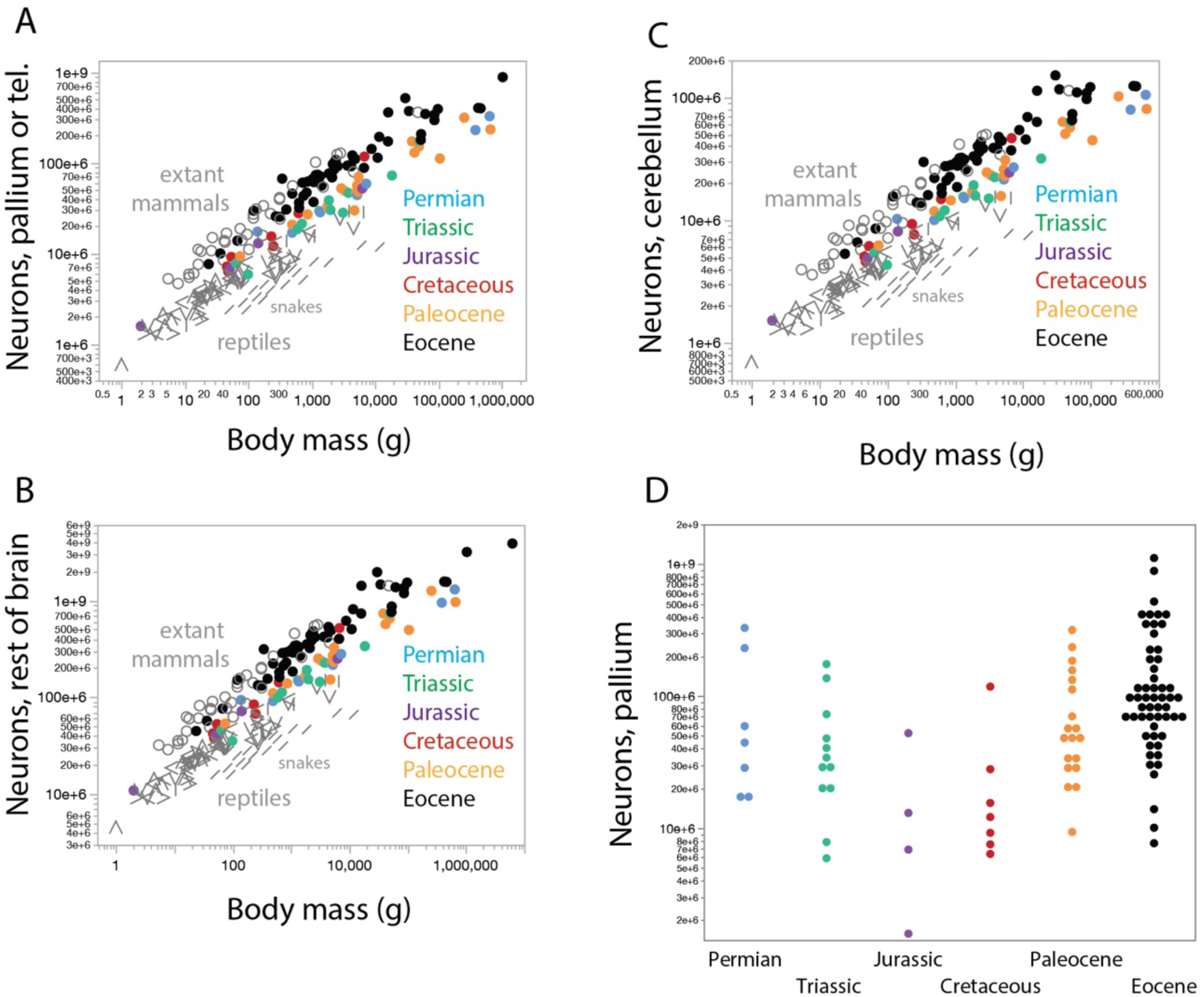
Predicted neuronal composition of the brains of fossil mammaliaform species. Note that for a given body mass, numbers of neurons in all parts of the brain (telencephalon or pallium, **A**; cerebellum, **B;** rest of brain, **C**) increase from the earliest species in the Permian/Triassic to the later species in the Eocene. **D,** In contrast, when body mass is not a confounding variable, the range of the distribution of inferred numbers of pallial neurons in fossil therapsid species appears to decrease from Permian to Jurassic periods, then increase progressively until the Eocene. Data listed in Table 19.

**Table 19.**
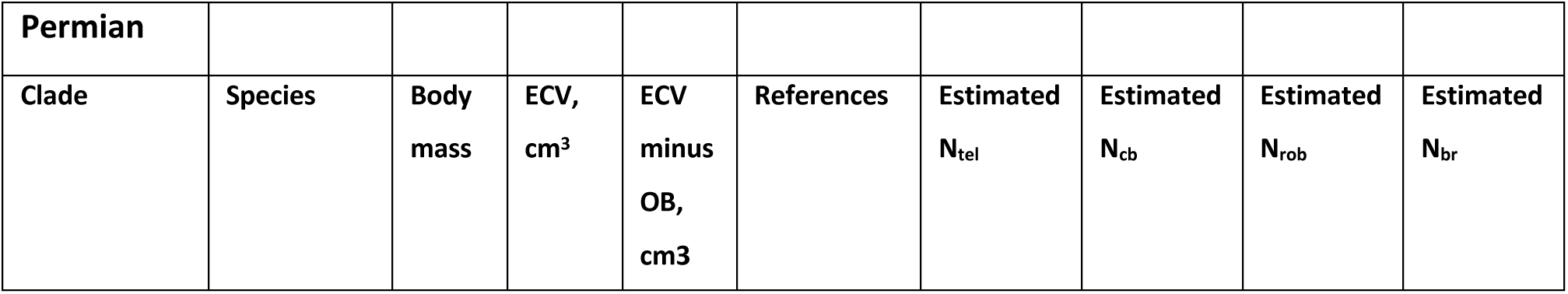
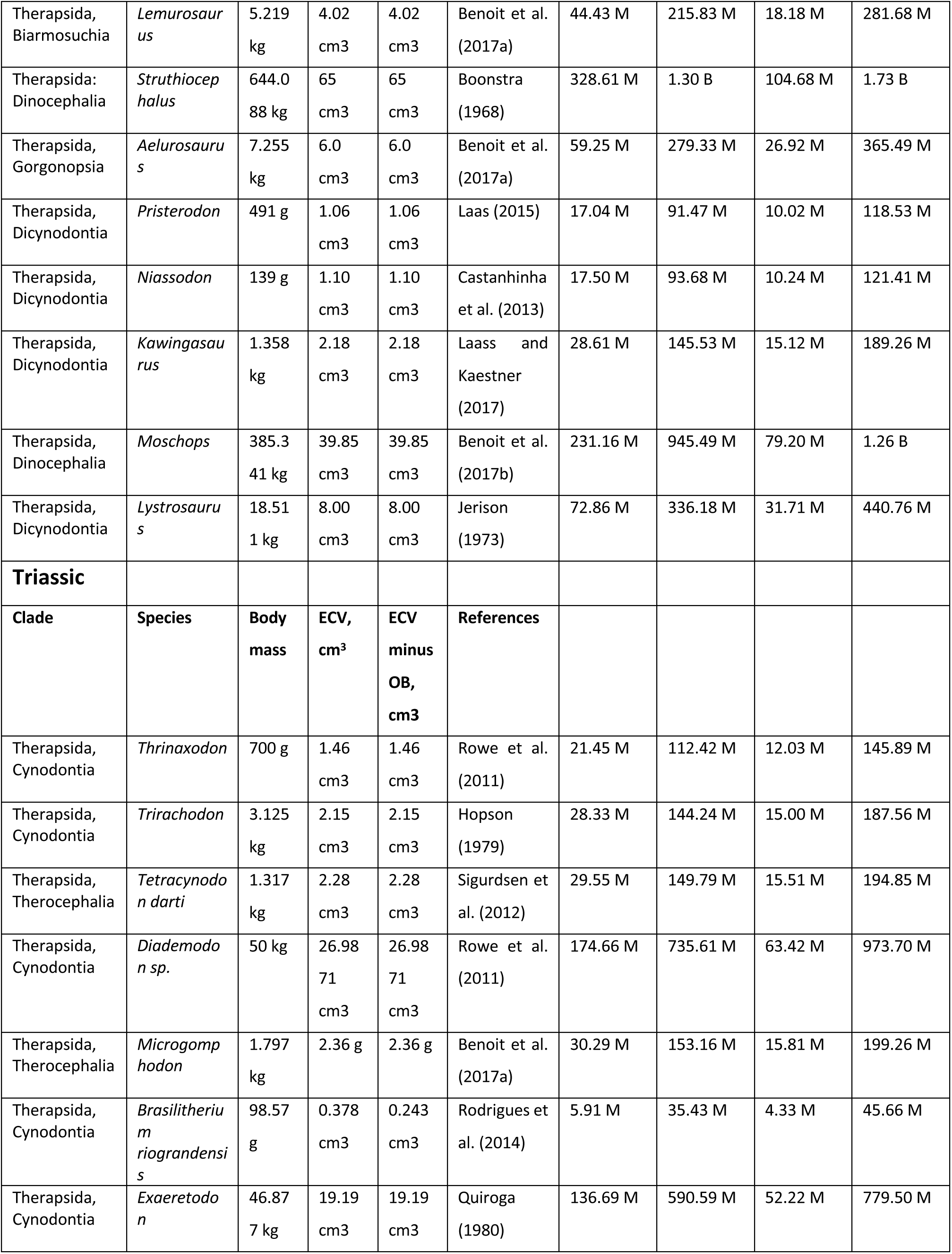
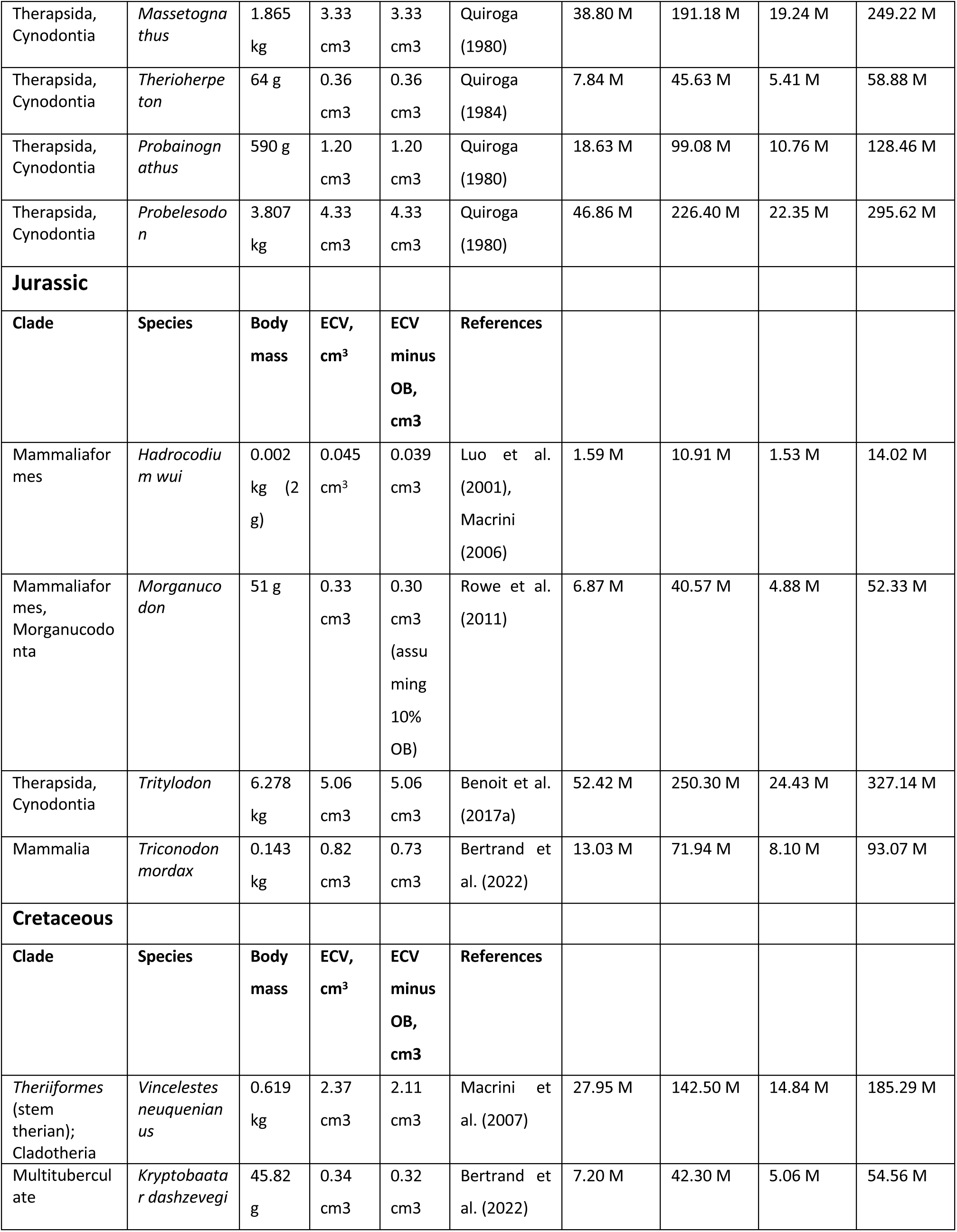
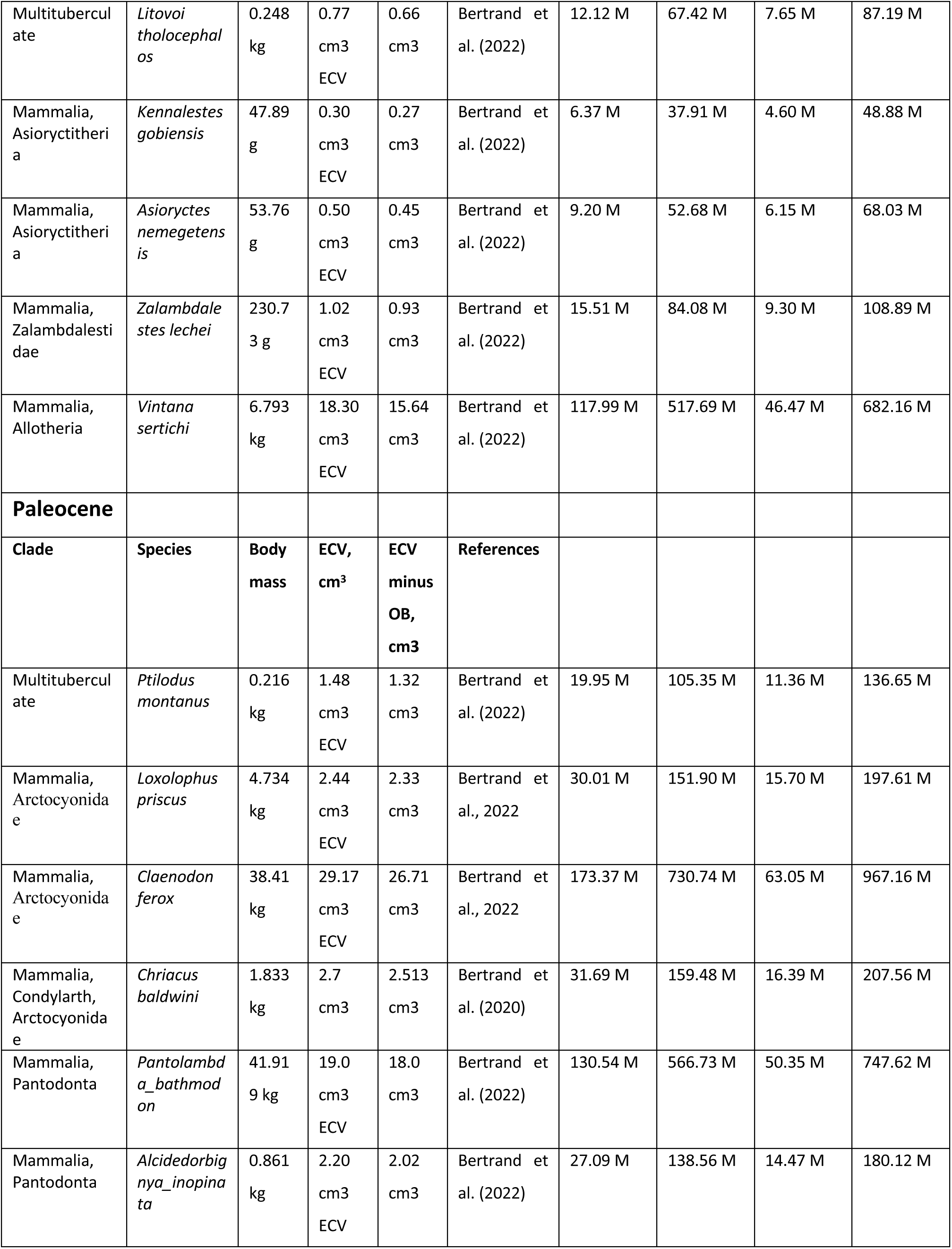
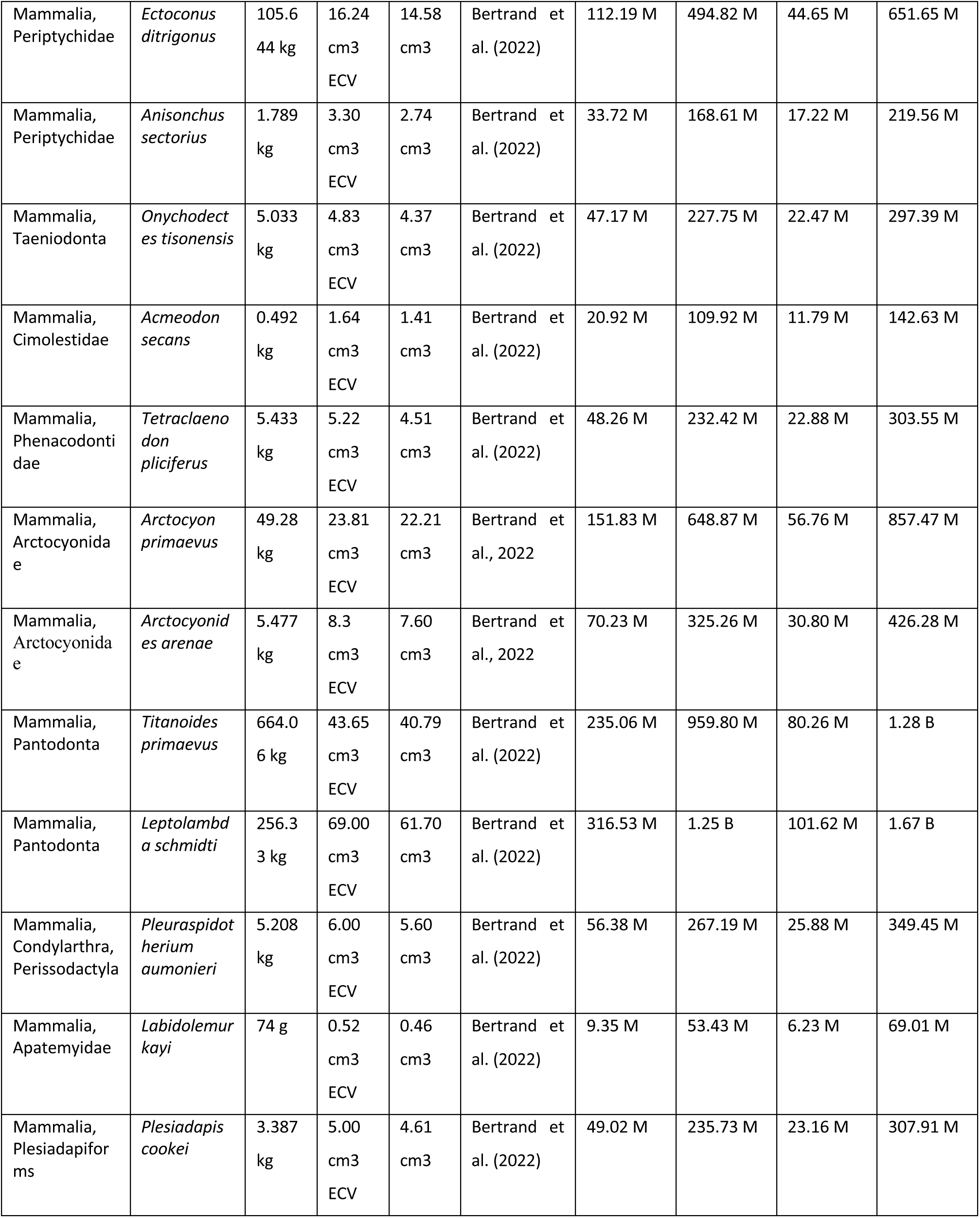
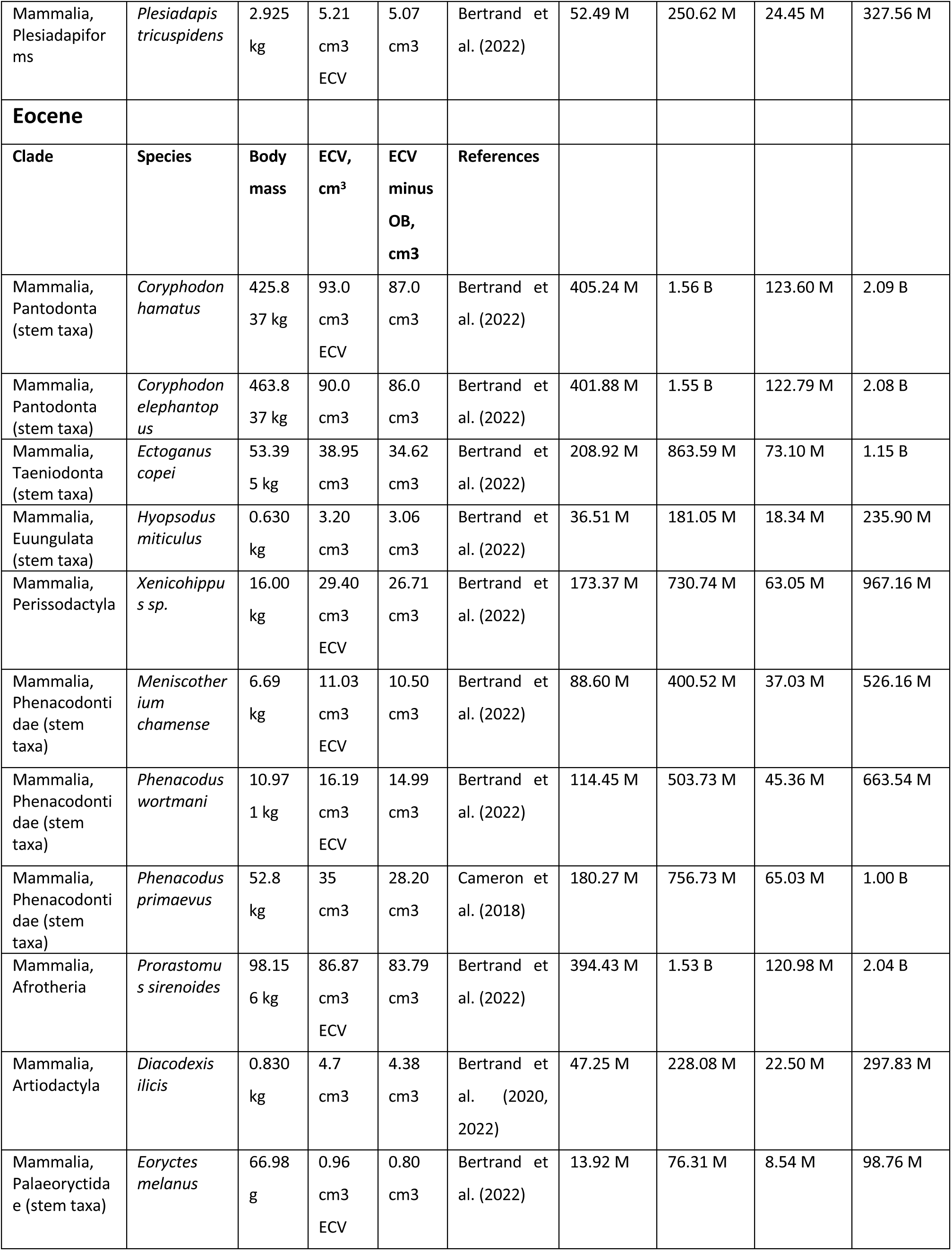
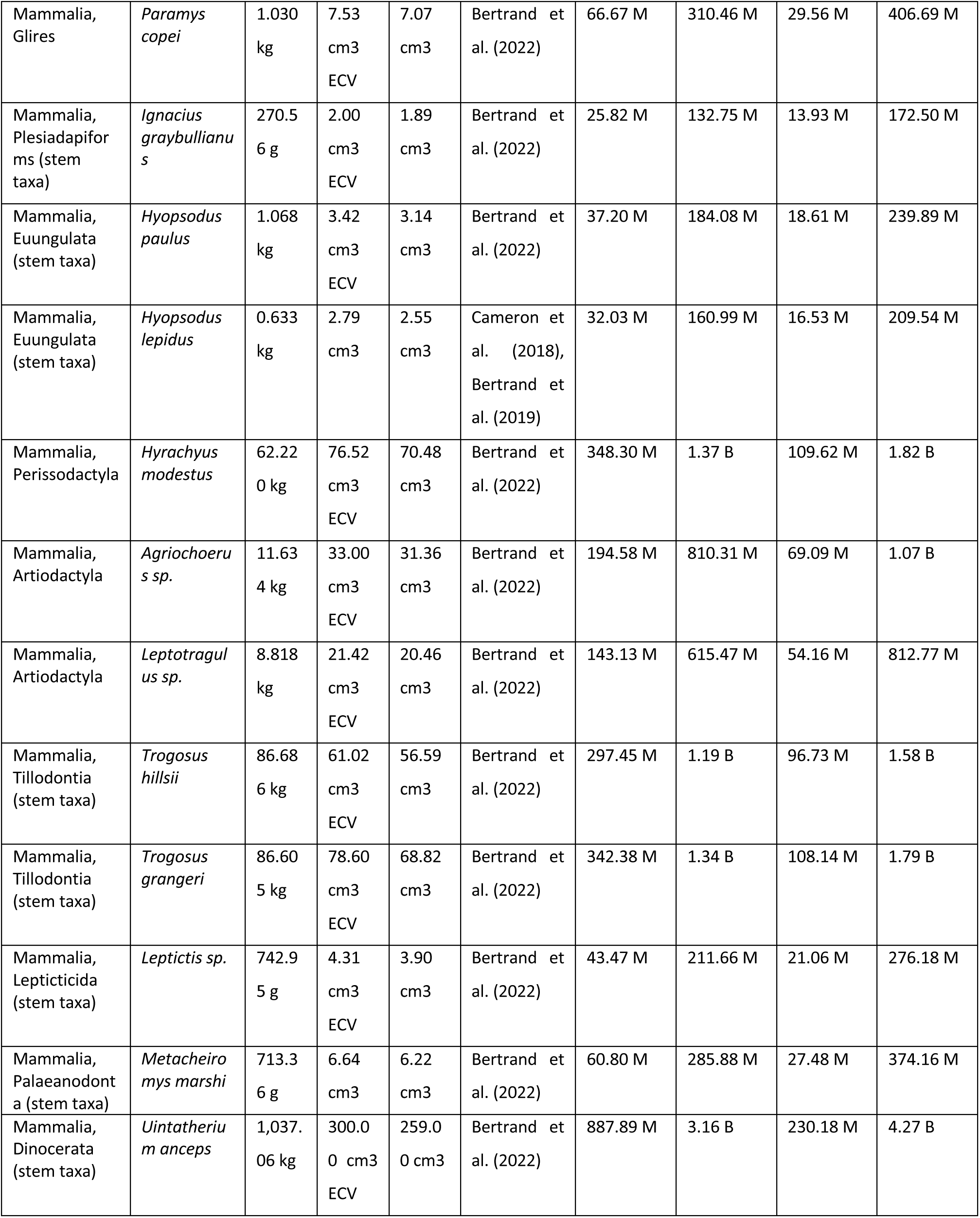
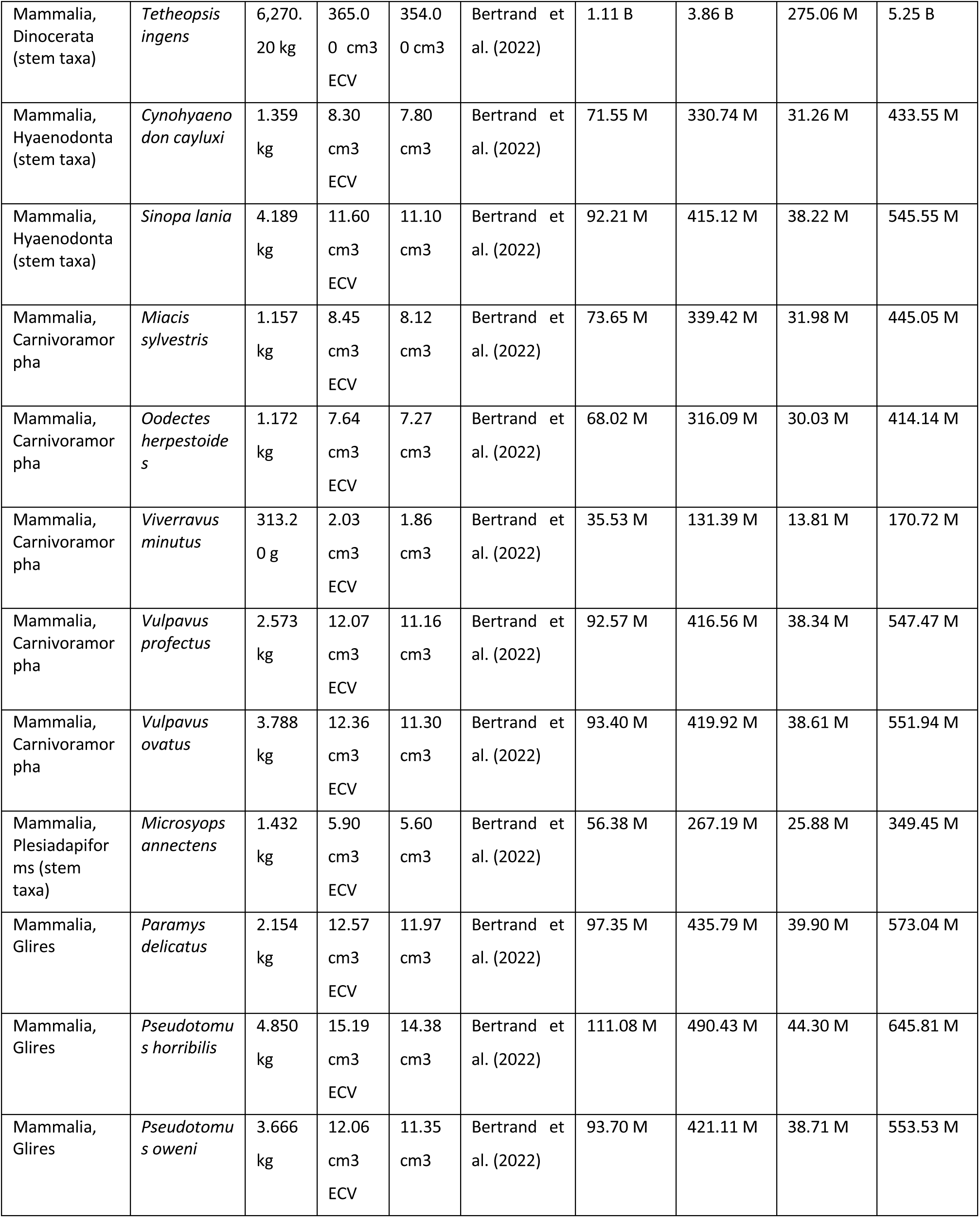
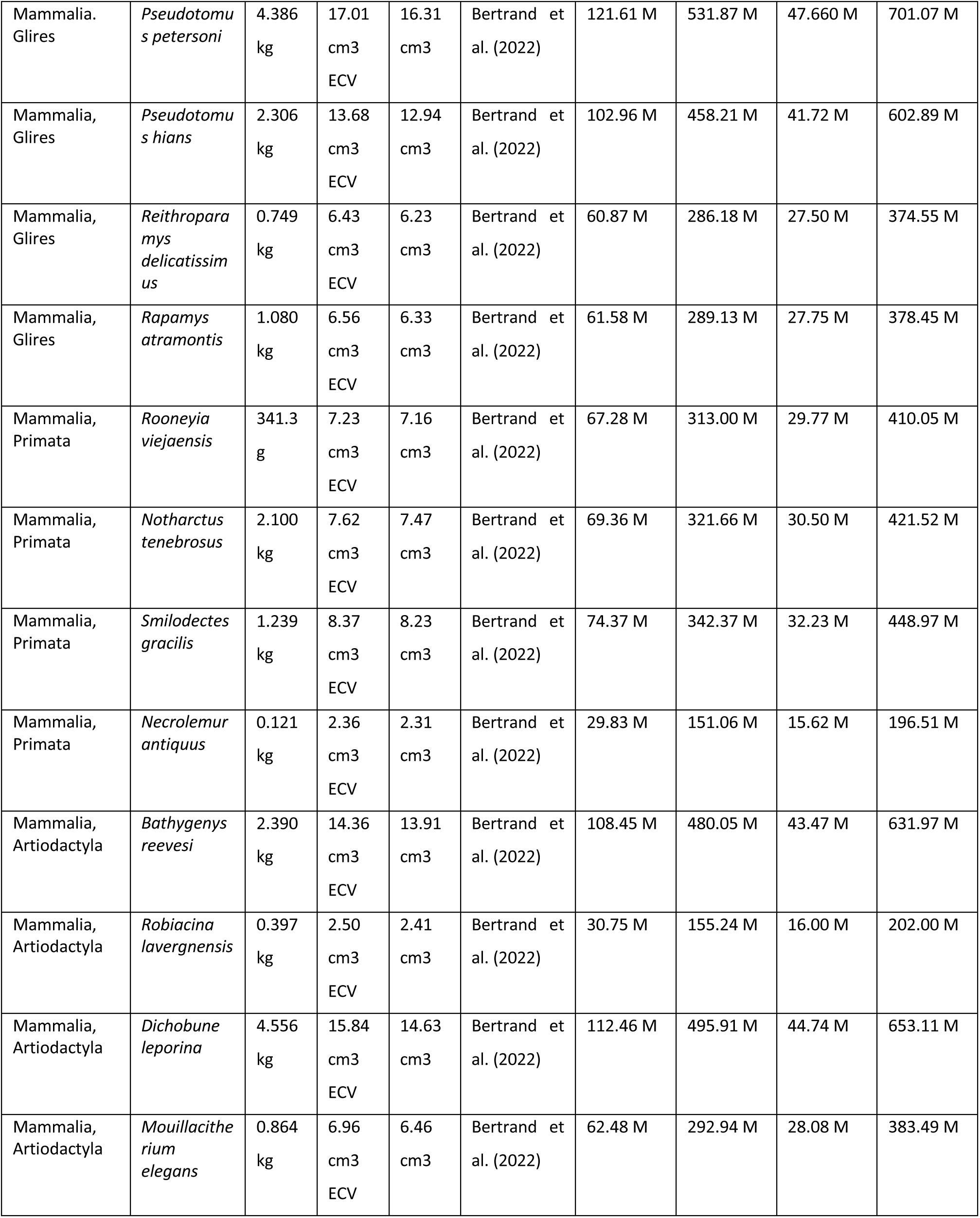
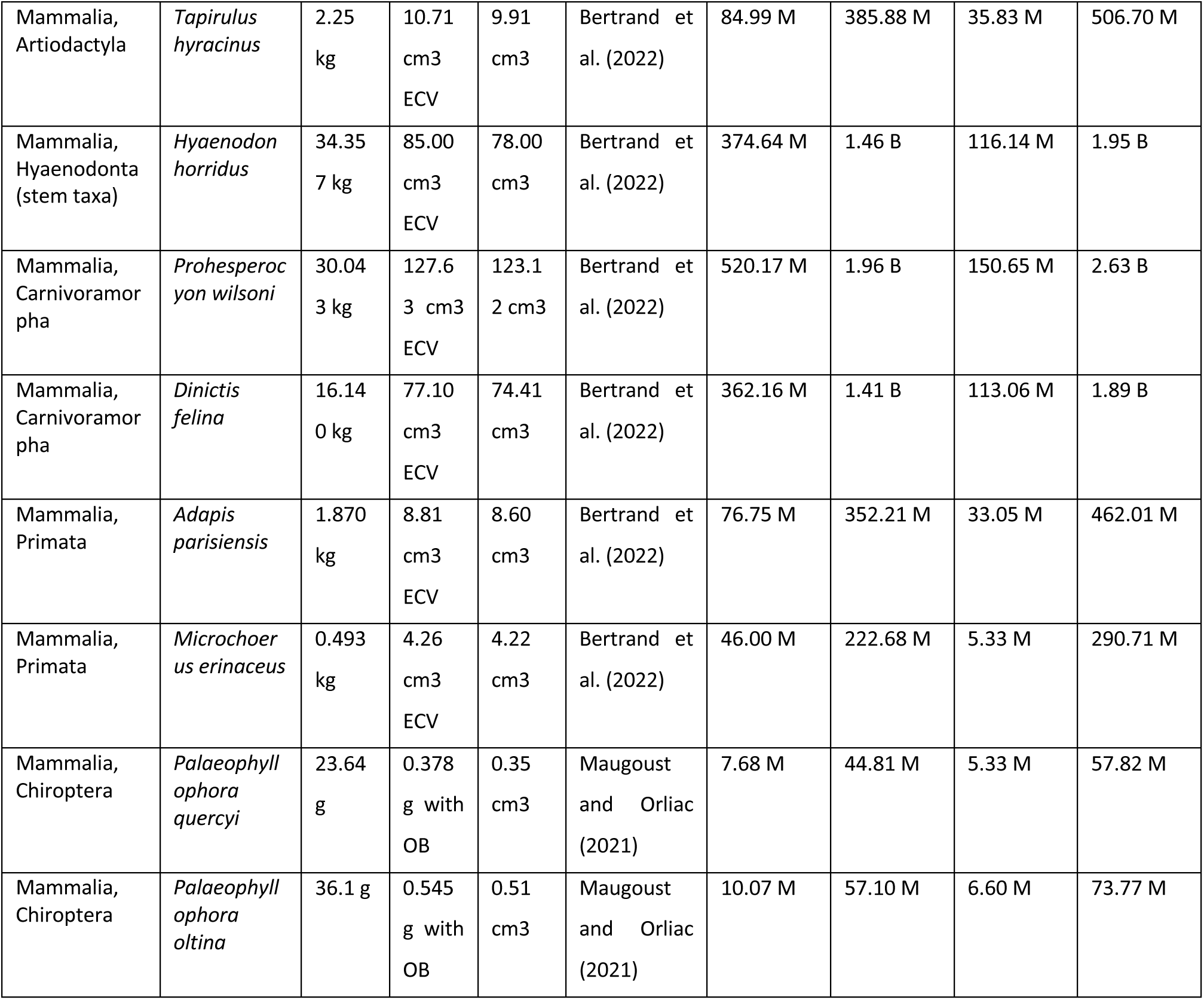
Numbers of neurons in the three parts of the brain of fossil Therapsids: telencephalon (N_tel_), cerebellum (N_cb_), and rest of brain (N_rob_), estimated from reported brain mass (M_br_, minus olfactory bulbs; in grams) inferred from cranial endocasts according to the equations N_tel_ = ^e16.609±0.040^ M_br_^0.719±0.021^ (p<0.0001, r^2^=0.906), N_cb_ = e^18.294±0.052^ M_br_^0.644±0.035^ (p<0.0001, r^2^ = 0.913, calculated for extant mammals) and N_rob_ = e^16.087±0.030^ M_br_ ^0.570±0.016^(p<0.0001, r^2^=0.913). Brain mass in grams was calculated from endocranial volume (ECV) in cm3 assuming 1 cm^3^ = 1 g. M, million neurons; B, billion neurons. N_br_, estimated total number of brain neurons, calculated as the sum N_tel_ + N_cb_ + Nr_ob_.

In Cretaceous therapsids of known brain volume, the range of numbers of cortical neurons is expanded compared to Jurassic species: while the lower values remain at 6-7 million cortical neurons (in the multituberculate *Kryptobaatar* and the mammal *Kennalestes*), the upper values extend to 118 million cortical neurons in the allotherian *Vintana sertichi*, comparable to the 111 million cortical neurons found in the modern agouti *Dasyprocta primnolopha*, although the Cretaceous animal weighed 6.8 kg and the modern agouti, only 2.8 kg (Herculano-Houzel et al., 2006; Table 18). In the Paleocene, the upper end of the range expanded further to reach 316.5 million cortical neurons in the pantodontan *Leptolambda schmiditi*, of 256 kg body mass (Table 17) – similar to the 306 million cortical neurons found in the much smaller modern capybara, at ca. 47 kg (Herculano-Houzel et al., 2006).

Eocene mammals attained even larger numbers of cortical neurons in bodies that were closer in size, but still larger, than modern mammals of similar numbers of cortical neurons. The largest estimated number of cortical neurons in the dataset belongs to *Tetheopsis ingens*, of the stem mammalian taxon Dinocerata, with 1.1 billion cortical neurons – about 1/5 the number of cortical neurons of the African elephant (Herculano-Houzel et al., 2014) - in a body of 6 tons (Table 19), even larger than the 4-5 tons of the modern African elephant. In contrast, the Eocene mammals with the fewest cortical neurons, the chiropterans *Palaeophyllophora quercyi* and *P. oltina*, had an estimated 8-10 million cortical neurons in bodies of 24-36 g, on par with the modern microchiropteran *Cardioderma cor*, with 11 million cortical neurons in a body of 26 g (Herculano-Houzel et al., 2020). All estimates are listed in Table 19.

Taken together, the numbers obtained here indicate that the earliest mammals of the Jurassic had fewer telencephalic neurons than the smallest modern species, but over time, both the lower and the upper ends of the range progressively increased, reaching modern numbers of neurons for body mass in the Eocene. In comparison, the modern range of mammalian body masses is already found in fossils of the Paleocene, showing that mammalian evolution was characterized by “brawn before brains”, and not by a concerted increase in body and brain masses (Bertrand et al., 2022). However, while that study suggested that larger brains evolved later under positive selective pressure because of the cognitive advantage that they posed, I favor an alternate interpretation, focused on possibilities and opportunities. The analysis of numbers of telencephalic neurons estimated by brain mass untainted by body mass, shown in Figure 25D, indicates that large brains with moderately large numbers of telencephalic neurons similar to those found in modern cats and dogs already occurred in Permian therapsids such as *Moschops* (231 million neurons) and *Struthiocephalus* (329 million neurons; Table 19). The ensuing decrease in brain mass and range of estimated numbers of telencephalic neurons towards an apparent minimum in the Jurassic period (driven by the diminutive Jurassic fossil *Hadrocodium*; Rowe et al., 2011) coincides with the rise of large dinosaurs that came to dominate land habitats in the Mesozoic (Brusatte, 2018), although the small range and low values could be an artifact of the small sample size. Still, it seems more parsimonious that the factors behind what range of numbers of brain neurons and brain mass were found at any period were first and foremost biological possibility (for instance, depending on rate of energy intake, which increases together with body mass; Fonseca-Azevedo and Herculano-Houzel, 2012) and opportunity (which depends on environmental conditions and competition with other species), before any considerations of “need” or “advantages”. Thus, I favor a data-based account of mammalian evolution in which moderately large, modern mammal-like numbers of brain neurons, and thus brain mass, were already viable very early, in large Permian endothermic species that could ingest proportionately large amounts of food; possibly decreased in early therapsids as these species faced the rise of dinosaurs, and as smaller mammals, which could only afford small numbers of brain neurons, had a survival advantage over larger mammals; then increased back once larger-bodied mammals became once again *viable* (not more advantageous) after the K-Pg mass extinction that obliterated dinosaurs (Alvarez et al.,1980; Brusatte, 2018). In this scenario, brains with more neurons evolved not because they were advantageous (for species with fewer neurons, that is, reptiles and small mammals, were never replaced out of existence), but simply because they were finally possible, thanks to the combination of physiological changes that also allowed endothermy, and to the energetic and ecological possibilities opened up by the demise of the dinosaurs. The association between physiological changes that allowed endothermy and the advent of increased numbers of neurons in the brains of mammals and birds are explored in the following chapter (Herculano-Houzel, 2025).

**Table 19** Numbers of neurons in the three parts of the brain of fossil Therapsids: telencephalon (Ntel), cerebellum (Ncb), and rest of brain (Nrob), estimated from reported brain mass (M_br_, minus olfactory bulbs; in grams) inferred from cranial endocasts according to the equations N_tel_ = e^16.609±0.040^ M_br_ ^0.719±0.021^ (p<0.0001, r^2^=0.906), Ncb = e^18.294±0.052^ M_br_ ^0.644±0.035^ (p<0.0001, r^2^ = 0.913, calculated for extant mammals) and N_rob_ = e^16.087±0.030^ M ^0.570±0.016^ (p<0.0001, r^2^=0.913). Brain mass in grams was calculated from endocranial volume (ECV) in cm^3^ assuming 1 cm^3^ = 1 g. M, million neurons; B, billion neurons. N_br_, estimated total number of brain neurons, calculated as the sum N_tel_ + N_cb_ + N_rob_.

### On disregarding body mass

I have made a point in this study to ignore body mass, even though it is obviously a relevant variable in terms of ecology, energy consumption and physiology. Part of the reason is that the original analysis of the same dataset already focused on relationships between brain mass and numbers of neurons to body mass (Kverkova et al., 2022), and established that birds and mammals evolved with increased numbers of brain neurons for their body mass compared to reptiles and ancestral species. Most importantly to this author, however, is the reiterated demonstration that there is no obligatory scaling relationship between body mass and number of neurons in any brain structure, specially the rest of brain, which directly operates the body (Burish et al., 2010; Herculano-Houzel et al., 2015a,c; Ngwenya et al., 2016). In this case, then body mass cannot be used as a universal predictor of brain scaling, and its use as a normalizer is unwarranted and should be discontinued, for there is much more information to be gained in the direct analysis of variables pertaining to brain size and composition without contamination by the confounding factor that is body mass. Naturally, the finding that brain mass increases independently in birds and mammals for a same body size with the advent of endothermy is fundamental to the understanding of how endothermy is transformative; but increased body mass should only be treated as a cause or driver of diversity where mechanistically warranted, such as through the ingestion of more energy per unit time allowed by increased body mass (Fonseca-Azevedo and Herculano-Houzel, 2012; Herculano-Houzel, 2015). Importantly, I have recently showed that body mass is not a direct predictor of life history variables such as age at sexual maturity and maximal longevity across warm-blooded amniotes; numbers of pallial neurons are, and with a much higher r^2^ of 0.74, compared to 0.44 for body mass (Herculano-Houzel, 2019). The finding that the pace of life slows down with increasing numbers of pallial neurons offers a new fundamental insight into the evolution of warm-blooded amniotes: namely, that larger numbers of pallial neurons in birds and mammals compensate for the shortening of lifespan that comes with endothermy (de Magalhães and Costa, 2009), however that happens. In this scenario, what lifespan the increased metabolic capacity of endotherms takes away, it provides back through the now affordable larger numbers of pallial neurons. It will be interesting to work out the mathematical (im)balance of this trade-off.

### The end of the “reptilian brain”

The unequivocal continuity shown here between the cousin clades of mammals and reptiles in the relationship between numbers of neurons and structure mass of the telencephalon or cerebral cortex should put to rest any lingering remnants of the incorrect, unwarranted, yet still popular notion of a “triune brain” with a “mammalian neocortex” expanding over a “reptilian brain” in evolution (MacLean, 1990; Herculano-Houzel, 2016). The pallium, whether organized in visible or in cryptic layers (Stacho et al., 2020), and regardless of how much it proliferates and expands over the rest of the brain, has the same developmental origin from the dorsal cap of the second neuromere in the neural tube across different amniotes (Puelles et al., 2013). Thus, reptiles do have a pallium, which forms as a major part of a telencephalon that I show to be composed according to the same neuronal scaling rules that are shared with mammals. Conversely, the neuronal composition of the supposed “reptilian” rest of brain is also shared between reptiles and mammals. In both cases, the main difference between mammals and reptiles seems to lie in how many neurons the developing body can afford in each brain structure. Whether the body has the increased oxidative capacity that also allows for endothermy seems to be a key factor in determining whether a modern amniote has fewer or more than 10 million neurons in its telencephalon, with all its consequences for cognitive capabilities, energetic economy, and pace of life, as I explore in the following chapter.

## Acknowledgments

Thanks to Jon Kaas for comments on this manuscript, and to all past collaborators for making the curated database possible. This work was not supported by any funding agencies.

## Notes

### Competing Interest Statement

The authors have declared no competing interest.

### Summary of Updates

Additional analyses, text edited. The revised manuscript will be resubmitted for publication.

## References

Alvarez LW, Alvarez W, Asaro F, Michel HV (1980) Extraterrestrial cause for the Cretaceous-Tertiary extinction. Science 208, 1095–1108.

Álvarez-Carretero S, Tamuri AU, Battini M, Nascimento FF, Carlisle E, Asher RJ, Yang Z, Donoghue PCJ, dos Reis M (2022) A species-level timeline of mammal evolution integrating phylogenomic data. Nature 602, 263–267.

Bennett AF, Ruben JA (1979) Endothermy and activity in vertebrates. Science 206, 649–655.

Benoit J, Fernandez V, Manger PR, Rubidge BS (2017a) Endocranial casts of pre-mammalian therapsids reveal an unexpected neurological diversity at the deep evolutionary root of mammals. Brain Behav Evol 90, 311–333.

Benoit J, Manger PR, Norton LA, Fernandez V, Rubidge BS (2017b) Synchrotron scanning reveals the palaeoneurology of the head-butting Moschops capensis (Therapsida, Dinocephalia). PeerJ 5, e3496.

Benoit J, Legendre LJ, Tabuce R, Obada T, Mararescul V, Manger P (2019) Brain evolution in Proboscidea (Mammalia, Afrotheria) across the Cenozoic. Scientific Reports 9, 9323.

Bertrand OC, Shelley SL, Wible JR, Williamson TE, Holbrook LT, Chester SGB, Butler IB, Brusatte SL (2020) Virtual endocranial and inner ear endocasts of the Paleocene “condylarth” *Chriacus*: new insight into the neurosensory system and evolution of early placental mammals. J Anat 236, 21–49.

Bertrand OC, Shelley SL, Williamson TE, Wible JR, Chester SGB, Flynn JJ, Holbrook LT, Lyson TR, Meng J, Miller IM, Püschel HP, Smith T, Spaulding M, Tseng ZJ, Brusatte SL (2022) Brawn before brains in placental mammals after the end-Cretaceous extinction. Science 376, 80–85.

Bininda-Emonds ORP, Cardillo M, Jonew KE, MacPhee RDE, Beck RMD, Grenyer R et al. (2007) The delayed rise of present-day mammals. Nature 446, 507–512.

Boonstra LD (1968) The braincase, basicranial axis and median septum in the Dinocephalia. Ann S Afr Mus 50, 195–273.

Brusatte SL (2008) The rise and fall of the dinosaurs: A new history of their lost world. New York, HarperCollins.

Brusatte SL, O’Connor JK, Jarvis ED (2015) The origin and diversification of birds. Curr Biol 25, R888–898.

Burish MJ, Peebles JK, Tavares L, Baldwin M, Kaas JH, Herculano-Houzel S (2010) Cellular scaling rules for primate spinal cords. Brain, Behavior and Evolution 76, 45–59.

Cameron J, Shelley SL, Williamson TE, Brusatte SL (2018) The brain and inner ear of the early Paleocene “Condylarth*” Carsioptychus coarctatus*: Implications for early placental mammal neurosensory biology and behavior. Anat Rec 302, 306–324.

Castanhinha R, Araújo R, Júnior LC, Angielczyk KD, Martins GG, Martins RM, Chaouiya C, Beckmann F, Wilde F (2013) Bringing dicynodonts back to life: paleobiology and anatomy of a new emydopoid genus from the Upper Permian of Mozambique. PLoS One 8, e80974.

Chen C-K, Chuang H-F, Wu S-M, Li W-H (2019) Feather evolution from precocial to altricial birds. Zool Studies 58, 24.

Clark DA, Mitra PP, Wang SS (2001) Scalable architecture in mammalian brains. Nature 411: 189–193.

De Magalhães JP, Costa J (2009) A database of vertebrate longevity records and their relation to other life-history traits. J Evol Biol 22, 1770–1774.

Dos Santos SE, Porfirio J, Da Cunha FB, Manger PR, Tavares W, Pessoa L, Raghanti MA, Sherwood CC, Herculano-Houzel S (2017) Cellular scaling rules for the brain of marsupials: not as “primitive” as expected. Brain Behav Evol 89, 48–63.

Dozo MT, Martínez G (2016) First digital cranial endocasts of late Oligocene Notohippidae (Notoungulata): Implications for endemic South American ungulates brain evolution. J Mammal Evol 23, 1–16.

Fonseca-Azevedo K, Herculano-Houzel S (2012) Metabolic constraint imposes tradeoff between body size and number of brain neurons in human evolution. Proc Natl Acad Sci USA 109, 18571–18576.

Goldowitz D, Hamre K (1998) The cells and molecules that make a cerebellum. Trends Neurosci 21, 375–382.

Gutiérrez-Ibáñez C, Iwaniuk AN, Wylie DR (2018) Parrots have evolved a primate-like telencephalic-midbrain-cerebellar circuit. Sci Rep 8, 9960.

Herculano-Houzel S (2010) Coordinated scaling of cortical and cerebellar numbers of neurons. Frontiers in Neuroanatomy 4, 12.

Herculano-Houzel S (2015) Decreasing sleep requirement with increasing numbers of neurons as a driver for bigger brains and bodies in mammalian evolution. Proc Biol Sci 282, 20151853.

Herculano-Houzel S (2016) The human advantage: a new understanding of how our brains became remarkable. CaM_br_idge, MIT Press.

Herculano-Houzel S (2019) Longevity and sexual maturity vary across species with number of cortical neurons, and humans are no exception. J Comp Neurol 527, 1689–1705.

Herculano-Houzel S (2023). Theropod dinosaurs had primate-like numbers of telencephalic neurons. J Comp Neurol 531, 962–974.

Herculano-Houzel S (2025). A supply-limited framework of brain function accounts for the evolution of endothermic brains. Accompanying chapter.

Herculano-Houzel S, Lent R (2005) Isotropic fractionator: a simple, rapid method for the quantification of total cell and neuron numbers in the brain. J Neurosci 25, 2518–2521.

Herculano-Houzel S, Dos Santos S (2018) You do not mess with the glia. Neuroglia 1, 193–219.

Herculano-Houzel S, Rothman DL (2022) From a demand-based to a supply-limited framework of brain metabolism. Front Integr Neurosci 16, 818685.

Herculano-Houzel S, Mota B, Lent R (2006) Cellular scaling rules for rodent brains. Proceedings of the National Academy of Sciences USA 103, 12138–12143.

Herculano-Houzel S, Collins C, Wong P, Kaas JH (2007) Cellular scaling rules for primate brains. Proceedings of the National Academy of Sciences USA 104, 3562–3567.

Herculano-Houzel S, Manger PR, Kaas JH (2014a) Brain scaling in mammalian brain evolution as a consequence of concerted and mosaic changes in numbers of neurons and average neuronal cell size. Front Neuroanat 8, 77.

Herculano-Houzel S, Avelino-de-Souza K, Neves K, Porfírio J, Messeder D, Calazans I, Mattos L, Maldonado J, Manger PM (2014b) The elephant brain in numbers. Front Neuroanat 8, 46.

Herculano-Houzel S, Kaas JH, Miller D, Von Bartheld CS (2015a) How to count cells: the advantages and disadvantages of the isotropic fractionator compared with stereology. Cell Tissue Research 360, 29–42.

Herculano-Houzel S, Catania K, Manger PR, Kaas JH (2015b) Mammalian brains are made of these: A dataset on the numbers and densities of neuronal and non-neuronal cells in the brain of glires, primates, scandentia, eulipotyphlans, afrotherians and artiodactyls, and their relationship with body mass. Brain Behav Evol 86, 145–163

Herculano-Houzel S, Messeder D, Fonseca-Azevedo K, Araujo Pantoja N (2015c) When larger brains do not have more neurons: Intraspecific increase in numbers of cells is compensated by decreased average cell size. Front Neuroanat 9, 64.

Herculano-Houzel S, Cunha F, Reed JL, Kaswera-Kyamakya C, Gillissen E, Manger PR (2020) Microchiropterans have a diminutive cerebral cortex, not an enlarged cerebellum, compared to megachiropterans and other mammals. Journal of Comparative Neurology 528, 2978–2993.

Herrup K (1986) Cell lineage relationships in the development of the mammalian CNS: role of cell lineage in control of cerebellar Purkinje cell number. Dev Biol 115, 148–154.

Hillman SS, Hancock TV, Hedrick MS (2013) A comparative meta-analysis of maximal aerobic metabolism of vertebrates: implications for respiratory and cardiovascular limits to gas exchange. J Comp Physiol 183, 167–179.

Hopson JA (1979) Paleoneurology. In Glans C, Northcutt RG, Ulinski P (eds), Biology of the Reptilia. NY, Academic Press, pp 39–146.

Jardim-Messeder D, Lambert K, Noctor S, Marques Pestana F, DeCastro Leal ME, Bertelsen MF, Alagaili AN, Mohammad OB, Manger PR, Herculano-Houzel S (2017) Dogs have the most neurons, though not the largest brain: Trade-off between body mass and number of neurons in the cerebral cortex of large carnivoran species. Front Neuroanat 11, 118.

Jerison HJ (1973) Evolution of the brain and intelligence. New York, Academic Press.

Kabadayi C, Taylor LA, von Bayern AMP, Osvath M (2016) Ravens, New Caledonian crows and jackdaws parallel great apes in motor self-regulation despite smaller brains. Roy Soc Open Sci 3, 160104.

Kemp TS (2006) The origin and early radiation of the therapsid mammal-like reptiles: a palaeobiological hypothesis. J Evol Biol 19, 1231–1247.

Kielan-Jaworowska Z, Cifelli RL, Luo ZX (2004) Mammals from the age of dinosaurs: Origins, evolution, and structure. NY, Columbia University Press.

Ksepka DT, Balanoff AM, Smith NA, Bever GS, Bhullar B-AS, Bourdon E et al. (2020) Tempo and pattern of avian brain size evolution. Curr Biol 30, 2026–2036.

Kullman A, Trivedi N, Howell D et al. (2020) Oxygen tension and the VHL-Hif1a pathway determine onset of neuronal polarization and cerebellar germinal zone exit. Neuron 106, 607–623.

Kverkova K, Marhounová L, Polonyiová A, Kocourek M, Zhang Y, Olkowicz S, Stratková B, Pavelková Z, Vodicka R, Frynta D, Nemec P (2022) The evolution of brain neuron numbers in amniotes. Proc Natl Acad Sci USA 119, e2121624119.

Laas M (2015) Virtual reconstruction and description of the cranial endocast of *Pristerodon mackayi* (Therapsida, Anomodontia). J Morphol 276, 1089–1099.

Laas M, Kaestner A (2017) Evidence for convergent evolution of a structure comparable to the mammalian neocortex in a Late Permian therapsid. J Morphol 278, 1033–1057.

Luo ZX, Crompton AW, Sun AL (2001) A new mammaliaform from the Early Jurassic and evolution of mammalian characteristics. Science 292, 1535–1540.

MacLean PD (1990) The triune brain in evolution. New York, Plenum.

Macrini TE (2006) The evolution of endocranial space in mammals and non-mammalian cynodonts. PhD Thesis, University of Texas at Austin.

Macrini TE, Rougier GW, Rowe T (2007) Description of a cranial endocast from the fossil mammal *Vincelestes neuquenianus* (Theriiformes) and its relevance to the evolution of endocranial characters in Therians. Anat Rec 290, 875–892.

Maugoust J, Orliac MJ (2021) Endocranial cast anatomy of the extinct Hipposiderid bats *Pallaeophyllophora* and *Hipposideros* (*Pseudorhinolophus*) (Mammalia: Chiroptera). J Mammal Evol 28, 679–706.

Mota B, Herculano-Houzel S (2014) All brains are made of this: a fundamental building block of brain matter with matching neuronal and glial masses. Front Neuroanat 8, 127.

Nespolo RF, Bacigalupe LD, Figueroa CC, Koteja P, Opazo JC (2011) Using new tools to solve an old problem: the evolution of endothermy in vertebrates. Trends Ecol Evol 26, 414–423.

Newham E, Gill PG, Corfe IJ (2022) New tools suggest a middle Jurassic origin for mammalian endothermy. BioEssays 44, 2100060.

Ngwenya A, Patzke N, Manger PR, Herculano-Houzel S (2016) Continued growth of the central nervous system without mandatory addition of neurons in the Nile crocodile (*Crocodylus niloticus*). Brain Behav Evol 87, 19–38.

Norell MA, Xu X (2005) Feathered dinosaurs. Annu Rev Earth Planet Sci 33, 277–299.

Olkowicz S, Kocourek M, Lucan RK, Portes M, Fitch WT, Herculano-Houzel S, Nemec P (2016) Birds have primate-like numbers of neurons in the telencephalon. Proc Natl Acad Sci USA 113, 7255–7260.

Puelles L, Harrison M, Paxinos G, Watson C (2013) A developmental ontology for the mammalian brain based on the prosomeric model. Trends Neurosci 36, 570–578.

Pukart L, Tuff JM, Shah M, Kaufmann LV, Altringer C, Maier E, Schneeweiss U, Tunckol E., Eigen L, Holtze S, Fritsch G, Hildebrandt T, Brecht M (2022) Trigeminal ganglion and sensory nerves suggest tactile specialization of elephants. Curr Biol 32, 904–910.

Pusch LC, Kammerer CF, Fröbisch J (2019) Cranial anatomy of the early cynodont *Galesaurus planiceps* and the origin of mammalian endocranial characters. J Anat 234, 592–621.

Quiroga JC (1980) The brain of the mammal-like reptile *Probainognathus jenseni* (Therapsida, Cynodontia): a correlative paleo-neoneurological approach to the neocortex at the reptile-mammal transition. J Hirnforsch 21, 299–336.

Quiroga JC (1984) The endocranial cast of the advanced mammal-like reptile *Therioherpeton cargnini* (Therapsida-Cynodontia) from the Middle Triassic of Brazil. J Hirnforsch 25, 285–290.

Ramnani N (2006) The primate cortico-cerebellar system: anatomy and function. Nat Rev Neurosci 7, 511–522.

Rodrigues PG, Ruf I, Schultz CL (2014) Study of a digital cranial endocast of the non-mammaliaform cynodont *Brasilitherium riograndensis* (Later Triassic, Brazil) and its relevance to the evolution of the mammalian brain. Pällaontol Z 88, 329–352.

Rowe TB, Macrini TE, Luo Z-X (2011) Fossil evidence on origin of the mammalian brain. Science 332, 955–957.

Sarko DK, Catania KC, Leitch DB, Kaas JH, Herculano-Houzel S (2009) Cellular scaling rules of insectivore brains. Frontiers in Neuroanatomy 3, 8.

Sigurdsen T, Huttenlocher AK, Modesto SP, Rowe TB, Damiani R (2012) Reassessment of the morphology and paleobiology of the therocephalian *Tetracynodon darti* (Therapsida), and the phylogenetic relationships of Bauroidea. J Vert Paleontol 32, 1113–1134.

Sol D, Olkowicz S, Sayol F, Kocourek M, Zhang Y, Marhounová L, Osadnik C, Corssmit E, Garcia-Porta J, Martin TE, Lefebvre L, Nemec P (2022) Neuron numbers link innovativeness with both absolute and relative brain size in birds. Nature Ecol Evol, 10.1038/s41559-022-01815-x.

Stacho M, Herold C, Rook N, Wagner H, Axer M, Amunts K, Güntürkün O (2020) A cortex-like canonical circuit in the avian forebrain. Science 369, eabc5534.

Ströckens F, Neves K, Kirchem S, Schwab C, Herculano-Houzel S, Güntürkün O (2022) High associative neuron numbers could drive cognitive performance in corvid species. J Comp Neurol 530, 1588–1605.

Sultan F (2002). Analysis of mammalian brain architecture. Nature 415, 133–134.

Ventura-Antunes L, Dasgupta OM, Herculano-Houzel S (2022) Resting rates of blood flow and glucose use per neuron are proportional to number of endothelial cells available per neuron across sites in the rat brain. Front Integr Neurosci 16, 821850.

